# Reference-free and robust copy number inference in single-cell tumor transcriptomes

**DOI:** 10.64898/2026.08.03.742406

**Authors:** M. Mert Gençtürk, A. Ercüment Çiçek

## Abstract

Copy number changes are a hallmark of cancer and reveal which cells are malignant and how tumors evolve. Single-cell RNA sequencing is widely used to infer them in individual tumor cells. Existing methods do so by normalizing each cell against a reference of normal cells. As a result, their output depends on which cells are analyzed together, they often label normal cells as aneuploid, and they struggle in samples without normal cells. Here we present REFCON, a deep-learning model that infers copy number from each cell’s own expression, without a reference. We evaluated REFCON on more than 410,000 held-out cells from cell lines, organoids and 112 tumors of nine cancer types, profiled with seven sequencing technologies. It agreed more closely with single-cell DNA than existing methods, was unaffected by sample composition, and labeled far fewer normal cells as aneuploid. In patient-derived organoids, which contain no normal cells, it gave consistent profiles across passages and tracked clonal evolution. REFCON addresses long-standing limitations of reference-based inference and enables robust, reference-free copy number calling from single-cell transcriptomes.

## 1 Introduction

Copy number alterations, gains and losses of genomic regions, are prevalent across human cancers and contribute to cancer initiation, progression, and therapeutic resistance [1]. Chromosome copy number (CN) changes can function as point-mutation-independent sources of drug resistance, generating intratumoral heterogeneity in the form of gene dosage alterations upon which selective pressures can act [2]. Copy number has been charted largely in bulk, that is, from an entire tumor pooled and profiled as one sample; pan-cancer surveys of thousands of such samples link copy number signatures to therapeutic response [3]. A bulk profile, however, averages over the millions of cells in a sample, whereas a tumor is a mosaic of genetically distinct subclones, and the copy number differences between these subclones encode the tumor’s clonal architecture and evolution [4, 5]. For instance, in early-stage non-small-cell lung cancer, tumors with a high proportion of subclonal copy number alterations carry a nearly fivefold higher risk of recurrence or death, independent of other prognostic factors [6]. At the single-cell level, these alterations directly shape cellular phenotypes: in glioblastomas, the relative frequency of cells in each transcriptional state is influenced by copy number amplifications of the *CDK4*, *EGFR*, and *PDGFRA* genes, each of which favors a distinct transcriptional state [7]. Yet, how cell-to-cell copy number alterations drive genomic and phenotypic variation in tumors remains understudied [8], in part because resolving copy number profiles at cellular resolution remains technically challenging.

Due to the limited accessibility of high-throughput single-cell DNA sequencing platforms [9, 10], and the early-stage development of paired multi-omics assays such as DNTR-seq and scONE-seq [11–13], single-cell copy number is predominantly inferred from scRNA-seq data, which is instead captured routinely and at scale. This inference has become the standard foundational step in single-cell oncology, used to distinguish malignant from non-malignant cells and to delineate subclonal architecture [14, 15]. Both downstream processes strongly depend on this step.

The inference of copy number profiles from scRNA-seq data remains insufficiently robust to be treated as a quantitative measurement, primarily because transcriptomic abundance serves as an imperfect proxy for genomic dosage. While elevated copy numbers generally upregulate expression and deletions downregulate it, the transcriptome is simultaneously confounded by complex transcriptional regulation, dynamic cell states, and technical noise [16]. State-of-the-art computational methods uniformly address this challenge through a shared paradigm: Normalizing tumor gene expression against a baseline derived from non-malignant reference cells. For instance, inferCNV, the foundational approach in the field, relies on user-annotated normal cells [14, 17]. CopyKAT utilizes an explicit reference when provided, or alternatively, implements a clustering heuristic to identify normal subpopulations within the sample [18]. Similarly, SCEVAN and CopyVAE detect intra-sample normal cells [19, 20]. In their absence, SCEVAN can also synthesize an artificial baseline. This shared dependence on a reference is the field’s structural weakness. First, the results are neither robust nor consistent: an independent benchmark of 21 datasets found large variance across datasets, strong sensitivity to which cells are used as the reference, and no method clearly superior to the others [16]. Across methods, calls on the same tumor can overlap by less than 30% and include numerous false positives [21], and when run on healthy cells, they report spurious copy number changes unless the reference cells are of the same type and from the same sample [16]. Second, the reference is often unavailable: Pure or highly homogeneous tumors, cancer cell lines [22], and tumor organoids [23] contain no normal cells and too little heterogeneity to synthesize a baseline, and the external reference cells used as an alternative may lower accuracy even when the cell or tissue type matches [16]. Reference-free copy number inference from scRNA-seq remains an open problem in single-cell data science.

Here, we introduce a novel, reference-free design paradigm that entirely circumvents this structural pitfall. Our method, REFCON, reformulates copy number inference as a local problem: it predicts copy number within windows of neighboring genes and assembles these local predictions into a genome-wide profile, without in-sample normal cells, a synthesized baseline or an external panel. Trained once on cell lines with bulk-DNA copy number, it is significantly more accurate than existing methods against bulk and paired single-cell DNA, on unseen cell lines and tumors and across seven sequencing platforms. It thereby makes samples without normal cells, such as cell lines and organoids, analyzable on their own. Its prediction for a cell does not depend on which other cells are analyzed with it, which removes cohort composition as a confounder of copy number inference. In tumor organoids, its profiles agreed within lines significantly more than those of existing methods, remained stable when only the expression state changed, and recovered without a reference the clonal turnover that previously required one. It classifies cells as aneuploid or diploid as accurately as the best existing method while labeling far fewer normal cells aneuploid. Where matched normal cells exist, REFCON can use them to refine its predictions, turning a requirement of previous methods into an optional enhancement. By enabling truly reference-free per-cell copy number profiling, REFCON turns an inference the field already depends on into a robust estimate, applicable across the tumor transcriptomes already generated at scale.

## 2 Results

### 2.1 Model and evaluation overview

REFCON is a relatively small, specialized transformer-based model [24], with about one million parameters, that predicts a genome-wide copy number profile for a single cell from its scRNA-seq expression alone, without a reference panel, a diploid baseline or a group of normal cells (Figure 1). It was trained on 60,179 cells from 140 cell lines with bulk-DNA copy number, together with 4,200 healthy cells from Tabula Sapiens [25, 26] labeled diploid (Methods, Section 4.1).

**Fig. 1.**
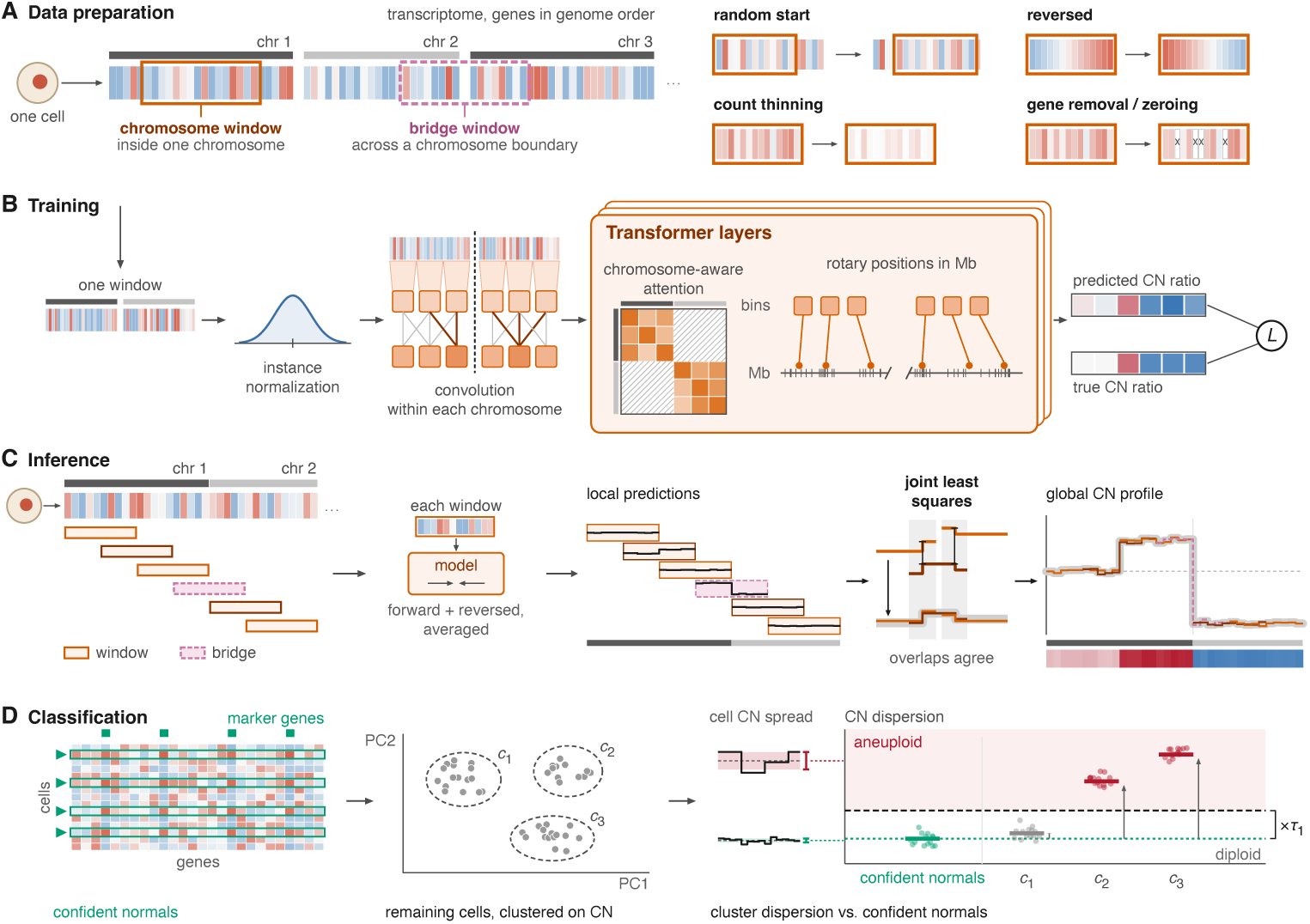
Overview of REFCON. **A** Training windows. Each training cell’s expression profile is read with its genes in genomic order. A window covers consecutive genes of one chromosome (*chromosome window* ) or spans the boundary between two adjacent chromosomes (*bridge window* ). During training, each window starts at a random gene offset and may be presented in reversed gene order, have its counts thinned, or have genes removed or set to zero. Its copy number targets stay unchanged. **B** Model. Each window is log-transformed, scaled by instance normalization and grouped into bins of eight consecutive genes. Convolutions and transformer layers never connect bins of different chromosomes, with rotary positions that encode the genomic distance between bins in megabases. The model predicts one copy number ratio per bin, relative to the window mean, and is trained against the true ratios with the loss *L*. True ratios come from bulk DNA copy number for cell lines and are flat for healthy cells. **C** Reference-free inference. A new cell’s genome is tiled with overlapping chromosome windows and bridge windows. Each window is predicted in its original and in reversed gene order, and the two predictions are averaged. Each local prediction is known only up to a scale. A joint least-squares fit, in which overlapping windows must agree, stitches them into one genome-wide copy number profile per cell, normalized to a mean of one. **D** Tumor classification in mixed cohorts. A marker gene test on gene expression identifies *confident-normal* cells. The remaining cells are clustered on their predicted copy number profiles. A cluster is called aneuploid when its median per-cell copy number dispersion is at least *τ*_1_ times that of the confident-normal cells, and diploid otherwise. Samples with fewer than two confident-normal cells receive one sample-level call (Methods).

REFCON treats copy number inference as a local problem. It reads a cell’s expression in windows of up to 2,048 consecutive genes of one chromosome in genomic order, groups the genes into bins of eight and predicts each bin’s copy number relative to the window mean, exploiting the tendency of neighboring genes to share copy number state [14]. The design aims at a single trained model that applies to any scRNA-seq dataset without retraining. Predicting copy number relative to a local window makes REFCON independent of the genes outside it. REFCON uses no gene identifiers, chromosome labels or absolute genomic coordinates, only the order of the genes and the genomic distances between them, so it does not depend on which genes a panel measures. It normalizes each cell’s input, which removes differences in sequencing depth and platform scale. It is trained on thinned counts and removed genes, which mimic shallower sequencing and sparser gene panels (Figure 1A,B). For a new cell, overlapping windows tile the genome. Bridge windows span the boundary between two adjacent chromosomes and place them on a common scale. Within a bridge, no attention or convolution connects the bins of the two chromosomes, as copy number segments do not cross chromosome boundaries. A joint fit stitches all windows into one genome-wide profile, normalized to a mean of one (Figure 1C). Like other expression-based callers, REFCON therefore estimates relative copy number dosage rather than integer copy number or absolute ploidy. All reported results use an ensemble of two models trained with different random seeds (Methods, Sections 4.3 and 4.4).

Reference-free per-cell copy number inference is the core contribution of REFCON. We also use the inferred profiles to classify cells as aneuploid or diploid, a separate, within-sample problem that requires a comparison with the diploid cells of the same sample. For this, confident-normal cells are first identified from gene expression with a marker-gene test [19]. The remaining cells are clustered on their predicted profiles [18], and each cluster is labeled by comparing the spread of its cells’ predicted profiles across their genes, which we call copy number dispersion, with that of the confident-normal cells (Figure 1D; Methods, Sections 4.5–4.7).

We evaluated REFCON on more than 410,000 cells not used in training, from held-out cell lines, 112 tumors, 48 organoid samples and 19 normal samples, profiled on seven platforms: 10x Chromium, BD Rhapsody, Seq-Well S^3^, Smart-seq2, MARS-seq, DNTR-seq and scONE-seq (Table 1). We compared it with the expression-based copy number callers SCEVAN [19], CopyKAT [18], CopyVAE [20] and inferCNV [14, 17] on four fronts: (i) reference-free copy number accuracy on held-out cell lines and a patient tumor with bulk and paired single-cell DNA (Section 2.2); (ii) independence of the predictions from cohort composition (Section 2.3); (iii) following the genomes of patient-derived organoids through culture and treatment (Section 2.4); and (iv) aneuploid versus diploid classification in tumors, the normal cells inside them and fully normal samples (Section 2.5). Unless otherwise stated, REFCON’s profiles were inferred without a reference and without any retraining or per-platform tuning.

**Table 1.** Datasets used for training, validation and testing, grouped by analysis.

| Dataset | Platform | Units | Cells | Ground truth |
| --- | --- | --- | --- | --- |
| *Training* |  |  |  |  |
| MCF7 [37] | 10x | 1 line | 6,749 | bulk |
| COLO320 [41] | 10x Multiome | 2 sublines | 9,000 | bulk |
| MDA-MB-231 [42] | wellDR-seq | 1 line | 3,834 | bulk |
| Andor gastric [43] | 10x | 3 lines | 9,951 | bulk |
| CCLE pan-cancer [22] | 10x | 135 lines | 30,645 | bulk |
| Tabula Sapiens [25, 26] | 10x | 14 organs, 6 donors | 4,200 | diploid |
| *Validation and development* |  |  |  |  |
| CCLE held-out [22] | 10x | 7 lines | 2,255 | bulk |
| Tabula Sapiens [25, 26] | 10x | 4 organs, 2 donors | 1,200 | diploid |
| Tirosh melanoma [17] | Smart-seq2 | 3 patients | 1,664 | annotation |
| *Test: copy number (Section 2.2)* |  |  |  |  |
| CCLE esophageal [22] | 10x | 7 lines | 2,543 | bulk |
| CCLE mixed tissue [22] | 10x | 3 lines | 1,512 | bulk |
| MIX-seq pan-cancer [27] | 10x | 49 lines | 3,055 | bulk |
| HCT116 [11] | DNTR-seq | 1 line | 1,467 | scWGS |
| A375 [11] | DNTR-seq | 1 line | 64 | scWGS |
| Astrocytoma [12] | scONE-seq | 1 tumor | 840 | scWGS |
| Colonic epithelium [29] | Fluidigm C1 | 96 libraries | 96 | normal |
| Epidermis [30] | Smart-seq2 | 200 cells | 200 | normal |
| *Test: cohort composition (Section 2.3)* |  |  |  |  |
| CellBench lung [31] | 10x | 4 lines | 3,078 | identity |
| CCLE esophageal [22] | 10x | 7 lines | 2,543 | identity, bulk |
| MIX-seq lung [27] | 10x | 21 lines | 1,268 | identity, bulk |
| MIX-seq pan-cancer [27] | 10x | 73 lines | 4,707 | identity |
| *Test: organoids (Section 2.4)* |  |  |  |  |
| Pancreatic organoids [33] | Seq-Well S<sup>3</sup> | 48 samples, 14 lines | 36,752 | — |
| Pancreatic biopsies [33] | Seq-Well S<sup>3</sup> | 22 biopsies | 7,875 | — |
| *Test: classification (Section 2.5)* |  |  |  |  |
| Atlas tumors [15, 34] | various | 94 tumors, 8 studies | 139,606 | annotation |
| Tirosh melanoma [17] | Smart-seq2 | 12 patients | 2,285 | annotation |
| Wang colorectal [35] | BD Rhapsody | 5 patients | 194,815 | annotation |
| PBMC 3K [36] | 10x | 1 donor | 2,700 | normal |
| Tabula Sapiens [25, 26] | 10x | 18 organs, 11 donors | 14,400 | normal |

Units: the lines, tumors, patients, samples or organs of a dataset. Colonic epithelium and Epidermis: normal cells supplied as the reference to the other methods on HCT116 and A375. Cells: the cells used in training or validation, or scored in a test. Astrocytoma: 390 malignant and 450 non-malignant cells. Bulk: DepMap 24Q4 gene-level copy number of the line. scWGS: paired single-cell whole-genome sequencing of the same cells. Annotation: the authors' malignant and non-malignant cell annotations. Normal: samples without malignant cells. Diploid: labeled diploid by construction. Identity: the known cell line of each cell. —: no ground truth (Section 2.4).

### 2.2 REFCON infers copy number without a reference more accurately than existing callers

We compare REFCON with four state-of-the-art reference-based CN callers across held-out cell lines and on multiple sequencing platforms. We used DNA-derived copy number profiles as ground truth. Because the methods output copy number on different scales, we use metrics that need no cutoff on the predictions: Pearson and Spearman correlation, and AUROC and AUPRC for loss and gain. Loss and gain thresholds apply to the ground truth only (Methods, Section 4.9).

#### Per cell line benchmark on held-out cell lines

We first evaluated REFCON against bulk-DNA copy-number profiles for 59 cell line profiles (57 distinct lines) held out from training: seven esophageal lines from a tissue absent from the training panel, three additional mixed-tissue lines from the CCLE panel [22], and 49 MIX-seq lines [27] (Methods, Section 4.1). We evaluated each cohort under two settings. In the *per cell line* setting, each line was analyzed separately, precluding construction of an internal reference. In the *pooled* setting, all lines within a cohort were analyzed together, allowing reference-based methods to estimate an internal baseline from between-line heterogeneity. REFCON was run on each line alone. In the per cell line setting, REFCON had the highest AUROC for copy-number gains and losses and the highest correlation with the bulk-DNA ground truth in every cohort. It ranked first in Pearson correlation for all 59 cell line profiles, ahead of the next-best method, SCEVAN. Across the 59 profiles, its correlation and its loss and gain AUROC were significantly higher than those of every other method. The remaining methods generally showed little correlation with the ground truth (Figure 2A). Pooling raised most scores of the reference-based methods, yet REFCON’s reference-free predictions matched or exceeded them on most cohorts and metrics and stayed competitive on the rest. Pooled inferCNV scored at most 0.03 higher in two of the 18 cohort and metric pairs, the gain AUROC and gain AUPRC of the mixed-tissue lines. Supplementary Table 1 gives every metric of each method per cohort, per cell line and pooled. Supplementary Table 2 gives the paired tests of REFCON against each method, over cell lines and, for the cohorts with paired single-cell DNA, over cells.

**Fig. 2.**
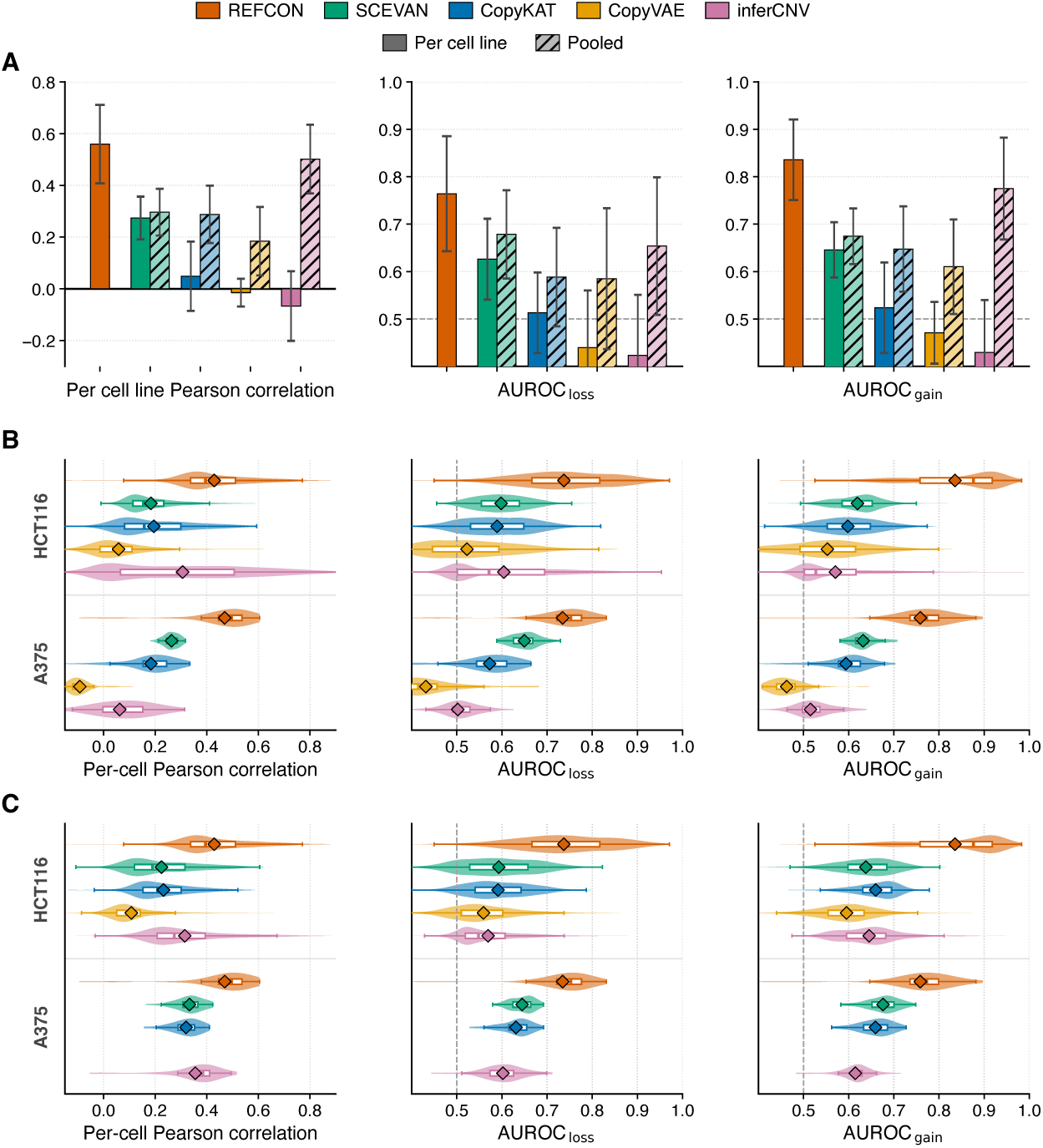
Copy number inference benchmark on held-out cell lines. Methods compared: REFCON, SCEVAN, CopyKAT, CopyVAE and inferCNV. **A** Per cell line Pearson correlation and loss and gain AUROC against bulk DNA copy number on 59 held-out cell line profiles (57 distinct lines): 7 esophageal and 3 mixed-tissue lines from the Cancer Cell Line Encyclopedia (CCLE) and 49 MIX-seq lines. Each metric is computed on the pseudobulk profile of a line. Bars are the mean across lines, and error bars the standard deviation. REFCON was run on each cell line alone with-out a reference. The other methods were run on each line alone (per cell line) and on all lines of a cohort together (pooled). **B** Per-cell Pearson correlation and loss and gain AUROC against paired single-cell DNA (DNTR-seq) on HCT116 (1,467 cells) and A375 (64 cells). All methods ran without a reference. **C** Same as **B**, with lineage-matched normal cells supplied as the reference to the other methods (colonic epithelium for HCT116, epidermis for A375). REFCON ran without a reference as in **B**. CopyVAE gave no output on A375 with the reference cells.

#### Per-cell reference-free benchmark on held-out cell lines

We next evaluated REFCON at single-cell resolution against paired single-cell DNA and on scRNA-seq protocols outside its predominantly 10x Chromium [28] training corpus (Table 1). The benchmark comprised two DNTR-seq [11] cell lines, HCT116 (colorectal; 1,467 cells) and A375 (melanoma; 64 cells), each with per-cell copy-number ground truth from paired single-cell DNA (Methods, Section 4.1). Figure 2B shows the per-cell results without reference cells. On both lines, REFCON had the highest correlation with the per-cell DNA-derived profiles, with Pearson correlations of 0.43 on HCT116 and 0.47 on A375, against 0.31 and 0.26 for the next-best expression-based method. It also detected losses and gains best, with loss and gain AUROC of 0.74 and 0.84 on HCT116 and 0.73 and 0.76 on A375. Its correlation and its loss and gain detection were significantly higher than those of the next-best method on each line. We additionally examined how reliable each method’s predictions are when a line is analyzed alone, by counting the gains and losses called on copy-neutral genes. On these two lines and the 49 MIX-seq lines, REFCON called the fewest, except on HCT116, where inferCNV called slightly fewer. Supplementary Table 3 gives these false calls per cohort and method. Supplementary Table 4 gives every per-cell metric on the three cohorts with paired single-cell DNA, with the class prevalences, for each method without and with a reference, and for the allele-based caller Numbat, which uses sequencing reads.

#### Per-cell benchmark with reference cells

Because the HCT116 and A375 cell line datasets contain no normal cells from which reference-based methods can estimate a baseline, we followed Schmid et al. [16] and supplied each competing method with a lineage-matched pseudo-reference panel: 96 colonic epithelial libraries from Kinchen et al. [29] for HCT116 and 200 normal epidermal cells, mostly melanocytes, from Belote et al. [30] for A375. Figure 2C shows that, even when the competing methods were supplied with these external references, REFCON’s reference-free predictions remained significantly higher in correlation and in loss and gain detection on both cell lines.

#### Per-cell benchmark on a tumor with rare clones

We also evaluated REFCON on a second-recurrence IDH-mutant astrocytoma tumor profiled by scONE-seq [12], a paired single-cell DNA and RNA assay. Among the 840 cells with both profiles that pass quality control, single-cell whole-genome sequencing of the same cells resolves 390 malignant cells into a main clone and two rare clones, clone 1 (15 cells) and clone 2 (17 cells), alongside 450 non-malignant cells. Figure 3A shows the per-cell results by clone with every method run on the malignant cells alone. REFCON had the highest correlation with the paired DNA and the best loss and gain detection in the main clone and in both rare clones. Its per-cell Pearson correlations were 0.36, 0.34 and 0.42, against 0.23, 0.30 and 0.29 for the next-best method. Over all malignant cells, its correlation and its loss and gain detection were significantly higher than those of the next-best method.

**Fig. 3.**
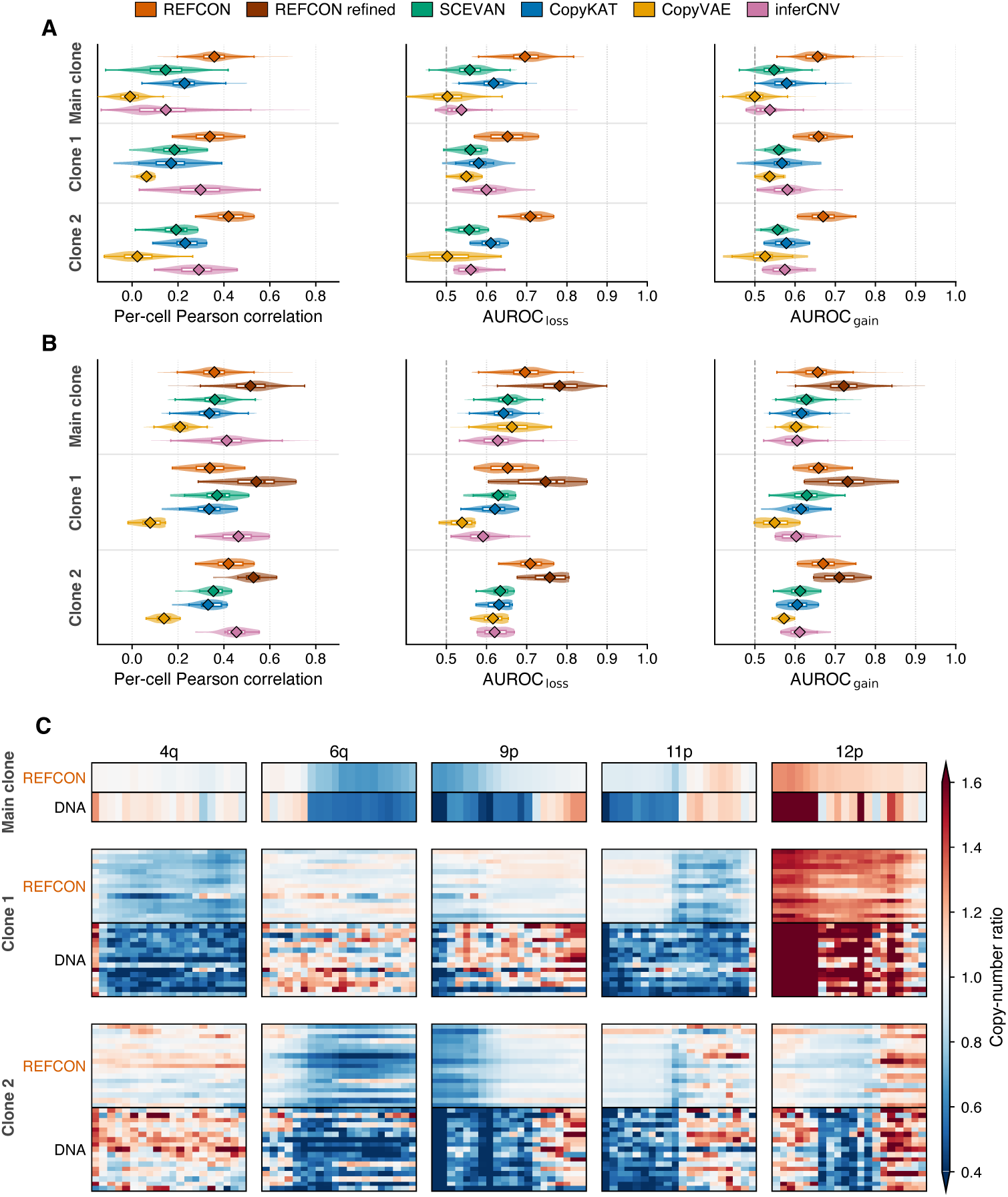
Per-cell copy number inference on a tumor with rare clones. scONE-seq astrocytoma with paired single-cell whole-genome sequencing (scWGS) of the same cells: 390 malignant cells in a main clone (358 cells) and two rare clones (clones 1 and 2, 15 and 17 cells), alongside 450 non-malignant cells. **A** Per-cell Pearson correlation and loss and gain AUROC by clone, every method run on the malignant cells alone. **B** As **A**, with the 450 non-malignant cells as the reference to every other method. REFCON is shown without a reference and with its predictions refined against its predictions for the same cells. **C** Per-cell copy number ratios of REFCON (refined, as in **B**) and scWGS on five chromosome arms that distinguish clone 1 from the main clone. 12p also distinguishes clone 2. The main clone is shown as its consensus and the rare clones per cell.

Figure 3B compares REFCON’s predictions with those of the other methods given the 450 non-malignant cells as the reference. Without a reference, REFCON still detected losses and gains marginally but significantly better than every method given these cells. Its correlation with the paired DNA was similar to SCEVAN’s (difference below 0.01) and lower than inferCNV’s. In clone 2, it detected losses and gains with AUROC of 0.71 and 0.67, against 0.63 and 0.61 for the next-best method given these cells, and its correlation was second only to inferCNV’s. To show that REFCON can also use matched normal cells from the same sample, we applied a refinement step. It is a simple SVD-based denoising of the malignant cells’ reference-free profiles against the profiles of the normal cells (Methods, Section 4.8). Figure 3B also shows the refined predictions. With them, REFCON outperformed every compared method in correlation and in loss and gain detection in the main clone and in both rare clones. Its per-cell Pearson correlations were 0.52, 0.54 and 0.53, against 0.41, 0.46 and 0.45 for the next-best method, and its gains were significant over all malignant cells. Figure 3C shows REFCON’s refined profiles and the paired DNA on the five arms that set clone 1 apart. Clone 1 retains 6q and 9p, including the *CDKN2A* locus, which are lost in the main clone. It carries a private loss of 4q and a more extensive 11p loss and 12p gain than the main clone. Clone 2 differs from the main clone by an 8q gain, a 12p loss and a 19p loss. REFCON recovered the direction of all eight events from RNA, also without a reference. On a UMAP of its profiles, both rare clones lay apart from the main clone. Supplementary Figure 1 shows the UMAP of every method. REFCON thus enables copy number inference from expression alone, without any reference. It makes cell lines analyzable on their own and outperforms existing methods on them against bulk DNA. Against single-cell DNA of the same cells, it is more accurate than existing methods in two cell lines and a tumor, including the tumor’s rare clonal events. When normal cells from the same sample are available, refinement of its predictions against these cells makes them significantly more accurate than those of every method given the same cells.

### 2.3 REFCON decouples copy number inference from cohort composition

In practice, the same cells can be analyzed alone, pooled with other samples, or alongside a different mix of cells. We call the cells analyzed together a cohort. Reference-based callers draw their baseline from the cohort, so their profile of a cell can change with it [16, 21] (Section 2.2, pooled setting). To show that REFCON’s design decouples its performance from cohort composition, we first tested whether inferred copy-number profiles recover known population identity, both when each population is analyzed independently and when all populations in a cohort are analyzed together. We then measured how the inferred copy-number profiles change as the composition of the cohort varies. We used four held-out cohorts spanning 4 to 73 cell lines: four lung cell lines from CellBench [31], seven esophageal cell lines from CCLE [22], a tissue absent from training, and two MIX-seq panels [27], 21 lung cell lines and 73 cell lines from various cancer types, which we call the pan-cancer cohort.

#### Population separation across cohort compositions

We clustered each method’s predicted profiles into the known cell line identities and scored the result with the adjusted Rand index (ARI). Figure 4A shows the ARI of each method with each cohort pooled and with each line analyzed alone. Pooled, the reference-based methods separated the lines, with ARIs of 0.59 to 0.99. Analyzed alone, most of them lost the line identity. The ARI of inferCNV fell to 0.04 or less in every cohort, and that of Copy-KAT and CopyVAE to 0.21 or less on both MIX-seq panels. REFCON, run on each line alone, kept the lines apart in every cohort, with ARIs of 0.86 to 0.96. Figure 4B shows the esophageal lines run alone. SCEVAN split each line into separate groups, CopyKAT merged several lines together and inferCNV mixed all of them, whereas REFCON’s profiles kept the lines apart. Supplementary Figure 2 shows the embeddings of every method on these lines, run per cell line and pooled, and of expression alone. Expression alone, as a baseline, separated the lines with ARIs of 0.82 to 0.97 (Figure 4A).

**Fig. 4.**
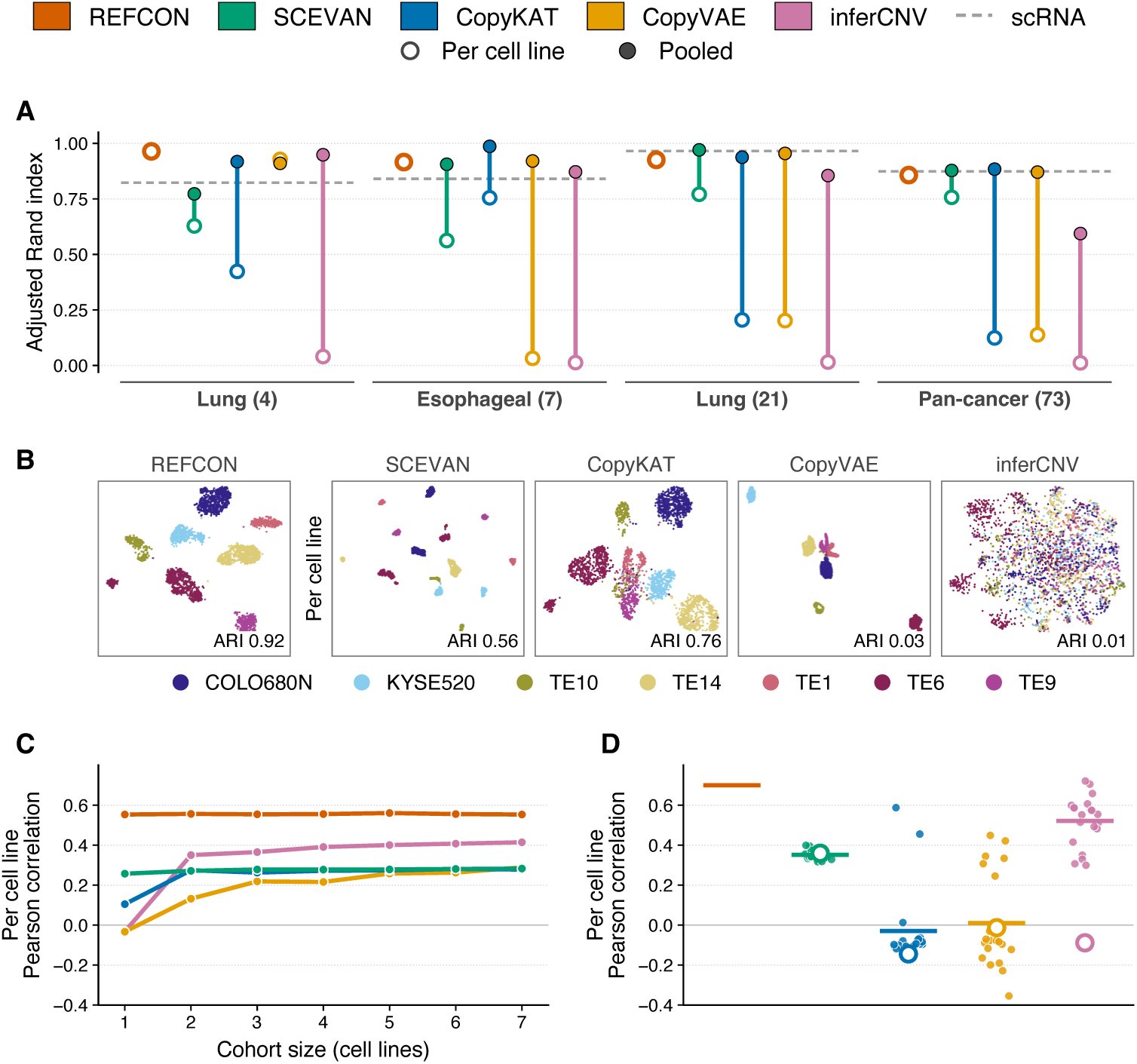
Copy number inference under changing cohort composition. **A** Adjusted Rand index (ARI) between each method’s clustering and the known cell line identity on four cohorts: CellBench lung (4 lines), CCLE esophageal (7), MIX-seq lung (21) and MIX-seq pan-cancer (73). The ARI is averaged over three clustering algorithms (k-means, Ward and Gaussian mixture) applied identically to every method. The reference-based methods were run on each line alone (per cell line) and on all lines of a cohort together (pooled). REFCON was run on each line alone. scRNA is the same clustering applied to log-transformed expression without copy number inference. **B** UMAP [32] embeddings of the seven esophageal lines, each line run on its own, with the ARI of each method. **C** Per cell line Pearson correlation with bulk DNA copy number for the seven esophageal CCLE lines, analyzed in cohorts of 1 to 7 lines, for REFCON and the four reference-based methods. Cohort size 1 is each line alone, sizes 2 to 6 are six random compositions each, and size 7 is all seven lines. Each point is the mean over the lines and compositions at that cohort size. **D** Per cell line Pearson correlation with bulk DNA copy number for one target line (NCIH1650) of the 21-line MIX-seq lung panel, analyzed together with each of the other 20 lines in turn, and alone. Each point is one partner, and the horizontal bar is the mean over the 20 partners. REFCON was run on the target line alone and has one value.

#### Copy number correlation across cohort compositions

We then measured how the correlation with bulk DNA changes with the cohort. We analyzed the seven esophageal lines in cohorts of increasing size: each line alone, six random compositions at each size from two to six lines, and all seven together. Figure 4C shows each method’s mean per cell line correlation at each size. Analyzed alone, CopyKAT, CopyVAE and inferCNV had correlations of −0.03 to 0.10, and with more lines they reached at most 0.41. We also analyzed one MIX-seq lung line, NCIH1650, together with each of the other 20 lung lines in turn. Figure 4D shows its correlation with each partner and when it was run alone. The correlation of these three methods ranged from −0.35 to 0.72 depending on the partner. SCEVAN found no normal cells in any of these cohorts, used its synthetic baseline instead and varied less. REFCON was run on each line alone and has one value per line, a mean of 0.55 for the esophageal lines and 0.70 for NCIH1650. Collectively, existing methods are sensitive to which cells are analyzed together. By inferring each cell’s copy number from that cell’s expression alone, REFCON decouples copy number inference from cohort composition.

### 2.4 REFCON follows the genomes of patient-derived organoids through culture and treatment without a reference

Patient-derived organoids consist solely of tumor cells and therefore provide no normal cells from which reference-based callers can estimate a baseline. We analyzed the pancreatic cancer cohort of Raghavan et al. [33], in which biopsies and the organoid lines derived from them were profiled by scRNA-seq across serial passages. The original study inferred copy number with inferCNV, using non-malignant cells from the biopsies as the reference. We ran REFCON and the four reference-based callers without a reference on 48 organoid samples from 14 lines of 13 patients and on 22 biopsies (Methods, Section 4.1). To show that REFCON follows these genomes, we evaluated the consistency of each line’s profiles across passages and culture conditions, followed one line through a TGF-*β*-induced change of expression state, and traced the clones of two lines derived from the same lesion before and after therapy.

#### Agreement between samples of a line

The original study authenticated each line by comparing its inferred copy number with that of the patient’s tissue and of earlier passages [33]. Following this rationale, we summarized each sample’s per-cell predictions as a profile over 39 chromosome arms and measured the agreement between two samples as the Pearson correlation of their profiles. Figure 5A shows this agreement for the 148 pairs of samples from the same line. REFCON’s profiles were highly consistent within lines, with a median Pearson correlation of 0.96, compared with 0.57 between samples of different patients. The next-best method, SCEVAN, reached a median of 0.68 within lines, and both SCEVAN and CopyKAT yielded no or negative correlation for some pairs. Per line, REFCON’s agreement was significantly higher than that of SCEVAN and CopyKAT in all 10 lines. CopyVAE’s profiles agreed less within lines. inferCNV returned values near zero for most genes, since without a reference its baseline is the mean of the sample’s cells, which share most of their genome. We further matched each organoid sample to the 22 biopsies by the correlation of the same method’s profiles. With REFCON, the closest biopsy was that of the sample’s own patient for 46 of the 48 samples, significantly more often than with SCEVAN (32) and CopyKAT (27). Supplementary Table 5 gives the agreement of every caller within lines, between patients and in each group of related samples, the biopsy matches and the tests.

**Fig. 5.**
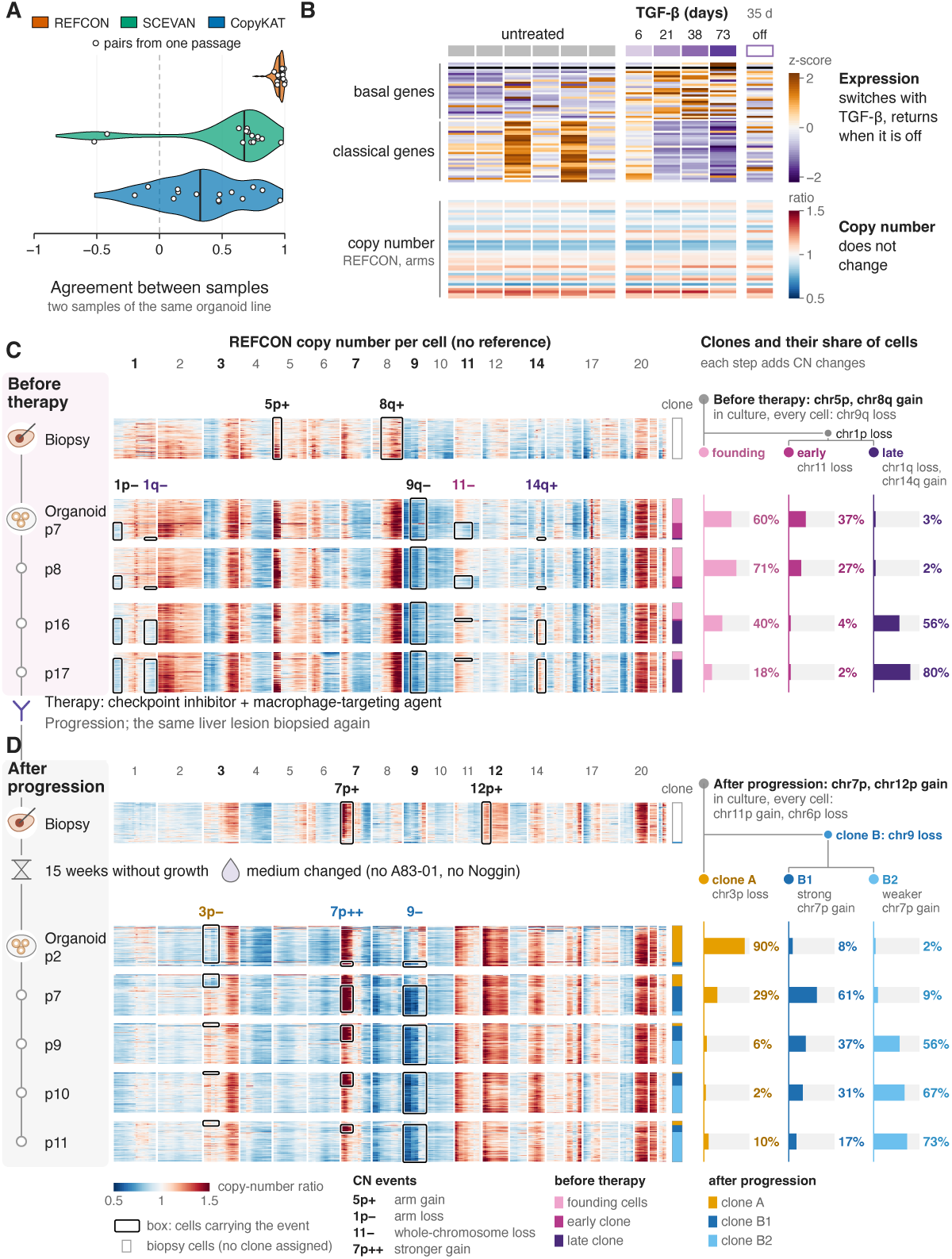
REFCON on patient-derived organoids without a reference. **A** Pearson correlation between the copy number profiles of two samples of the same organoid line (148 pairs from the 10 lines with two or more samples), for REFCON, SCEVAN and CopyKAT, each run without a reference. A sample’s profile is the median over its cells of each cell’s mean prediction per chromosome arm (39 arms). **B** PANFR0562 organoids untreated, under TGF-*β* for 6 to 73 days, and 35 days after its removal. Expression of the study’s 27 basal and 29 classical genes (z-score per gene) and REFCON’s copy number of 39 chromosome arms. **C** Patient PANFR0489 before therapy: the liver biopsy and the organoid line grown from it (passages 7 to 17). **D** The same lesion after therapy with a checkpoint inhibitor and a macrophage-targeting agent, and progression: the second biopsy and its organoid line (passages 2 to 11), which grew only after a 15-week arrest and a change of medium. In **C** and **D**, rows show REFCON’s per-cell copy number without a reference (100 cells per sample, sampled to the clone shares). The clone tree is built from shared events, with each clone’s share of cells per sample.

#### Copy number under a change of cell state

TGF-*β*, a cytokine of the tumor microenvironment, shifts PANFR0562 organoids from the classical toward the basal expression state, the two states the study describes in pancreatic cancer [33]. We analyzed samples of this line without TGF-*β*, after 6 to 73 days of exposure and 35 days after its removal. Figure 5B shows the study’s basal and classical genes alongside REFCON’s copy number in these samples. The expression state switched within 21 days of treatment, with the basal-minus-classical score rising from −0.05 in untreated organoids to between 0.59 and 0.71. After removal, the expression returned to its previous state (0.06). Since the expression state reverted, the copy number profile of the line is expected to stay the same. Accordingly, REFCON’s copy number did not change. The line’s seven arm losses and three arm gains were present at similar levels before, during and after TGF-*β* exposure. The profiles of the treated samples and of the sample after removal correlated with the mean untreated profile (Pearson 0.93 to 0.99) as closely as each untreated sample did with the mean of the remaining untreated samples (0.95 to 1.00). REFCON therefore did not mistake the change in expression state for a change in copy number. SCEVAN’s and CopyKAT’s profiles already disagreed between samples without TGF-*β*.

#### Clonal evolution without a reference

Unlike a reversible change of expression state, therapy and prolonged culture can select clones with distinct genomes, as the original study observed in culture [33]. One liver lesion of patient PANFR0489 was biopsied before treatment with a checkpoint inhibitor and a macrophage-targeting agent, and again after the tumor progressed. An organoid line was derived from each biopsy, PANFR0489 from the first and PANFR0489R from the second [33]. Figure 5C and D show both lines alongside their biopsies. The first biopsy was marked by gains of 5p and 8q, and the second by a 12p gain and a stronger 7p gain. Each line retained the events of its biopsy at every passage and additionally carried clonal events absent from it: a 9q loss in PANFR0489, and an 11p gain and a 6p loss in PANFR0489R. Supplementary Note 1 describes how the clones were identified. PANFR0489R expanded only after a 15-week growth arrest and a change of medium. In this line, a clone with loss of the whole chromosome 9, including *CDKN2A*, made up 10% of cells at passage 2 and 71% to 98% from passage 7 onward, replacing a clone with a 3p loss (Figure 5D). The original study reported this turnover using inferCNV with the non-malignant biopsy cells as the reference [33]. Without a reference, REFCON’s clones and their shares across passages closely matched those reported, with a Pearson correlation above 0.99 for both main clones. Supplementary Table 6 gives the clone shares of the study and of REFCON for every sample. SCEVAN, when supplied with the biopsy’s normal cells, separated the same clones in both lines, and Supplementary Figure 3 compares its clones with REFCON’s. In PANFR0489, we further observed a late clone that was not reported in the original study. Carrying a distal 1q loss and a distal 14q gain on top of a 1p loss shared with an early clone, it expanded from 3% of cells at passage 7 to 80% at passage 17, while the early clone, marked by loss of chromosome 11, declined from 37% to 2% (Figure 5C). As no DNA of these organoids was profiled, this turnover could not be validated. Organoid lines change both in expression state and in clonal makeup over culture and treatment. Without any reference, REFCON distinguished the two: its profiles stayed stable under a change of expression state and followed the clones that culture and therapy selected.

### 2.5 REFCON enables aneuploid cell classification with high specificity

Distinguishing aneuploid from diploid cells is a common use of inferred single-cell copy number profiles. Although REFCON performs copy-number profiling without a reference, aneuploid classification is operationally a within-sample comparison against putative diploid cells. Our classification procedure therefore compares the cells of a sample with confident-normal cells identified from gene expression in the same sample, using REFCON’s reference-free profiles (Methods, Sections 4.5–4.7). We scored the labels against the authors’ malignant and non-malignant annotations, used as proxies for aneuploid and diploid status (Methods, Section 4.1). Over-calling aneuploidy in normal cells is a recognized limitation of expression-based callers [16, 21]. We evaluated REFCON on tumor samples and on the settings where this problem arises, the normal cells inside tumors and fully normal samples.

#### Classification on tumors

We first evaluated classification on 94 tumors of eight cancer types from the Curated Cancer Cell Atlas [15, 34], comprising 59,699 malignant and 79,907 non-malignant cells, with each method run on the whole tumor sample and given no external reference (Methods, Section 4.1). Figure 6A shows the precision, recall and F1 of both classes, pooled over all cells. REFCON labeled every cell, whereas SCEVAN and CopyKAT left 11% and 13% of the cells unlabeled. Its aneuploid F1 was similar to that of the next-best method, SCEVAN (difference 0.01), while it labeled 94% of the non-malignant cells diploid, compared with 74% for SCEVAN. Across individual tumors, the aneuploid F1 of the two methods did not differ significantly, whereas REFCON’s diploid recall and SCEVAN’s aneuploid precision were each significantly higher. The advantage of REFCON was largest at both extremes of tumor content. In the 19 tumors with at most 20% malignant cells, it labeled 92% of the non-malignant cells diploid, compared with 53% for the next-best method, Copy-KAT. In the 16 tumors with more than 90% malignant cells, its aneuploid F1 was 0.84, compared with 0.76 for SCEVAN. Supplementary Table 7 gives the per-tumor comparison with its tests, the results by cancer type and by malignant share, and the results of 17 further patients from two studies. These include five colorectal patients whose colon tumor, adjacent normal colon and liver metastasis, among other tissues, were profiled together [35]. To examine the effect of cohort composition on tumor samples (Section 2.3), we also ran each patient on the colon tumor alone, on the colon tumor with the adjacent normal colon and on the liver metastasis alone. REFCON had the highest aneuploid and diploid F1 in every configuration, with an aneuploid F1 of 0.86 to 0.93, compared with 0.59 to 0.71 for the next-best method, CopyKAT. Supplementary Figures 4 and 5 show these runs and the patients’ expression embeddings, and Supplementary Note 2 describes the cases where the labeling failed.

**Fig. 6.**
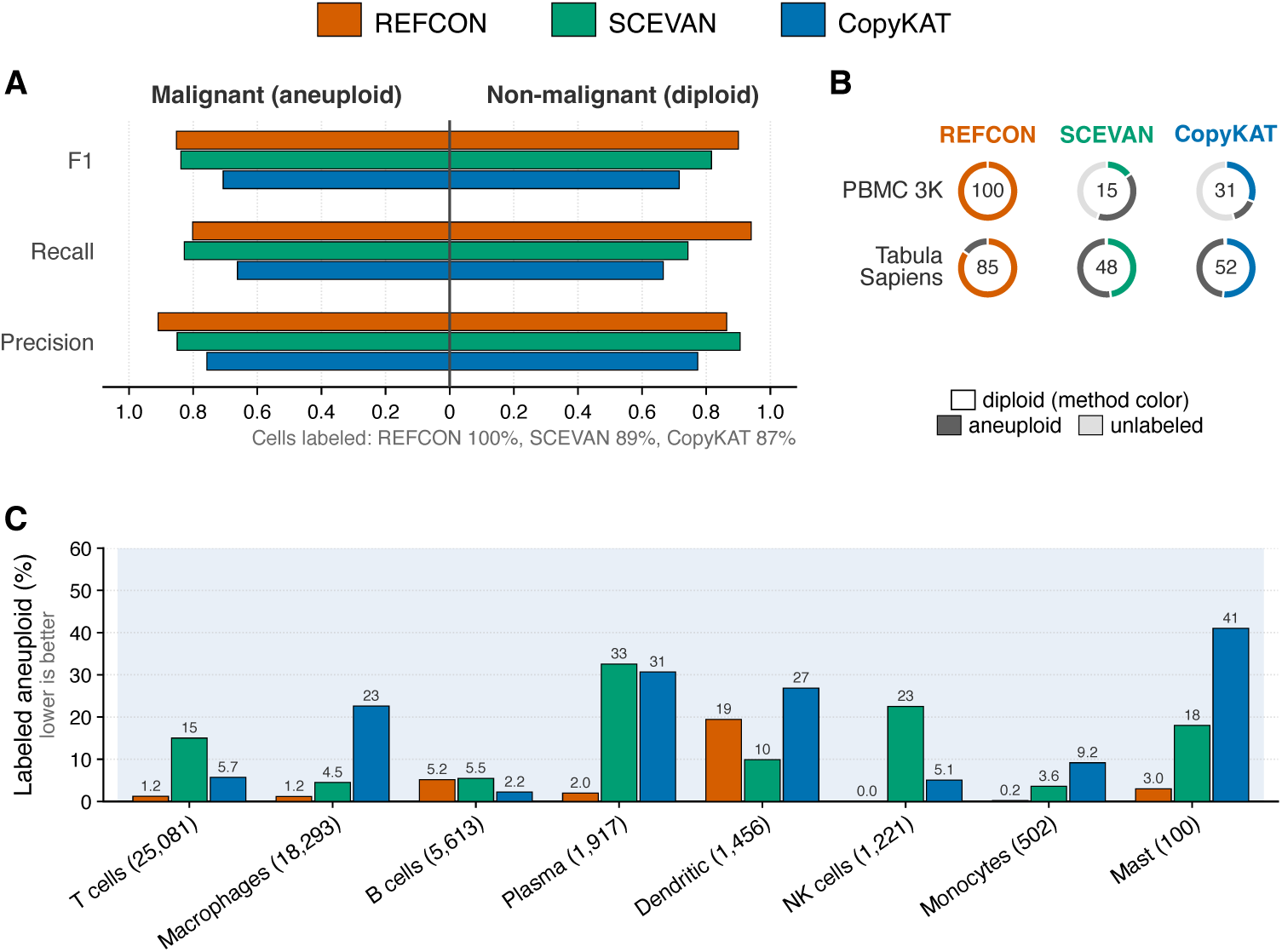
Per-cell aneuploid versus diploid classification on patient samples, normal controls and immune cells inside tumors. Methods compared: REFCON, SCEVAN and CopyKAT. inferCNV and CopyVAE return copy number without labeling cells as malignant or normal. On tumor samples, the authors’ malignant and non-malignant annotations are used as proxies for aneuploid and diploid status. A method may leave cells unlabeled, and an unlabeled cell counts as a miss for its annotated class. **A** Aneuploid- and diploid-class precision, recall and F1 on 94 atlas tumors of eight cancer types (59,699 malignant and 79,907 non-malignant cells), pooled over cells. **B** Labels on two fully normal control groups: the 10x Genomics PBMC 3K healthy-donor dataset (2,700 cells) and 14,400 healthy cells of 18 organs from Tabula Sapiens. Each cell is labeled diploid, labeled aneuploid or left unlabeled. The number in each ring is the percentage labeled diploid. **C** Percentage of immune cells labeled aneuploid in the 94 atlas tumors, by cell type. Cell types are the atlas annotations, taken from the original studies or made by the atlas and checked against known marker genes [34].

#### Immune cells inside tumors

Tumor samples also contain immune cells, which can be identified reliably by marker genes and, outside hematological cancers, are generally diploid [18]. They thus provide an internal set of normal cells whose expected label is known even without DNA. Figure 6C shows, for each immune cell type, the percentage of cells labeled aneuploid by each method. REFCON labeled 2.1% of all immune cells aneuploid, compared with 11.0% for the next-best method, SCEVAN. The only cell type in which REFCON labeled more cells aneuploid than SCEVAN was dendritic cells (19.4% against 9.9%), most of them in a single pancreatic tumor. Supplementary Table 8 gives these percentages by cancer type.

#### Specificity on normal controls

Following Schmid et al. [16], who tested callers on a healthy sample, we evaluated specificity on fully normal samples: the 10x Genomics PBMC 3K healthy-donor dataset [36] and healthy cells of 18 organs from Tabula Sapiens [25, 26], taken from donors who contributed no training or validation cells. Figure 6B shows how each method labeled the cells of these controls. REFCON classified every cell and labeled all 2,700 PBMC cells diploid, a specificity of 1.00. On the Tabula Sapiens cells, its specificity was 0.85, ranging from 0.42 to 1.00 per organ, compared with 0.52 for the next-best method, CopyKAT. On the PBMC control, SCEVAN and CopyKAT classified only 55% and 45% of the cells, yet labeled 73% and 32% of these aneuploid. Over the 19 normal-control samples, REFCON’s specificity was significantly higher than that of both methods. Supplementary Table 9 gives the call rate and specificity of each method on every control sample, together with the tests. REFCON thus labels every cell, classifies malignant cells as well as the best existing caller and largely avoids the over-calling of normal cells that hinders expression-based callers.

## 3 Discussion

REFCON reframes single-cell copy number inference as a learned local problem. Within windows of neighboring genes, it predicts local copy number from expression alone and assembles these predictions into a genome-wide profile per cell, with no reference at any step. This departs from existing expression-based callers, which normalize tumor expression against a reference of normal cells, whether matched, found within the sample or synthesized. Because expression is an indirect, regulation-shaped readout of copy number, no fixed rule maps one to the other. REFCON instead learns this mapping from cell lines with bulk copy number and from healthy cells, and applies it to each cell on its own. As it reads only the order of the genes and the genomic distances between them, invariant to library size and to which genes a protocol retains, a single trained model generalized across platforms and tumor types absent from training. In effect, the model has learned *how a copy number change imprints on a cell’s expression*, with no reference to compare against.

Removing the reference changes which samples can be analyzed and how their results can be compared. Cell lines and organoids, which contain no normal cells, become analyzable on their own. In organoids, REFCON kept the samples of a line consistent through culture and a change of expression state, and recovered a clonal turnover that previously required normal cells from the patient’s biopsy. Since cell lines and organoids evolve genetically in culture [37, 38], such profiles could help authenticate these models and monitor their drift from routine scRNA-seq. Because a cell’s prediction does not depend on the other cells analyzed with it, profiles from different samples, batches or studies are directly comparable, whereas the output of reference-based callers changes with the cohort. This makes REFCON suited to reanalyzing tumor atlases, such as the Curated Cancer Cell Atlas used here, whose samples differ widely in composition. REFCON also needs only a count matrix. Allele-based callers such as Numbat [39] require sequencing reads and genotypes, which much of the published scRNA-seq does not provide.

REFCON does not discard normal cells where they exist. Separating aneuploid from diploid cells is inherently a comparison within the sample, and REFCON uses the normal cells it finds only to label the cells, not to shape the predictions. Used this way, it classified malignant cells as accurately as the best existing caller, labeled every cell and labeled far fewer normal cells aneuploid, both in fully normal samples and among the immune cells inside tumors. When normal cells of the same sample are available, REFCON can also refine its predictions against them, which made them significantly more accurate than those of every method given the same cells. The reference thus becomes an optional enhancement rather than a requirement.

The method has certain limitations. Gene expression is an imperfect, regulation-shaped proxy for dosage, which bounds what any RNA-based method can recover, and the per-cell correlation of REFCON’s predictions with paired single-cell DNA remained modest (0.36 to 0.47). Because each profile is normalized to a mean of one, REFCON reports relative dosage and cannot detect absolute ploidy or a whole-genome doubling that scales every chromosome alike. REFCON was trained on autosomal genes only and was not benchmarked on the sex chromosomes or the mitochondrial genome, whose expression is shaped by X inactivation, sex and mitochondrial copy number rather than by chromosomal dosage. Detection improves with event size. Arm-and chromosome-scale alterations were detected with AUROC 0.75 to 0.81, whereas detection fell for shorter events and reached chance level for events of only a few genes, for REFCON and for every compared method (Supplementary Table 10). The scarcity of paired cell-level DNA and RNA shapes the method itself, which learns from bulk copy number labels and is judged against bulk and low-coverage single-cell profiles, all stand-ins for the per-cell truth we lack.

Most of the remaining limitations concern the labeling procedure rather than the copy number profiles. The procedure labels whole clusters, so a tumor population too small or too weakly altered to form its own cluster is labeled diploid. It also relies on the confident-normal cells found from gene expression, and a sample with fewer than two such cells receives a single sample-level call. Its specificity on healthy tissues varied by organ, from 0.42 to 1.00. Copy number labels are therefore a starting point, and calling a cell malignant requires additional checks on individual genes, such as cell type markers and known driver alterations.

As paired single-cell DNA and RNA datasets grow in number and depth, training and validating directly on high-depth per-cell measurements becomes the natural next step, and lifts the ceiling these stand-ins impose. Since much of the residual expression signal reflects co-regulation rather than dosage, integrating gene-regulatory or co-expression network structure to separate copy number effects from coordinated transcriptional programs could further sharpen per-cell resolution.

Copy number inference from scRNA-seq has so far served as a sample-dependent step, shaped by the reference each analysis provides, rather than as a measurement. By removing the reference, REFCON makes each cell’s copy number profile a property of its own expression, comparable across samples, studies and platforms. Copy number can then be read alongside expression in every cell of the tumors already profiled, the setting needed to study how it shapes clonal architecture and cell states at cellular resolution.

## 4 Methods

### 4.1 Data

Training. REFCON is trained on 64,379 cells (Table 1): 60,179 cells of 140 cell lines from five public scRNA-seq datasets, each line labeled with its bulk DNA copy number, and 4,200 healthy Tabula Sapiens cells [25, 26] from CZ CELLxGENE Discover [40], labeled diploid (*Ground truth*, below). The healthy cells are 300 from each of 14 organs, from six donors. Supplementary Data 1 lists every cell line and healthy cell of every split.

Preprocessing. Each training dataset was processed separately and kept its own genes. Cells were kept with at least 200 detected genes, at least 500 counts and at most 20% mitochondrial reads, and doublets predicted by Scrublet were removed. Genes were placed at their GRCh38 positions (GENCODE release 46). Mitochondrial and sex-chromosome genes were removed, together with HLA, immunoglobulin, T-cell receptor, ribosomal protein, hemoglobin, cell-cycle, dissociation-response, nuclear long noncoding RNA, pseudogene and olfactory receptor genes, and genes detected in fewer than 10% of the dataset’s cells. Test cohorts were not preprocessed this way. Their genes were placed at the same GRCh38 positions, and only mitochondrial and sex-chromosome genes were removed. In the copy-number benchmark cohorts, genes without counts in the analyzed sample are also removed. The gene removal and zeroing augmentations of training (Section 4.3) expose the model to gene sets of varying density.

#### Validation and development

Two validation sets, both disjoint from the test cohorts, are used only for hyperparameter tuning and checkpoint selection. Seven CCLE lines that appear in no training set, one from each of seven randomly chosen tissues, probe generalization to unseen cell lines. 1,200 Tabula Sapiens cells from two donors who contribute no training or test cells measure false copy number events on diploid cells. The classification thresholds (*τ*_1_ and *T_σ_*; Section 4.7) are fixed on three development melanoma patients of Tirosh et al. [17] (tir 79, tir 80, tir 88), the three with the largest minority class (the smaller of their malignant and non-malignant populations), so that both a diploid and an aneuploid population are well represented. They are disjoint from the 17 test patients.

#### Test cohorts

Table 1 lists the test cohorts by analysis. All are line- or patient-disjoint from training and validation, and no held-out cell line shares a DepMap model with a training line. The 73 MIX-seq lines [27] are the held-out lines of one SNP-demultiplexed pool of 99 lines (DMSO, 24 h). The 49 with DepMap bulk copy number are scored against it, and all 73 are used for clustering. The 21 lung lines among them (1,268 cells) form the lung panel. The public Wang et al. [35] data hold seven patient identifiers (the study reports six). Five are the test patients, and two hold CD45^+^ cells only.

#### Atlas tumors

For classification inside tumors, we used the tumor samples of the Curated Cancer Cell Atlas (3CA) [15, 34] whose expression is given as raw UMI counts. They come from eight studies, one per cancer type: glioblastoma [7] (its 10x samples), pancreatic cancer [33], colorectal cancer [44], melanoma [45] (sorted CD45^+^ cells, some of which the atlas annotates as malignant), liver cancer [46], ovarian cancer [47], head and neck cancer [48] and renal cell carcinoma [49] (seven clear cell tumors and one chromophobe tumor). They were profiled with 10x, except the pancreatic (Seq-Well S^3^) and melanoma (MARS-seq) studies. We kept the tumor samples with at least 10 malignant cells and excluded samples of normal tissue (as the atlas annotates them, which for four kidney pairs is the reverse of the GEO sample titles), which leaves 94 tumor samples. We scored the cells the atlas annotates as malignant (59,699) against all cells it annotates as non-malignant with a cell type (79,907; *Ground truth labels*, below). False positives were also scored on the immune cell types alone: T, NK, B and plasma cells, macrophages, monocytes, dendritic and mast cells (54,183 cells). Stromal and epithelial cells, which can carry copy-number changes, are not scored. Each tumor sample was processed whole by every method and given no external reference. The atlas also holds the melanoma study of Tirosh et al. [17], but only as TPM values (Smart-seq2), so it is not among the 94 atlas tumors.

#### Patient-derived organoids

We used the Curated Cancer Cell Atlas [15, 34] release of Raghavan et al. [33]. We kept the organoid and cell line samples with at least 30 cells and, as biopsies, the malignant cells of the 22 of the 23 atlas pancreatic tumors that have at least 30 such cells. Three organoid samples duplicate others under a second name (identical count matrices) and were removed. The two samples of the CFPAC1 cell line were excluded. This leaves 48 organoid samples (36,752 cells) from 14 lines of 13 patients. Ten lines have two or more samples. Passage order gives the time order within a line. The expression state score is the study’s scBasal signature score minus its scClassical score [33], averaged per sample.

#### Ground truth labels

Three kinds of ground truth are used, one per evaluation regime. *Cell line bulk copy number*, for the cell line training labels and the per cell line evaluation, is taken from the DepMap 24Q4 release [50, 51], which calls gene-level absolute copy number from whole-genome and whole-exome sequencing of each line. The healthy Tabula Sapiens training and validation cells carry no measured copy number and are labeled diploid by construction (copy number two at every gene, so every bin’s target ratio is one). *Per-cell copy number*, for the single-cell evaluation, is the paired single-cell whole-genome-sequencing profile: For the DNTR-seq cohorts, it is called from the paired DNA reads with BWA-MEM [52], SAMtools [53], and AneuFinder [54]. Supplementary Note 3 describes this pipeline. Most HCT116 cells of the DNTR-seq study were irradiated or treated with etoposide [11]. Their truth comes from each cell’s own DNA, so it includes any copy-number changes the treatment caused. For the scONE-seq astrocytoma, it is the single-cell DNA matrix that Yu et al. [12] deposited (GSE185269): read depth in variable bins of about 500 kb (median 513 kb), normalized to each cell’s mean. Their integer copy number calls were not deposited. We multiply the matrix by two, so each cell’s mean is two copies, and give each gene the value of the bin that contains its midpoint. The values therefore stay continuous, and the loss and gain thresholds (*<* 1.5 and *>* 2.5) are applied to them. Correlations do not depend on this scaling. Of the 1,055 nuclei with a DNA profile, 843 also have an RNA profile, and 840 pass RNA quality control (3 main-clone nuclei are removed). These 840 are the cohort. *Per-cell malignant versus non-malignant labels*, for the classification evaluation, are the author annotations of the Tirosh melanoma [17] and Wang colorectal [35] cohorts and the atlas annotations of the 94 atlas tumors, used as the aneuploid versus diploid ground truth; none of these cohorts has paired single-cell DNA data. In Tirosh et al., cells with copy-number changes inferred from expression were labeled malignant. These cells also formed tumor-specific expression clusters, the non-malignant cells were typed by marker genes, and the inferred copy-number pattern was confirmed by bulk whole-exome sequencing in two tumors. In Wang et al., cells were annotated from marker gene expression and a 7-AAD/CD45 sort gate without copy-number-based refinement. The atlas takes the annotations of the original studies and annotates cells itself where these are missing or limited. It verifies every annotation in three ways: cell types must cluster apart, the top differentially expressed genes of each type must match known marker genes, and cells annotated as malignant must carry inferred copy-number aberrations [15, 34, 55]. We map malignant to aneuploid and non-malignant to diploid, and exclude cells the authors left unresolved. The two are close but not identical. Malignant cells can be near-diploid and some non-malignant cells can be aneuploid, so scores against these labels have a ceiling below 1. The loss and gain thresholds applied to the ground truth are given with the metrics (Section 4.9).

### 4.2 Model architecture

REFCON takes a window of *L* ≤ *C* per-gene expression values 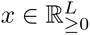 (raw UMI counts, or TPM-derived values where only TPM is released) in genomic order, padded to a fixed length *C* with zeros, where *C* is the maximum context length. The inference pipeline uses the genes’ GRCh38 coordinates and chromosome assignments to cut the windows (Section 4.4). The model itself receives, with each window, only the genes’ offsets from the window’s first gene. The pipeline below is the same for both ensemble members (Section 4.3).

1. log1p: *x* ← log(1 + *x*).
2. Masked InstanceNorm: per-cell mean and standard deviation are computed over the real gene prefix *x*_1:_*_L_* only; padding is held at zero. A learnable affine (*γ, β*) is applied to the prefix. This removes per-cell library-size and protocol-scale differences and enables a single checkpoint to work across droplet (10x, BD Rhapsody, Seq-Well S^3^), plate-based (Smart-seq2 [56], MARS-seq) and paired DNA and RNA (DNTR-seq, scONE-seq) data.
3. One-dimensional convolutional binner. A single 1D convolution (bin size *b* = 8 genes: kernel size 8, stride 8, 64 output channels) followed by a GELU activation. This reduces the sequence to *C/*8 bin tokens of dimension 64.
4. Convolutional block. Two 1D convolutions (kernel size 7, padding 3, 128 output channels) with BatchNorm and GELU activations, followed by a post-block LayerNorm. At inference, BatchNorm uses the statistics stored during training, so a cell’s prediction does not depend on the other cells in the batch. In a window spanning two chromosomes, the block is applied to each chromosome’s bins as if they formed a window of their own, with the other chromosome’s bins replaced by the binner output of an empty bin.
5. Four-layer Transformer. *d*_model_ = 128, 4 attention heads, feed-forward dimension 512, dropout 0.1, pre-norm. Rotary position embeddings (RoPE) [57] are applied to the query and key projections, using each bin’s genomic offset rather than its index. A bin’s offset is the mean distance, in megabases along GRCh38 with the chromosomes concatenated, of its genes from the window’s first gene in reading order, so that a reversed window reads as the mirrored genome. Because rotary embeddings depend only on differences of these offsets, the attention between two bins depends only on their genomic distance. Attention is masked to valid bins (those whose 8-gene window contains at least one actual gene’s expression) and, in a window spanning two chromosomes, to bins of the same chromosome.
6. Linear ratio head. A two-layer MLP (Linear 128 → 64, GELU, dropout 0.1, Linear 64 → 1), producing one scalar ratio prediction per bin. Bin predictions are broadcast across their 8 constituent genes for gene-level evaluation.

In a window that spans two chromosomes (a bridge window, Section 4.4), the two chromosomes share only the window’s input normalization and its target, the copy number of each bin relative to the window mean. The model locates the boundary from a gap of 1,000 Mb inserted between chromosomes in the offsets. The gap never enters as a distance, because no attention or convolution crosses the boundary, and no bin contains genes of two chromosomes.

Each ensemble member has 975,299 parameters and a 3.9 MB checkpoint; the published 2-member ensemble therefore totals ∼1.95 M parameters and ∼7.9 MB. Neither the megabase positions nor the separation of chromosomes adds parameters. The main architectural and training choices are ablated in Section 4.10.

### 4.3 Training

Training data is structured as *window* s: each epoch draws one window per cell, cycling over the six training datasets, so that the healthy cells supply 6.5% of an epoch’s windows. With probability 0.5 the window is a bridge window spanning the boundary between two adjacent chromosomes, built as at inference (Section 4.4) but starting at a random bin-aligned position. Otherwise it covers a single chromosome chosen at random for that cell. With probability 0.5 the window is presented in reversed gene order, its expression, offsets and targets reversed together. For a chromosome whose gene count *L_c_* exceeds *C*, a contiguous sub-window of length *C* starting at a uniformly random offset ∈ [0*, L_c_* −*C*] is sampled per epoch (chunk-stochastic training); for *L_c_* ≤ *C*, the full chromosome is used and zero-padded to *C*, with attention masked to the real-gene prefix. Each chromosome in a window starts at a uniformly random offset of 0 to *b* − 1 genes, so that every grouping of genes into bins is seen while chromosome boundaries stay on bin edges. Chromosomes are drawn with unequal weight. A chromosome carrying at least one copy-number loss in any of its dataset’s profiles is given twice the weight of a neutral one, so loss-bearing chromosomes are sampled about twice as often over an epoch. This offsets the strong class imbalance of the cell line training profiles, which contain mostly gain events (typical mean copy number 2.8 to 3.7). Neutral genes outnumber loss genes by roughly 20:1 and gain genes outnumber loss by roughly 40:1.

Three perturbations change the counts of a window but never its targets. With probability 0.5, every count is thinned binomially at a rate drawn log-uniformly from [0.1, 1]. With probability 0.25, a random fraction of the window’s genes, uniform in [0, 0.5], is removed, and the remaining genes keep their order, offsets and targets. With probability 0.25, a random fraction of the genes, uniform in [0, 0.3], is set to zero. They mimic shallower sequencing, smaller gene panels and genes that are present but not captured.

REFCON is trained to predict a per-bin copy-number ratio. Within a training window, gene *g* has copy number *c_g_* (the bulk copy number of its cell line, or two for a healthy cell); dividing *c_g_* by the window’s mean copy number *c̄* = mean*_g_ c_g_* yields a per-gene ratio *c_g_/c̄* that averages to one across the window. The target for bin *j*, written *r_j_*, is the mean of these ratios over the (at most eight) genes the bin spans, and the model outputs one prediction *r̂_j_* per bin (^· denotes a model prediction throughout). The objective is a Huber (smooth-*L*_1_) loss over the bins:

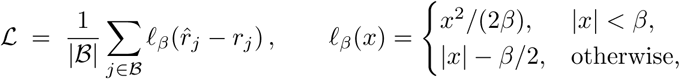

It is taken only over the bins B of the window whose eight-gene span contains at least three real, non-padded genes. Padded bins are not considered. We set *β* = 1, the standard Huber transition.

We use AdamW (initial learning rate *η* = 10*^−^*^4^, weight decay 0.01), gradient clipping at ∥∇∥ _2_ = 1.0, batch size 64, and a cosine learning-rate decay from 10*^−^*^4^ to 10*^−^*^6^ over 150 epochs, fixed in advance and independent of any validation score. There is no early stopping. After every epoch, the complete inference pipeline (Section 4.4) is run on a fixed set of 350 cells of the seven unseen validation lines (50 per line, separate from the 150 cells per line on which the validation scores of the Supplementary Tables are computed), and the mean per cell line Pearson correlation of the stitched profiles with the lines’ bulk copy number is recorded. Weights are saved every 10 epochs, and a run’s weights are the snapshot at the maximum of the 10-epoch moving average of this score, centered on the snapshot (epochs *e* − 5 to *e* + 4). A run takes about 6 hours (150 epochs of about 146 s) on a single NVIDIA GeForce RTX 2080 Ti. Three runs with different random seeds were trained at *C* = 2,048 with this protocol. The released ensemble averages the two with the highest Pearson correlation on the validation lines, whose snapshots are those of epochs 120 and 100.

REFCON is reported throughout the manuscript as the ensemble of these two checkpoints. Each checkpoint produces an independent matrix of ratios (*n*_cells_*, n*_genes_); the ensemble prediction is the mean over members followed by the renormalization per cell to mean = 1 (computed only on finite entries). No learned weighting and no per-dataset tuning is used. Supplementary Table 11 gives the metrics of each trained seed, each two-model pair and the three-model ensemble.

### 4.4 Inference

At inference, each cell’s expression vector is segmented into per-chromosome chunks using the chromosome assignment and the genomic start coordinate (GRCh38) supplied as input metadata. Genes are pre-sorted by genomic position within each chromosome. A sliding window of length *C* is run over each chromosome (chromosome-internal windows, stride *C/*2) and over each pair of consecutive chromosomes (bridge windows of at most *C* genes that span the boundary between the two chromosomes, with the boundary on a bin edge and at least two bins on each side; stride *C/*4 across the boundary). Each window is passed through the model twice (once in original gene order, once with the real-gene prefix reversed), and the two bin-level outputs are averaged after un-reversing (bidirectional averaging). Each window carries its genes’ offsets from its first gene, which are reversed together with the window in the reversed pass. Running the forward pass only is available as an option that halves the model passes at a small cost in accuracy (per-cell Pearson correlation 0.417 instead of 0.429 on HCT116; Table 2).

**Table 2.** Inference runtime and memory of the released model on four compute platforms.

| Compute platform | Throughput<br>(cells per<br>second) | Throughput,<br>forward pass<br>only (cells per<br>second) | Peak accelerator<br>memory | Peak host<br>memory |
| --- | --- | --- | --- | --- |
| NVIDIA RTX 2080 Ti | 17.2 | 27.0 | 1.0 GB | 1.7 GB |
| Intel Xeon Gold 6140 (16<br>threads) | 0.91 | 1.78 | — | 1.9 GB |
| Apple M4 Pro integrated<br>GPU | 5.99 | 11.5 | Unified memory | 2.0 GB |
| Apple M4 Pro CPU (10<br>performance cores) | 0.89 | 1.76 | — | 2.5 GB |

Measurements use the released two-model ensemble (two seeds at context length 2,048, 975,299 parameters per member), with the members evaluated one after the other. GPU measurements used the 1,468 HCT116 cells with paired DNA (17,685 genes), and CPU measurements the first 500 of these cells, all at 512 cells per batch. Throughput is given for the default inference, which averages forward and reversed passes, and for the forward pass only. Peak accelerator memory is the PyTorch allocation. On the Apple M4 Pro, accelerator and host memory share one unified memory and cannot be separated. The NVIDIA and Intel runs shared their server with other jobs.

The per-window bin predictions are stitched into one genome-wide profile by a joint log-space least squares. Index the windows *i* = 1*, … , W* . Window *i* predicts a bin-level ratio *r_i_*[*u*] at each global bin *u* it covers. The model emits a mean-one ratio, so each window’s profile is fixed only up to an overall scale. That scale is one additive log-offset *s_i_* per window, and the stitching recovers it. Two overlapping windows *i* and *j* that share a bin *u* must agree there after rescaling, which in log space is the linear constraint

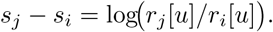

We stack one such equation for every shared bin of every overlapping window pair. This gives an overdetermined system *A*s = y, where s = (*s*_1_*, … , s_W_* )*^⊤^* is the vector of per-window log-scales. Each row of *A* has one +1 and one −1, in columns *j* and *i* (other entries are zero) and the matching entry of y is the log-ratio above. Every constraint is a difference *s_j_* − *s_i_*, so s is determined only up to a global additive constant. This single gauge freedom is the overall log-scale. We fix it by adding one weighted equation *λ s*_1_ = 0 that pins the first window’s scale. We set *λ* = 10^4^, large enough to make the pin effectively exact. The system is solved by ordinary least squares:

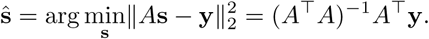

The solution *ŝ_i_* is each window’s estimated log-scale. We rescale window *i*’s bin predictions by *e^−s^*^^^*^i^* and average across the windows covering each bin. We then renormalize the profile to mean one over valid bins. This removes the gauge, so the result does not depend on *λ*. Finally, each bin value is mapped to its genes, with each 8-gene bin repeated across its group member genes.

Three of these choices are dictated by the structure of copy number rather than by tuning. Copy-number segments do not cross chromosome boundaries, so a segment on one chromosome and a segment on another are distinct events. The model therefore never relates the expression patterns of two chromosomes (Section 4.2; Supplementary Table 12, Part D). Within-chromosome windows fix the relative copy number inside each chromosome, but leave each chromosome’s overall level as a free scale. The bridge windows tie those scales together, placing all chromosomes on one genome-wide scale. Copy number is also invariant to the direction a chromosome is read, and the model is trained on windows in both reading directions. Averaging each window with its gene-order-reversed counterpart therefore acts as a two-view ensemble of the same window.

Supplementary Table 13 gives the accuracy with each stitching component removed or replaced. The bridge windows contribute most, and bidirectional averaging is a small robustness step. A sequential per-chromosome-then-bridge chaining scores slightly higher on the validation lines and lower on all three paired single-cell-DNA cohorts. The pin weight *λ* leaves every metric unchanged across five orders of magnitude. The joint least squares is kept because it fits all window overlaps at once, whereas a sequential chaining depends on the order in which windows and chromosomes are joined and carries each step’s error into the next.

### 4.5 Identifying confident-normal cells

The first step of the classification identifies confident-normal cells, which serve as the in-sample diploid reference of the labeling. They are found with the marker-based normal-cell detector of SCEVAN [19], except that every passing cell is kept (SCEVAN keeps at most the 30 top-scoring cells). Each cell is scored against SCEVAN’s 12 normal-lineage gene-set signatures and is called confident-normal when, for some signature, NES ≥ 1 and FDR *<* 10*^−^*^10^. The scores are relative to the other cells of the sample, so a sample without internal contrast yields few or no such cells. A sample with fewer than two confident-normal cells is treated as pure (Section 4.7).

### 4.6 Clustering of the remaining cells

Cells not flagged confident-normal in Section 4.5 are partitioned into clusters by their predicted CN-ratio profiles (the unrefined predictions). Let *X* ∈ R*^n×m^* be the ensemble’s gene-level prediction matrix for the *n* non-confident-normal cells across *m* genes, with out-of-coverage entries imputed to their per-gene mean over cells. We project *X* onto its top five principal components and standardize each to zero mean and unit variance. This gives *Z* ∈ R*^n×^*^5^. We fit a Gaussian mixture with *G* components and full per-component covariance to the rows of *Z*,

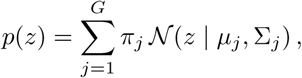

where *π_j_*, *µ_j_*, and Σ*_j_* are the mixing weight, mean, and covariance of component *j*. We fit the mixture for each *G* from one to eight with five random restarts. The component count is chosen by the Bayesian information criterion,

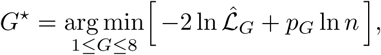

with L^^^*_G_* the maximized likelihood and *p_G_* the number of free parameters of the *G*-component model. Each cell *i* is assigned to its most probable component, *c_i_* = arg max*_j_ P* (*j* | *z_i_*), deterministically given a fixed random seed.

The number of retained principal components (*n*_pcs_ = 5) is a fixed default rather than a tuned value. We checked its effect on the validation data of Section 4.1, comprising the seven held-out CCLE validation cell lines and the three development patients. With five PCs, the BIC selects seven clusters on the seven CCLE validation lines (Adjusted Rand Index 0.993), and the development patients’ cells are labeled with an accuracy of 0.981 (over all 1,664 author-labeled cells). The labels depend little on this choice: from one to ten PCs the accuracy stays between 0.979 and 0.996, except at nine PCs (0.915). We use five PCs as the default. The full per-component covariance form is chosen because it allows each Gaussian component to take an arbitrary ellipsoidal shape in the PCA-projected space. The upper bound of eight on *G* is arbitrary. It only caps how many components the BIC search may select, and may be raised for samples expected to carry many sub-populations.

### 4.7 Labeling clusters as diploid or aneuploid

Each cluster produced by Section 4.6 is labeled diploid or aneuploid by comparing the CN-ratio dispersion of its cells with that of the confident-normal cells of Section 4.5. Confident-normal cells themselves are labeled diploid by default; every other cell inherits the label assigned to its cluster.

Let D ⊆ {1*, … , n*_cells_} denote the indices of the confident-normal cells in a sample, and let *r̂_i,g_* denote the predicted CN ratio of cell *i* at gene *g* (with out-of-coverage entries imputed as in Section 4.6). The dispersion of cell *i* is the standard deviation of its predicted profile across genes, *d_i_* = std*_g_ r̂_i,g_* . The diploid reference is the median dispersion of the confident-normal cells,

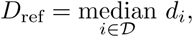

and the dispersion ratio of cluster *c* is the median dispersion of its members relative to that reference,

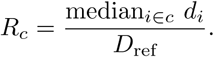

The cluster is labeled

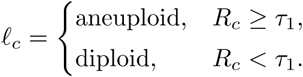

The threshold *τ*_1_ = 1.05 is fixed across samples and is not per-dataset tuned. It was set on the three development patients (Section 4.1). On these patients the diploid and aneuploid clusters overlap in dispersion ratio: one 27-cell normal cluster (1.060) lies above the least dispersed tumor cluster (1.051), so no threshold separates them. Every *τ*_1_ from 0.94 to 1.05 labels 7 of the 8 development clusters (holding 98.2% of the 1,482 clustered cells) correctly, and *τ*_1_ = 1.05, the top of that range, keeps the largest margin above the confident-normal reference, whose ratio is one by construction. The cluster dispersion ratio requires a confident-normal cell set, so it is undefined on a pure-aneuploid sample such as a CCLE validation cell line, and those lines do not enter the derivation of *τ*_1_. The exact value of *τ*_1_ is not critical. Per-patient classification stays robust across a broad range of *τ*_1_ around 1.05 (mean test-patient accuracy above 0.90 for *τ*_1_ from 0.96 to 1.30). Supplementary Figure 6 shows the classification of each patient across *τ*_1_. A full precision-recall sensitivity analysis over *τ*_1_ is provided in Supplementary Figure 7.

The cluster-and-threshold paradigm of this step follows the tumor and normal cell classification of CopyKAT [18]. Both methods cluster cells and call clusters by a per-cluster statistic computed against an in-sample diploid reference.

When a sample yields fewer than two confident-normal cells (Section 4.5), the diploid reference dispersion *D*_ref_ is undefined and the cluster dispersion-ratio rule above does not apply. We instead assume the sample is internally homogeneous (pure-diploid or pure-aneuploid) and assign one shared label to every cell of the sample. The decision uses an absolute aneuploidy score derived from the predicted profiles themselves: The mean dispersion of its cells,

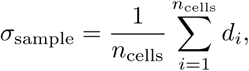

which is high when the predicted profiles depart from the cell’s own mean ratio (the signature of aneuploidy) and low when the predicted profiles are nearly flat. The sample is called aneuploid if *σ*_sample_ ≥ *T_σ_* and diploid otherwise, with *T_σ_* = 0.125. Unlike the cluster dispersion ratio, *σ*_sample_ is an absolute measure of per-cell dispersion, so it can be set on samples that carry no diploid reference. It was found empirically, on data disjoint from the test set, by treating the malignant and the non-malignant cells of the three development patients as separate pure cohorts, together with the seven pure-aneuploid CCLE validation cell lines (Section 4.1). The pure-diploid cohorts (the non-malignant cells) have low dispersion and the pure-aneuploid cohorts (the malignant cells and the CCLE lines) high dispersion, and *T_σ_* = 0.125 sits at the midpoint of the gap between them (0.115 to 0.135), the maximum-margin operating point. This branch returns a single sample-level call rather than per-cell labels, and applies only when a sample carries no in-sample diploid population.

### 4.8 Refining the predicted ratios

An optional step refines the predicted CN ratios against a panel of normal cells of the same sample. It runs after labeling, does not change the labels, and sharpens the relative CN-ratio profiles of the aneuploid cells. The classification steps of Sections 4.5–4.7 therefore run on the unrefined reference-free predictions. Refining beforehand would flatten the confident-normal profiles that set the labeling’s reference dispersion *D*_ref_, distorting the cluster dispersion ratio *R_c_*.

Let D ⊆ {1*, … , n*_cells_} be the indices of the diploid panel, a set of normal cells of the same sample (the author-annotated non-malignant cells in Section 2.2, or the confident-normal cells of Section 4.5), for a sample of *n*_cells_ cells and *n*_genes_ genes. Let *r*^ ∈ R*^n^*^cells^ *^×n^*^genes^ be the predicted CN-ratio matrix, with entry *r̂_i,g_* the ratio of cell *i* at gene *g* (out-of-coverage entries imputed as in Section 4.6). The refinement proceeds in three steps.

1. Per-gene consensus subtraction. For each gene *g*, the reference panel’s mean deviation from one is

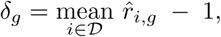

and this per-gene bias vector *δ* = (*δ*_1_*, … , δ_n_*_genes_ ) is subtracted from every cell, *r̂^′^* = *r*^− *δ* (mapped over cells).

2. SVD bias-mode removal. Let *R* ∈ R*^|D|×n^*^genes^ be the centered reference residual *R* = *r̂^′^* − 1, formed from the reference rows of *r̂^′^*. Its singular value decomposition is *R* = *USV ^⊤^*, with *S* the diagonal matrix of singular values. Let *V_k_* ∈ R*^k×n^*^genes^ hold the top *k* right singular vectors as rows. These *k* modes are projected out of every cell,

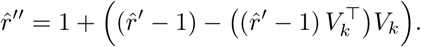

3. Per-cell renormalization. Each cell’s refined profile is divided by its mean over finite entries.

For the released ensemble, each member’s prediction is refined and the refined profiles are averaged.

The single hyperparameter is the SVD rank *k*. We use *k* = 3, and Supplementary Table 14 gives its sensitivity. The panel should consist of normal cells of the same sample, whose true profile is flat, so that the refinement removes only a shared bias.

The refinement is applied to every cell in the cohort; the per-gene bias and the modes *V_k_*are estimated from the diploid panel alone and applied to each cell independently, so a given cell’s refined profile does not depend on which other cells are present, other than the normal panel.

### 4.9 Benchmark protocol

We compared REFCON with four expression-based callers (SCEVAN, CopyKAT, CopyVAE and inferCNV), each run with one fixed configuration, and with the allele-based caller Numbat on HCT116 and A375. Supplementary Note 4 gives each method’s configuration, the injection of the pseudo-references and a comparison without filled genes. Each method’s predictions are placed on the dataset’s own genes, and genes a method does not call are filled with the cell’s median prediction. When several cell lines are analyzed together, each line is scored on the genes it expresses, as when it is analyzed alone. REFCON was run on each line alone. The methods report copy number on different scales, from ratios and copy numbers to log-scale values. We therefore use metrics that need no cutoff on the predictions, and apply thresholds to the ground truth only. A gene is labeled loss if its copy number is *<* 1.5, neutral if in [1.5, 2.5], and gain if *>* 2.5. Labeling loss and gain relative to each cell’s ploidy instead leaves the method ranking unchanged, as Supplementary Table 15 shows. We report Pearson and Spearman correlation, and AUROC and AUPRC for loss and gain. AUPRC is included because the copy number classes are imbalanced.

All copy number metrics are computed per cell and averaged over cells, or on pseudobulk profiles against bulk DepMap copy number for the per cell line cohorts. Spearman correlation, AUROC and AUPRC depend only on the rank order of each method’s predictions.

Classification is scored against the authors’ malignant versus non-malignant annotation, and a cell a method leaves unlabeled counts as a miss. Precision, recall and F1 are pooled over the cells of each evaluation group. On the fully normal controls, specificity is computed among the cells a method labels, and the call rate is the fraction of cells it labels. Cell line clustering is scored by the adjusted Rand index at the known number of lines. On the astrocytoma, the per-cell metrics are also averaged within each clone defined by the paired DNA. Supplementary Note 5 gives the reasons for filling missing genes and for thresholding the ground truth only, and the protocol of the clustering and cohort composition analyses.

### 4.10 Ablation protocol

The main architectural and hyperparameter choices in REFCON are documented in dedicated supplementary tables as referenced below. Each table compares the released configuration with variants that change one choice. Supplementary Table 12 gives the ablations of the released model and Supplementary Table 16 the control without the transformer. Supplementary Table 17 gives the accuracy of the released model and of the models trained without gene removal and zeroing or without count thinning when genes are removed or each cell’s counts are thinned. Ablations are scored on the held-out test cohorts and, for retrained models, also on the seven validation lines. Each retrained variant uses the released recipe with one choice changed and the two seeds of the released members (seeds 2 and 3), and its two models are averaged per cell like the release. Differences smaller than the seed spread of the released recipe (validation Pearson 0.701 to 0.716 across three seeds) do not distinguish a variant from the release.

#### Architecture ablations

Models without rotary position embeddings, with rotary positions over the bin index instead of genomic distance, without per-cell input InstanceNorm, or with InstanceNorm replaced by BatchNorm were each retrained at *C* = 2,048.

#### Transformer contribution

We evaluated REFCON’s exact inference pipeline (8-gene bins, overlapping 2,048-gene windows and bridge windows, and joint stitching) with the transformer replaced by plain averaging, thereby isolating the contribution of the learned model from that of the surrounding inference pipeline.

#### Context lengths

We evaluated context lengths *C* ∈ {512, 1,024, 2,048}, two seeds each, and compared these models with the released two-model ensemble.

#### Leave-one-out of the training additions

Starting from the released recipe, we left out one addition at a time (the healthy training cells, count thinning, gene removal and zeroing, or the separation of the two chromosomes in bridge windows, keeping the megabase positions) and retrained.

#### Refinement SVD-rank sensitivity

We evaluated *k* ∈ {0, 1*, … ,* 10} on the scONE-seq astrocytoma, refined against the 63 confident-normal cells REFCON detects itself and against the 450 author-annotated non-malignant cells.

#### Per-seed and ensemble metrics

We evaluated each of three converged *C* = 2,048 seeds individually, every pair of seeds, and all three together, and compared them with the released two-model ensemble. Adding the third seed does not improve the ensemble (validation Pearson 0.714 for three models, 0.718 for the released two), and a single *C* = 2,048 model is a competitive lower-compute and lower-storage alternative.

### 4.11 Statistical analyses

All tests are two-sided unless stated otherwise. No statistical method was used to predetermine sample sizes. On the held-out cell lines, copy-number profiles are compared between methods with Wilcoxon signed-rank tests on the per cell line pseudobulk Pearson and Spearman correlations and loss and gain AUROC (*n* = 59 cell line profiles from 57 distinct lines, 58 for SCEVAN, which gave no output on one MIX-seq line, with the cell line as the unit of replication). P values are adjusted with the Benjamini–Hochberg procedure, and the rank-biserial correlation is the effect size. The cohorts with paired single-cell DNA each hold one cell line or tumor, so no test across units applies. For these, we report the mean per-cell difference in Pearson correlation and in loss and gain AUROC between REFCON and the strongest other method for each metric and setting, on the cells both return, with a 95% percentile confidence interval from 2,000 bootstrap resamples of cells. Per-sample classification results are compared between methods with paired Wilcoxon signed-rank tests over the 94 atlas tumors. These per-sample *p* values are not adjusted for multiple comparisons. Samples in which two methods score the same are left out of that test. F1 is 0 when a class present in the sample is never called. Precision is undefined when a method calls no cell of the class, and such samples are left out of the precision test. Specificity on the normal controls, among the cells each method classifies, is compared with a paired Wilcoxon signed-rank test over the 19 normal-control samples, the PBMC donor and each Tabula Sapiens organ counted once. For the organoids, the agreement within a line is compared over the 10 lines with a paired Wilcoxon signed-rank test, and whether each sample’s profile is closest to the biopsy of its own patient with an exact McNemar test over the 48 samples. Tests were computed with SciPy 1.16.3.

### 4.12 Software versions

All model training, inference, and downstream analyses were run in Python 3.12 with PyTorch 2.10 (CUDA 12.8), NumPy 2.3.5, SciPy 1.16.3, pandas 2.3.3, scikit-learn 1.8.0, statsmodels 0.14.6, scanpy 1.12, anndata 0.12.10, and h5py 3.15.1. Doublets in the training data were flagged with Scrublet as implemented in scanpy 1.12. UMAP embeddings used umap-learn 0.5.11 (0.5.12 for the astrocytoma). Per-cell DNA copy-number ground truth was generated with BWA-MEM 0.7.17 [52], SAMtools 1.9 [53], and AneuFinder 1.34.0 [54] under R 4.4.3. The external copy-number callers were SCEVAN 1.0.3 [19] and CopyKAT 1.1.0 [18] (R 4.4.3), CopyVAE 1.0 [20], inferCNV [14, 17] (run as the Python implementation inferCNVpy 0.6.1 [58]), and Numbat 1.5.2 [39] (R 4.4.1).

### 4.13 Runtime and resource usage

REFCON was developed and benchmarked on NVIDIA GeForce RTX 2080 Ti GPUs. To show how it runs for a typical user, it was also timed on an Apple M4 Pro laptop, on its integrated GPU and on its CPU. Table 2 gives the throughput and memory on each platform, for the default inference and for the forward pass only (Section 4.4). The ensemble members run one after the other, so peak accelerator memory is that of one member. Runtime grows linearly with the number of cells, as each cell is processed independently. Peak GPU memory grew with the batch size, from 0.03 GB at 8 cells per batch to 2.8 GB at 2,048, and was 2.3 GB at 512 including the CUDA context. The output does not depend on the batch size. On the seven Wang samples (the five test patients and two patients with CD45^+^ cells only; Section 2.5; 34,052 cells per sample on average), REFCON took 65 minutes per sample: 34 minutes of inference on one RTX 2080 Ti and 31 minutes of labeling on the CPU. SCEVAN and CopyKAT took 6.8 and 17.3 hours per sample on four CPU cores. The methods ran on different hardware, so these times are given for information only.

## Data availability

All datasets analyzed in this study are publicly available. Training data (Table 1): GSE114461 (MCF7 [37]), GSE160148 (COLO320 [41]), GSE261713 (MDA-MB-231, wellDR-seq [42]), GSE142750 (gastric cancer cell lines, Andor et al. 2020 [43]), GSE157220 / SCP542 (CCLE pan-cancer panel [22]), and Tabula Sapiens healthy tissue [25, 26] (4,200 training and 1,200 validation cells labeled diploid, and 14,400 test cells from 18 organs used as a normal control), from the Tabula Sapiens collection of CZ CELLxGENE Discover [40] (https://cellxgene.cziscience.com/collections/e5f58829-1a66-40b5-a624-9046778e74f5, accessed 24 September 2026); bulk copy number ground truth from the DepMap 24Q4 Public release [50, 51] (https://doi.org/10.25452/figshare.plus.27993248.v1; DepMap portal [59]). **Held-out evaluation cohorts:** GSE157220 / SCP542 (esophageal and mixed-tissue cell lines), figshare https://doi.org/10.6084/m9.figshare.10298696.v3 [60] (MIX-seq pan-cancer and lung panels [27]), GSE144296 (HCT116 and A375, DNTR-seq [11]), GSE185269 (scONE-seq astrocytoma [12]), GSE72056 (Tirosh et al. melanoma [17]), GSE225857 (Wang et al. colorectal cancer with liver metastasis [35]), GSE95435 (Kinchen et al. colonic epithelial cells [29]), GSE151091 (Belote et al. epidermal cells [30]), SCP1644 on the Single Cell Portal (pancreatic cancer biopsies and patient-derived organoids, Raghavan et al. [33]; count matrices from the Curated Cancer Cell Atlas [15, 34]), and the 10x Genomics PBMC 3K healthy-donor demonstration dataset [36]. The atlas tumors are from the Curated Cancer Cell Atlas [15, 34] (https://www.weizmann.ac.il/sites/3CA/), which collects the eight studies listed in Methods (original accessions GSE131928, SCP1644, GSE132465, GSE123139, GSE125449, EGAS00001004987, GSE164690 and GSE159115). Supplementary Data 2 lists the tumor samples used, with each method’s results per tumor. Supplementary Data 3 gives each method’s classification results per patient and per control sample, and for the colorectal patients run per tissue. **Additional unsupervised-clustering cohort**: CellBench (Tian et al. 2019 [31], GSE118767). The processed, model-ready copy number benchmark cohorts (the CCLE and MIX-seq cell lines, HCT116, A375 and the astrocytoma) and the two trained checkpoints of the released ensemble are deposited at Zenodo (https://doi.org/10.5281/zenodo.23534511 [61]).

## Code availability

Model definitions, training code, and the inference, refinement, classification and scoring scripts, with a notebook that reproduces the per-cell results on the bundled cohorts, are available on GitHub (https://github.com/ciceklab/refcon) under an MIT license; the processed data and model checkpoints are archived at Zenodo (https://doi.org/10.5281/zenodo.23534511 [61]).

## Supplementary information

Supplementary Information accompanies this manuscript. It comprises five Supplementary Notes (the organoid clones and their checks; the cases where labeling failed; the per-cell copy number of the DNTR-seq cell lines; external-method configurations, pseudo-reference injection and gene coverage; and the benchmark protocol), 17 Supplementary Tables, seven Supplementary Figures, and three Supplementary Data files, each referenced from the main text at its point of use. The Supplementary Tables give the full per-cell and per cell line metric tables and paired significance tests for the reference-free copy-number benchmark (Section 2.2), the agreement between organoid samples and the organoid clone shares (Section 2.4), the per-sample classification, full-diploid specificity and atlas immune-cell results (Section 2.5), the detection by event size (Discussion), and the ensemble analysis, the ablations (positions, normalization, context length and training additions), the stitching and refinement analyses, the ploidy-relative scoring, the transformer contribution and the robustness to fewer genes and lower sequencing depth (Methods). The Supplementary Figures are the embeddings of the astrocytoma, the cell line embeddings of the clustering benchmark, the checks of the organoid clones, the colorectal patients run per tissue and their expression embeddings, the *τ*_1_ threshold sweep and the precision-recall sensitivity curve. Supplementary Data 1 provides the roster of the training, validation and test splits (cell lines with DepMap identifiers, and the healthy Tabula Sapiens cells by organ and donor), Supplementary Data 2 lists the 94 atlas tumor samples with their study, original accession, cell counts and each method’s per-tumor classification, and Supplementary Data 3 gives each method’s classification results per patient and per control sample with their cell counts, and for the colorectal patients run per tissue.

## Acknowledgements.

The authors acknowledge the computational resources of the Bilkent University Department of Computer Engineering and thank Mehmet Can Bastem for their assistance with exploratory experiments.

## Ethics approval

Not applicable. All datasets are publicly available and previously published.

## Author contributions

M.M.G. conceived and designed the method, implemented the model and analysis pipelines, performed the experiments and analyses, and wrote the manuscript. A.E.Ç . supervised the study and revised the manuscript. Both authors read and approved the final manuscript.

## References

[1] Steele, C. D. et al. Signatures of copy number alterations in human cancer. Nature 606, 984–991 (2022).

[2] Lukow, D. A. & Sheltzer, J. M. Chromosomal instability and aneuploidy as causes of cancer drug resistance. Trends in Cancer 8, 43–53 (2022).

[3] Drews, R. M. et al. A pan-cancer compendium of chromosomal instability. Nature 606, 976–983 (2022).

[4] Watkins, T. B. K. et al. Pervasive chromosomal instability and karyotype order in tumour evolution. Nature 587, 126–132 (2020).

[5] McGranahan, N. & Swanton, C. Clonal heterogeneity and tumor evolution: Past, present, and the future. Cell 168, 613–628 (2017).

[6] Jamal-Hanjani, M. et al. Tracking the evolution of non–small-cell lung cancer. New England Journal of Medicine 376, 2109–2121 (2017).

[7] Neftel, C. et al. An integrative model of cellular states, plasticity, and genetics for glioblastoma. Cell 178, 835–849.e21 (2019).

[8] Funnell, T. et al. Single-cell genomic variation induced by mutational processes in cancer. Nature 612, 106–115 (2022).

[9] Otonicar, J., et al. HIPSD&R-seq enables scalable genomic copy number and transcriptome profiling. Genome Biology 25, 316 (2024).

[10] Shao, D. D. et al. Advances in single-cell DNA sequencing enable insights into human somatic mosaicism. Nature Reviews Genetics 26, 761–774 (2025).

[11] Zachariadis, V., Cheng, H., Andrews, N. & Enge, M. A highly scalable method for joint whole-genome sequencing and gene-expression profiling of single cells. Molecular Cell 80, 541–553.e5 (2020).

[12] Yu, L. et al. scONE-seq: A single-cell multi-omics method enables simultaneous dissection of phenotype and genotype heterogeneity from frozen tumors. Science Advances 9, eabp8901 (2023).

[13] Vandereyken, K., Sifrim, A., Thienpont, B. & Voet, T. Methods and applications for single-cell and spatial multi-omics. Nature Reviews Genetics 24, 494–515 (2023).

[14] Patel, A. P. et al. Single-cell RNA-seq highlights intratumoral heterogeneity in primary glioblastoma. Science 344, 1396–1401 (2014).

[15] Gavish, A. et al. Hallmarks of transcriptional intratumour heterogeneity across a thousand tumours. Nature 618, 598–606 (2023).

[16] Schmid, K. T. et al. Benchmarking scRNA-seq copy number variation callers. Nature Communications 16, 8777 (2025).

[17] Tirosh, I. et al. Dissecting the multicellular ecosystem of metastatic melanoma by single-cell RNA-seq. Science 352, 189–196 (2016).

[18] Gao, R. et al. Delineating copy number and clonal substructure in human tumors from single-cell transcriptomes. Nature Biotechnology 39, 599–608 (2021).

[19] De Falco, A., Caruso, F., Su, X.-D., Iavarone, A. & Ceccarelli, M. A variational algorithm to detect the clonal copy number substructure of tumors from scRNA-seq data. Nature Communications 14, 1074 (2023).

[20] Kurt, S. et al. CopyVAE: a variational autoencoder-based approach for copy number variation inference using single-cell transcriptomics. Bioinformatics 40, btae284 (2024).

[21] Oketch, D. J. A., Giulietti, M. & Piva, F. A comparison of tools that identify tumor cells by inferring copy number variations from single-cell experiments in pancreatic ductal adenocarcinoma. Biomedicines 12, 1759 (2024).

[22] Kinker, G. S. et al. Pan-cancer single-cell RNA-seq identifies recurring programs of cellular heterogeneity. Nature Genetics 52, 1208–1218 (2020).

[23] Wang, R. et al. Systematic evaluation of colorectal cancer organoid system by single-cell RNA-Seq analysis. Genome Biology 23, 106 (2022).

[24] Vaswani, A. et al. Attention is all you need. Advances in Neural Information Processing Systems 30, 5998–6008 (2017).

[25] Tabula Sapiens Consortium et al. The Tabula Sapiens: A multiple-organ, single-cell transcriptomic atlas of humans. Science 376, eabl4896 (2022).

[26] Jones, R. C. et al. Tabula Sapiens 2.0: A comprehensive transcriptomic atlas of human cell types. Cell (2026). URL 10.1016/j.cell.2026.08.010.

[27] McFarland, J. M. et al. Multiplexed single-cell transcriptional response profiling to define cancer vulnerabilities and therapeutic mechanism of action. Nature Communications 11, 4296 (2020).

[28] Zheng, G. X. Y. et al. Massively parallel digital transcriptional profiling of single cells. Nature Communications 8, 14049 (2017).

[29] Kinchen, J. et al. Structural remodeling of the human colonic mesenchyme in inflammatory bowel disease. Cell 175, 372–386.e17 (2018).

[30] Belote, R. L. et al. Human melanocyte development and melanoma dedifferentiation at single-cell resolution. Nature Cell Biology 23, 1035–1047 (2021).

[31] Tian, L. et al. Benchmarking single cell RNA-sequencing analysis pipelines using mixture control experiments. Nature Methods 16, 479–487 (2019).

[32] McInnes, L., Healy, J. & Melville, J. UMAP: Uniform manifold approximation and projection for dimension reduction. *arXiv preprint arXiv:1802.03426* (2018).

[33] Raghavan, S. et al. Microenvironment drives cell state, plasticity, and drug response in pancreatic cancer. Cell 184, 6119–6137.e26 (2021).

[34] Tyler, M. et al. The Curated Cancer Cell Atlas provides a comprehensive characterization of tumors at single-cell resolution. Nature Cancer 6, 1088–1101 (2025).

[35] Wang, F. et al. Single-cell and spatial transcriptome analysis reveals the cellular heterogeneity of liver metastatic colorectal cancer. Science Advances 9, eadf5464 (2023).

36. 10x Genomics. 3k PBMCs from a Healthy Donor. Public scRNA-seq dataset, Chromium Single Cell Gene Expression (2016). URL https://www.10xgenomics.com/datasets/3-k-pbm-cs-from-a-healthy-donor-1-standard-1-1-0. Dataset.

[37] Ben-David, U. et al. Genetic and transcriptional evolution alters cancer cell line drug response. Nature 560, 325–330 (2018).

[38] Bolhaqueiro, A. C. F. et al. Ongoing chromosomal instability and karyotype evolution in human colorectal cancer organoids. Nature Genetics 51, 824–834 (2019).

[39] Gao, T. et al. Haplotype-aware analysis of somatic copy number variations from single-cell transcriptomes. Nature Biotechnology 41, 417–426 (2023).

40. CZI Cell Science Program et al. CZ CELLxGENE Discover: a single-cell data platform for scalable exploration, analysis and modeling of aggregated data. Nucleic Acids Research 53, D886–D900 (2025).

[41] Hung, K. L. et al. ecDNA hubs drive cooperative intermolecular oncogene expression. Nature 600, 731–736 (2021).

[42] Wang, K. et al. Coalescing single-cell genomes and transcriptomes to decode breast cancer progression. Cell 188, 6355–6369.e16 (2025).

43. Andor, N., et al. Joint single cell DNA-seq and RNA-seq of gastric cancer cell lines reveals rules of in vitro evolution. NAR Genomics and Bioinformatics 2, lqaa016 (2020).

[44] Lee, H.-O. et al. Lineage-dependent gene expression programs influence the immune landscape of colorectal cancer. Nature Genetics 52, 594–603 (2020).

[45] Li, H. et al. Dysfunctional CD8 T cells form a proliferative, dynamically regulated compartment within human melanoma. Cell 176, 775–789.e18 (2019).

[46] Ma, L. et al. Tumor cell biodiversity drives microenvironmental reprogramming in liver cancer. Cancer Cell 36, 418–430.e6 (2019).

[47] Olbrecht, S. et al. High-grade serous tubo-ovarian cancer refined with single-cell RNA sequencing: specific cell subtypes influence survival and determine molecular subtype classification. Genome Medicine 13, 111 (2021).

[48] Kürten, C. H. L., et al. Investigating immune and non-immune cell interactions in head and neck tumors by single-cell RNA sequencing. Nature Communications 12, 7338 (2021).

[49] Zhang, Y. et al. Single-cell analyses of renal cell cancers reveal insights into tumor microenvironment, cell of origin, and therapy response. Proceedings of the National Academy of Sciences 118, e2103240118 (2021).

[50] DepMap, Broad. DepMap 24Q4 Public (2024). URL 10.25452/figshare.plus.27993248.v1. Figshare+. Dataset.

[51] Arafeh, R., Shibue, T., Dempster, J. M., Hahn, W. C. & Vazquez, F. The present and future of the Cancer Dependency Map. Nature Reviews Cancer 25, 59–73 (2025).

52. Li, H. Aligning sequence reads, clone sequences and assembly contigs with BWA-MEM. arXiv preprint arXiv:1303.3997 (2013).

[53] Danecek, P. et al. Twelve years of SAMtools and BCFtools. GigaScience 10, giab008 (2021).

[54] Bakker, B. et al. Single-cell sequencing reveals karyotype heterogeneity in murine and human malignancies. Genome Biology 17, 115 (2016).

55. Curated Cancer Cell Atlas. Curated Cancer Cell Atlas: Methods. https://www.weizmann.ac.il/sites/3CA/methods (2026). Weizmann Institute of Science. Accessed 15 October 2026.

[56] Picelli, S. et al. Smart-seq2 for sensitive full-length transcriptome profiling in single cells. Nature Methods 10, 1096–1098 (2013).

[57] Su, J. et al. RoFormer: Enhanced Transformer with rotary position embedding. Neurocomputing 568, 127063 (2024).

[58] Sturm, G. inferCNVpy: a scalable Python library to infer copy number variation events from single-cell transcriptomics data. Python package, version 0.6.1 (2025). URL https://github.com/icbi-lab/infercnvpy.

59. DepMap, Broad. DepMap: The Cancer Dependency Map Project at Broad Institute. https://depmap.org/portal. Accessed 4 October 2026.

[60] Cancer Data Science. MIX-seq data (2019). URL 10.6084/m9.figshare.10298696.v3. Figshare. Dataset.

[61] Gençtürk, M. M. & Çiçek, A. E. REFCON: model checkpoints and copy number test cohorts (2026). URL 10.5281/zenodo.23534511. Zenodo. Dataset.

