## Supplementary Information for "Reference-free and robust copy number inference in single-cell tumor transcriptomes"

Citation numbers refer to the reference list of the main article.

### Supplementary Figures

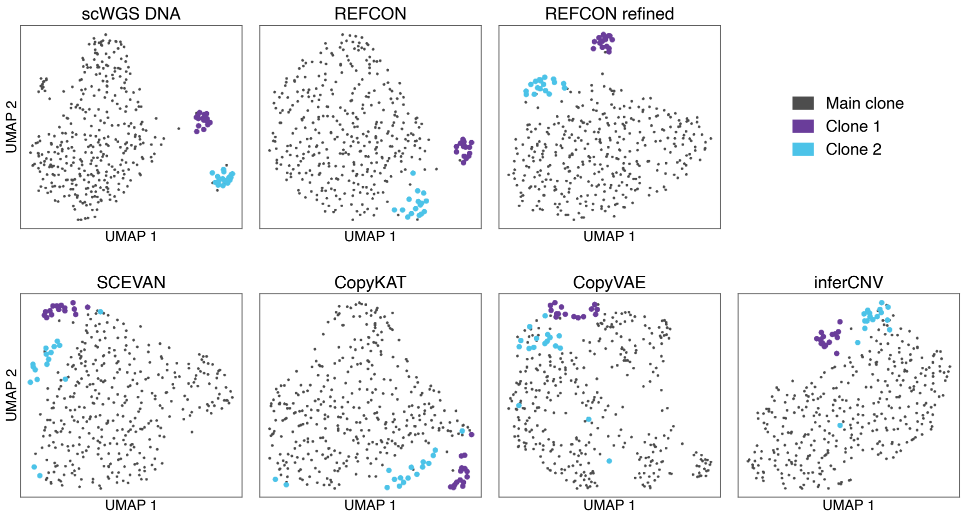


**Supplementary Figure 1. Embeddings of the scONE-seq astrocytoma.** UMAP embeddings of the 390 malignant cells from the paired scWGS, REFCON, REFCON refined against the 63 confident-normal cells it detects, SCEVAN, CopyKAT, CopyVAE and inferCNV. All profiles come from runs on the whole sample (840 cells, with no cells marked as normal). Genes a method never calls are dropped, missing values are set to the gene mean over the 390 cells, and the top 20 principal components (not standardized) are embedded with UMAP [32] (30 neighbors, minimum distance 0.3, random seed 42).

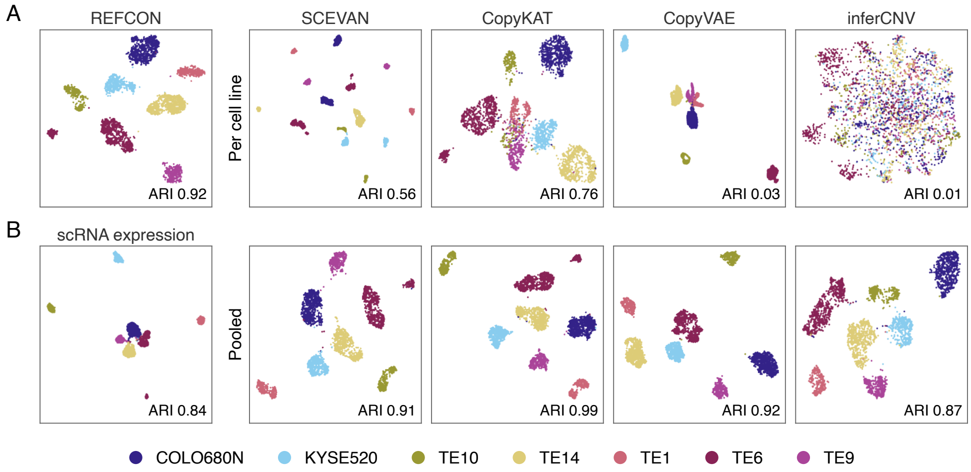

**Supplementary Figure 2. Embeddings of the seven esophageal CCLE lines, per cell line and pooled.** UMAP embeddings of the clustering input of Figure 4B, with the ARI of each embedding. **A** Each cell line run on its own: REFCON, SCEVAN, CopyKAT, CopyVAE and inferCNV. **B** Log-transformed expression without copy number inference, and SCEVAN, CopyKAT, CopyVAE and inferCNV with all seven lines run together.

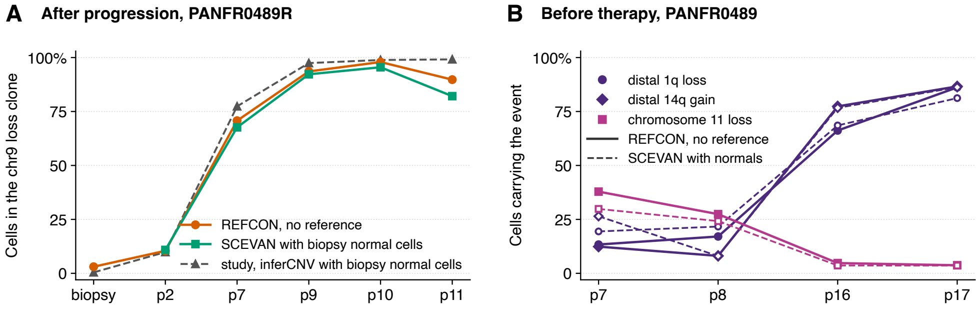


**Supplementary Figure 3. Independent checks of the organoid clones.** **A** PANFR0489R: share of cells in the chromosome 9 loss clone per sample, from REFCON without a reference, from SCEVAN with the biopsy's normal cells as reference, and from the study's inferCNV clones D, E and F (Raghavan et al. [33], Figure S5J; Supplementary Table 6). For both methods, each cell's chromosome 9 value is split into two groups (Supplementary Note 1). **B** PANFR0489: share of cells carrying each clonal event per passage, from REFCON without a reference and from SCEVAN with the biopsy's normal cells. Each segment is split into two groups of cells separately for each method.

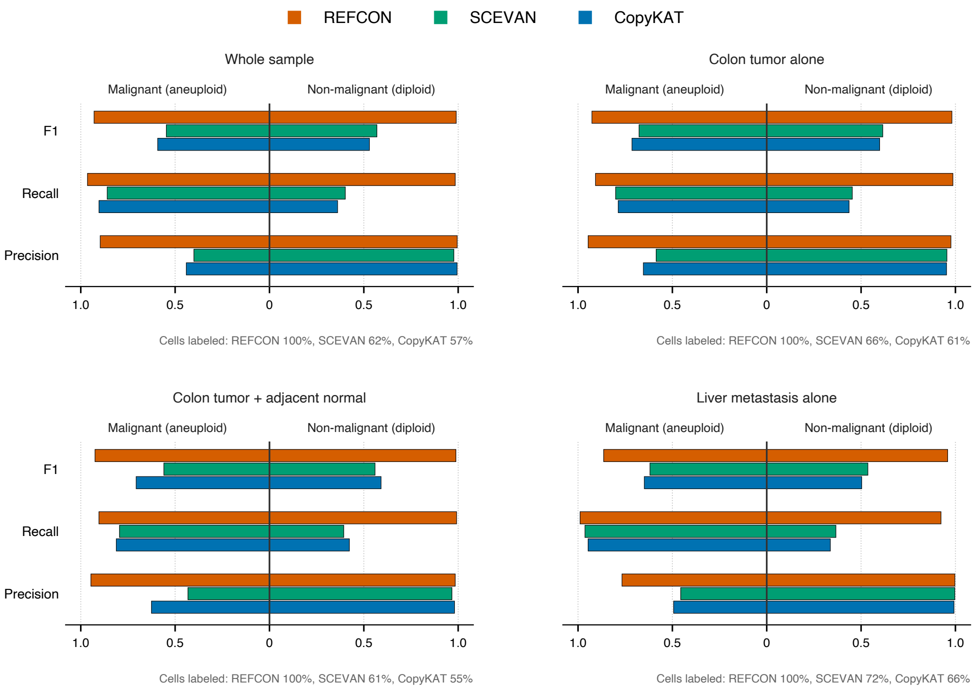


**Supplementary Figure 4. Classification of the five colorectal patients by tissue.** Aneuploid- and diploid-class precision, recall and F1 of REFCON, SCEVAN and CopyKAT on the 5 colorectal patients of Wang et al. [35], pooled over cells. Each method was run on four inputs per patient: all tissues together (as in Section 2.5), the colon tumor alone, the colon tumor with the adjacent normal colon, and the liver metastasis alone.

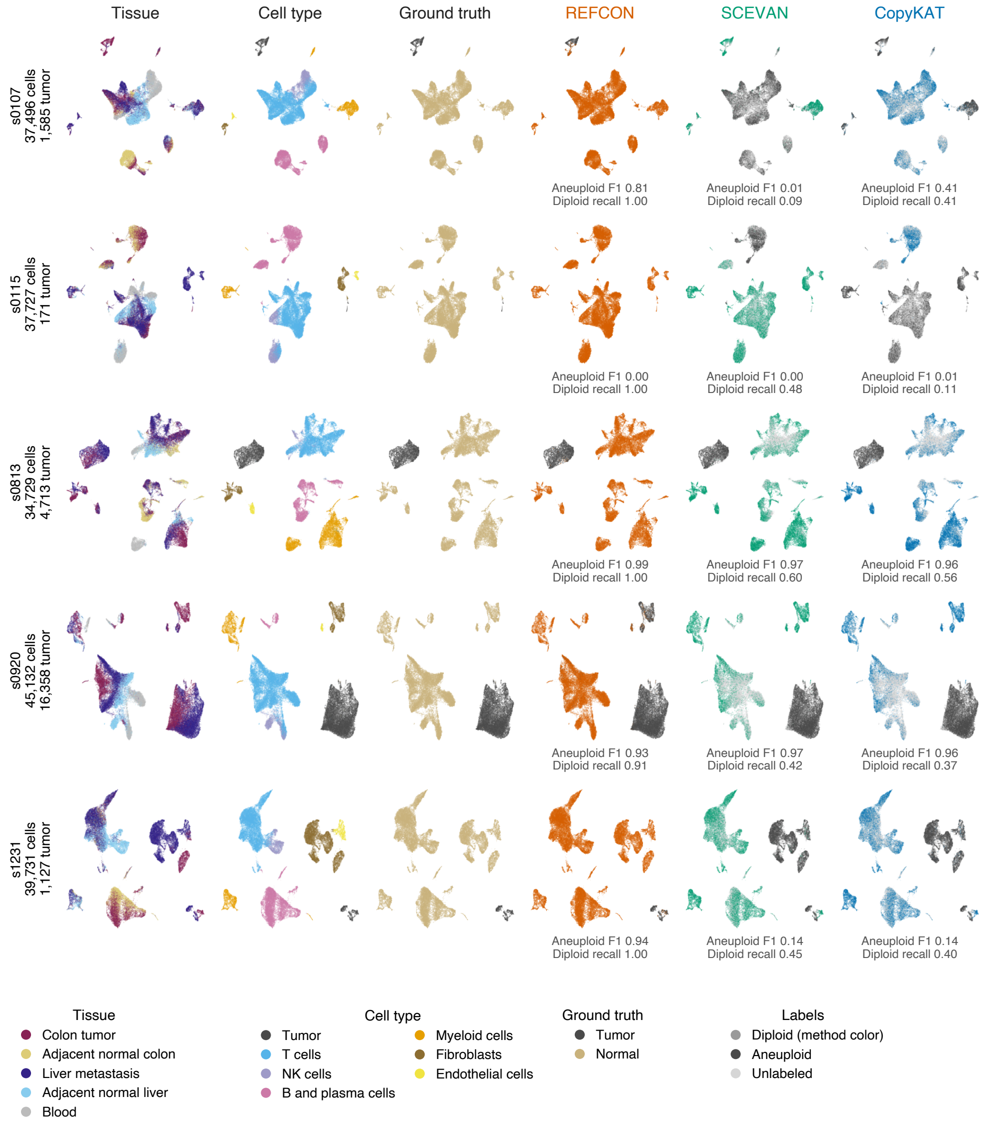


**Supplementary Figure 5. Expression embeddings of the five colorectal patients.** Expression UMAP of each colorectal patient with all its tissues, colored by tissue, author cell type, ground truth and the labels of REFCON, SCEVAN and CopyKAT from the whole-sample runs. The UMAP uses 2,000 highly variable genes of log-normalized counts, 30 principal components and 15 neighbors.

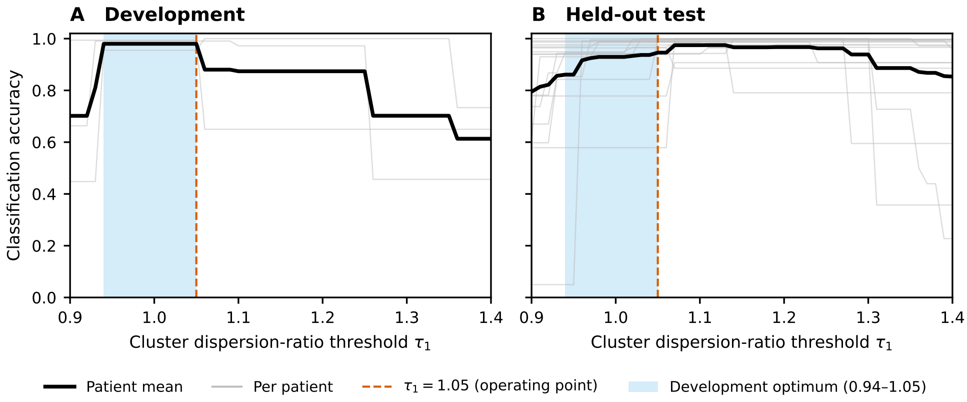


**Supplementary Figure 6. Per-patient classification accuracy across the cluster dispersion-ratio threshold $\tau_1$.** Aneuploid versus diploid classification accuracy of REFCON per patient as a function of $\tau_1$ (Methods, Section 4.7). **A** The three development patients, three melanoma patients of Tirosh et al. [17]. **B** The 17 held-out test patients, 12 further melanoma patients of the same study and 5 colorectal patients of Wang et al. [35]. Accuracy is the fraction of a patient's cells whose label matches the author annotation. The released operating point is $\tau_1 = 1.05$.

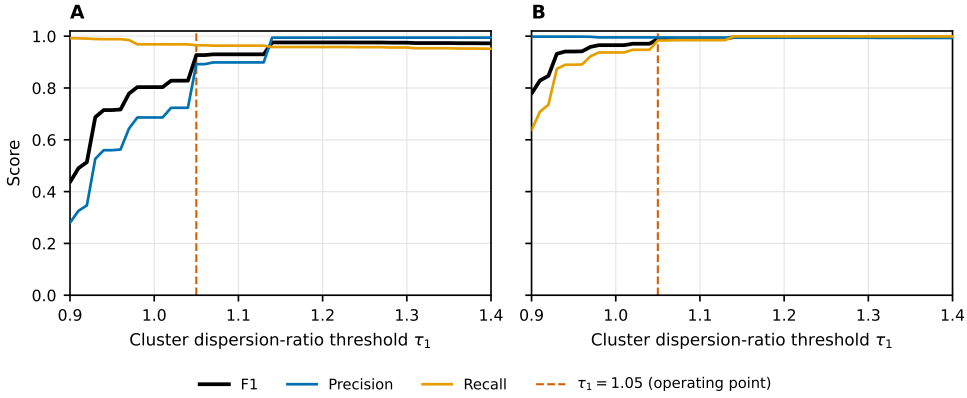


**Supplementary Figure 7. Precision, recall and F1 of REFCON's per-cell classification across the cluster dispersion-ratio threshold $\tau_1$.** Aneuploid-class (**A**) and diploid-class (**B**) precision, recall and F1, pooled over the cells of the 17 held-out patients (12 melanoma patients of Tirosh et al. [17] and 5 colorectal patients of Wang et al. [35]; 197,100 evaluable cells). The released operating point is $\tau_1 = 1.05$ (Methods, Section 4.7).

### Supplementary Tables

**Supplementary Table 1** Per cell line copy number metrics on the 59 held-out cell line profiles. REFCON was run on each line alone. SCEVAN, CopyKAT, CopyVAE and inferCNV were run on each line alone and on all lines of a cohort together (pooled). No method received a reference. Each metric compares the pseudobulk prediction of a line with its bulk DNA copy number and is averaged over the lines of the cohort

| Method | Pearson | Spearman | $AUROC_L$ | $AUROC_G$ | $AUPRC_L$ | $AUPRC_G$ |
|---|---|---|---|---|---|---|
| *Esophageal cohort (7 cell lines, 2,543 cells)* | | | | | | |
| REFCON | **0.553** | **0.639** | **0.711** | **0.819** | **0.132** | **0.937** |
| SCEVAN | 0.257 | 0.233 | 0.590 | 0.594 | 0.042 | 0.824 |
| CopyKAT | 0.105 | 0.115 | 0.552 | 0.536 | 0.026 | 0.797 |
| CopyVAE | −0.033 | −0.063 | 0.349 | 0.443 | 0.011 | 0.746 |
| inferCNV | −0.032 | −0.065 | 0.445 | 0.440 | 0.011 | 0.786 |
| Pooled (all lines of the cohort run together) | | | | | | |
| SCEVAN | 0.283 | 0.277 | 0.637 | 0.640 | 0.064 | 0.842 |
| CopyKAT | 0.278 | 0.295 | 0.589 | 0.623 | 0.051 | 0.845 |
| CopyVAE | 0.289 | 0.281 | 0.565 | 0.644 | 0.030 | 0.833 |
| inferCNV | 0.414 | 0.479 | 0.635 | 0.733 | 0.036 | 0.914 |
| *Mixed-tissue cohort (3 cell lines, 1,512 cells)* | | | | | | |
| REFCON | **0.467** | **0.476** | **0.721** | 0.764 | **0.057** | 0.623 |
| SCEVAN | 0.161 | 0.129 | 0.510 | 0.557 | 0.016 | 0.488 |
| CopyKAT | 0.067 | 0.055 | 0.548 | 0.548 | 0.023 | 0.534 |
| CopyVAE | −0.009 | −0.070 | 0.484 | 0.504 | 0.021 | 0.440 |
| inferCNV | −0.058 | −0.066 | 0.572 | 0.530 | 0.028 | 0.457 |
| Pooled (all lines of the cohort run together) | | | | | | |
| SCEVAN | 0.163 | 0.151 | 0.549 | 0.568 | 0.030 | 0.489 |
| CopyKAT | 0.179 | 0.141 | 0.541 | 0.592 | 0.019 | 0.501 |
| CopyVAE | 0.055 | 0.029 | 0.544 | 0.553 | 0.049 | 0.492 |
| inferCNV | 0.407 | 0.448 | 0.693 | **0.776** | 0.034 | **0.651** |
| *MIX-seq cohort (49 pan-cancer cell lines, 3,055 cells)* | | | | | | |
| REFCON | **0.566** | **0.618** | **0.775** | **0.843** | **0.191** | **0.918** |
| SCEVAN | 0.284 | 0.287 | 0.640 | 0.659 | 0.052 | 0.810 |
| CopyKAT | 0.039 | 0.050 | 0.505 | 0.520 | 0.021 | 0.740 |
| CopyVAE | −0.013 | −0.024 | 0.451 | 0.473 | 0.021 | 0.720 |
| inferCNV | −0.072 | −0.093 | 0.410 | 0.421 | 0.013 | 0.712 |
| Pooled (all lines of the cohort run together) | | | | | | |
| SCEVAN | 0.306 | 0.321 | 0.694 | 0.686 | 0.092 | 0.819 |
| CopyKAT | 0.296 | 0.310 | 0.591 | 0.655 | 0.041 | 0.825 |
| CopyVAE | 0.177 | 0.202 | 0.591 | 0.609 | 0.032 | 0.782 |
| inferCNV | 0.520 | 0.571 | 0.654 | 0.781 | 0.114 | 0.904 |

Cohorts: 7 esophageal and 3 mixed-tissue lines (CL34, NCIH460, SKMEL5) from Kinker et al. [22], and 49 MIX-seq lines from McFarland et al. [27]. Truth: bulk DepMap 24Q4 copy number [50, 51]. Loss: copy number below 1.5. Gain: above 2.5. Subscripts L and G denote loss and gain. AUPRC is the average precision. Genes a method does not call are filled with the median prediction (Methods, Section 4.9). In the pooled rows, each line is scored on the genes it expresses. The MIX-seq pool holds the 49 scored lines. Best per column within each cohort, over both settings, in bold.

**Supplementary Table 2** Paired comparison of REFCON against each other method. Held-out cell lines: per cell line pseudobulk Pearson and Spearman correlation and loss and gain AUROC against bulk copy number (per cell line setting of Supplementary Table 1), compared over lines. Cohorts with paired single-cell DNA: per-cell Pearson correlation and loss and gain AUROC against the paired DNA, compared over cells with the strongest other method of each metric, with every method run without a reference and with reference cells given to the other methods

| Comparison | Metric | $n$ | $\Delta$ | Effect size | $p_{\text{adj}}$ / 95% CI |
|---|---|---|---|---|---|
| *Held-out cell lines (59 cell line profiles from 57 distinct lines; unit = cell line)* | | | | | |
| vs. SCEVAN | Pearson | 58 | 0.30 | 1.00 | $5.4 \times 10^{-11}$ |
| vs. CopyKAT | Pearson | 59 | 0.55 | 1.00 | $5.4 \times 10^{-11}$ |
| vs. CopyVAE | Pearson | 59 | 0.60 | 1.00 | $5.4 \times 10^{-11}$ |
| vs. inferCNV | Pearson | 59 | 0.65 | 1.00 | $5.4 \times 10^{-11}$ |
| vs. SCEVAN | Spearman | 58 | 0.37 | 1.00 | $5.4 \times 10^{-11}$ |
| vs. CopyKAT | Spearman | 59 | 0.58 | 1.00 | $5.4 \times 10^{-11}$ |
| vs. CopyVAE | Spearman | 59 | 0.67 | 1.00 | $5.4 \times 10^{-11}$ |
| vs. inferCNV | Spearman | 59 | 0.72 | 1.00 | $5.4 \times 10^{-11}$ |
| vs. SCEVAN | loss AUROC | 54 | 0.14 | 0.96 | $6.9 \times 10^{-10}$ |
| vs. CopyKAT | loss AUROC | 55 | 0.24 | 0.98 | $2.7 \times 10^{-10}$ |
| vs. CopyVAE | loss AUROC | 55 | 0.33 | 0.98 | $2.6 \times 10^{-10}$ |
| vs. inferCNV | loss AUROC | 55 | 0.38 | 0.98 | $2.6 \times 10^{-10}$ |
| vs. SCEVAN | gain AUROC | 57 | 0.20 | 0.99 | $1.1 \times 10^{-10}$ |
| vs. CopyKAT | gain AUROC | 58 | 0.34 | 1.00 | $5.4 \times 10^{-11}$ |
| vs. CopyVAE | gain AUROC | 58 | 0.37 | 1.00 | $5.4 \times 10^{-11}$ |
| vs. inferCNV | gain AUROC | 58 | 0.40 | 1.00 | $5.4 \times 10^{-11}$ |
| *Per-cell paired-DNA cohorts, every method without a reference (single line or tumor)* | | | | | |
| A375 vs. SCEVAN | Pearson | 60 | +0.23 | n/a | [+0.22, +0.25] |
| A375 vs. SCEVAN | loss AUROC | 60 | +0.10 | n/a | [+0.09, +0.11] |
| A375 vs. SCEVAN | gain AUROC | 60 | +0.14 | n/a | [+0.14, +0.15] |
| HCT116 vs. inferCNV | Pearson | 1,356 | +0.13 | n/a | [+0.12, +0.14] |
| HCT116 vs. inferCNV | loss AUROC | 1,465 | +0.13 | n/a | [+0.13, +0.14] |
| HCT116 vs. SCEVAN | gain AUROC | 1,458 | +0.22 | n/a | [+0.21, +0.22] |
| Astrocytoma vs. CopyKAT | Pearson | 390 | +0.13 | n/a | [+0.13, +0.14] |
| Astrocytoma vs. CopyKAT | loss AUROC | 390 | +0.08 | n/a | [+0.08, +0.08] |
| Astrocytoma vs. CopyKAT | gain AUROC | 390 | +0.08 | n/a | [+0.08, +0.08] |
| *Per-cell paired-DNA cohorts, reference cells given to the other methods* | | | | | |
| A375 vs. inferCNV | Pearson | 64 | +0.11 | n/a | [+0.10, +0.13] |
| A375 vs. SCEVAN | loss AUROC | 60 | +0.11 | n/a | [+0.10, +0.12] |
| A375 vs. SCEVAN | gain AUROC | 60 | +0.10 | n/a | [+0.09, +0.11] |
| HCT116 vs. inferCNV | Pearson | 1,467 | +0.11 | n/a | [+0.11, +0.12] |
| HCT116 vs. SCEVAN | loss AUROC | 1,460 | +0.14 | n/a | [+0.14, +0.15] |
| HCT116 vs. CopyKAT | gain AUROC | 1,456 | +0.18 | n/a | [+0.17, +0.18] |
| Astrocytoma, unrefined vs. inferCNV | Pearson | 390 | -0.06 | n/a | [-0.06, -0.05] |
| Astrocytoma, unrefined vs. CopyVAE | loss AUROC | 390 | +0.04 | n/a | [+0.03, +0.04] |
| Astrocytoma, unrefined vs. SCEVAN | gain AUROC | 390 | +0.03 | n/a | [+0.03, +0.03] |
| Astrocytoma, refined vs. inferCNV | Pearson | 390 | +0.10 | n/a | [+0.10, +0.10] |
| Astrocytoma, refined vs. CopyVAE | loss AUROC | 390 | +0.12 | n/a | [+0.12, +0.13] |
| Astrocytoma, refined vs. SCEVAN | gain AUROC | 390 | +0.09 | n/a | [+0.09, +0.10] |

Held-out cell lines: tests as in Methods, Section 4.11, with Benjamini–Hochberg (BH) adjusted $p$ values. $\Delta$ is the median per cell line difference, REFCON minus the other method. Effect size is the rank-biserial correlation. CopyKAT's 15 MIX-seq lines that stopped with an error are scored from the rerun with its bug fixed (Supplementary Note 4). COLO680N and TE6 are in both the CCLE esophageal panel and MIX-seq. Loss AUROC is defined for the lines with loss genes (54 or 55). With the two profiles of COLO680N and TE6 averaged, every Pearson and Spearman $p_{\text{adj}}$ is at most $7.6 \times 10^{-11}$. Cohorts with paired single-cell DNA: $\Delta$ is the mean per-cell difference on the cells both methods return, and the last column gives its 95% bootstrap confidence interval over cells.

The compared method is the other method with the highest mean of that metric in that setting (Supplementary Table 4). Astrocytoma rows with references compare REFCON, unrefined or refined against its predictions for the 450 non-malignant cells, with the methods given these cells.

**Supplementary Table 3** False-positive copy number calls on copy-neutral genes at matched sensitivity. Every method was run without a reference on HCT116, A375 and the 49 MIX-seq lines, each line alone. For each cell and event type, genes are ranked by the method's prediction and the cutoff is set where half of the cell's true events are called. The table gives the fraction of copy-neutral genes called at that cutoff, and the fraction of genes each method predicts (call)

| | HCT116 | | | A375 | | | MIX-seq | | |
|---|---|---|---|---|---|---|---|---|---|
| Method | gain | loss | call | gain | loss | call | gain | loss | call |
| REFCON | **0.11** | 0.23 | 1.00 | **0.18** | **0.20** | 1.00 | **0.16** | **0.14** | 0.99 |
| inferCNV | **0.11** | **0.15** | 1.00 | 0.30 | 0.29 | 1.00 | 0.49 | 0.49 | 0.99 |
| CopyKAT | 0.28 | 0.25 | 0.47 | 0.31 | 0.34 | 0.56 | 0.46 | 0.45 | 0.60 |
| SCEVAN | 0.27 | 0.27 | 0.50 | 0.24 | 0.32 | 0.60 | 0.31 | 0.30 | 0.63 |
| CopyVAE | 0.43 | 0.46 | 0.95 | 0.52 | 0.55 | 0.94 | 0.50 | 0.53 | 0.94 |

Gene classes are relative to each cell's median DNA copy number: copy-neutral within $[0.90, 1.10]$, gain above 1.15 and loss below 0.85. Genes in $(1.10, 1.15]$ and $[0.85, 0.90)$ are excluded. MIX-seq: a cell's copy number is the bulk DepMap copy number of its line. G401 has no gene above the gain cutoff, so MIX-seq is scored on 48 lines. Gain and loss are scored separately. Each method is scored only on the genes it predicts, and genes it does not predict are dropped. Call: the fraction of evaluable genes with a finite prediction. Lowest false-positive rate per gain and loss column in bold. The call column is not ranked.

**Supplementary Table 4** Per-cell copy number metrics on the three cohorts with paired single-cell DNA. Each method was run without a reference and with a normal reference panel. On the astrocytoma, each method was also run on the whole sample with no cells marked as normal.

Each cell's prediction is compared with the copy number of the same cell from single-cell DNA sequencing, and metrics are averaged over cells

| Method | Pearson | Spearman | $\text{AUROC}_\text{L}$ | $\text{AUROC}_\text{G}$ | $\text{AUPRC}_\text{L}$ | $\text{AUPRC}_\text{G}$ |
|---|---|---|---|---|---|---|
| *HCT116 (DNTR-seq,* 1,467 *cells;* $\pi_{loss} = 10.2\%$*,* $\pi_{gain} = 13.9\%$*)* | | | | | | |
| Reference-free | | | | | | |
| REFCON | **0.429** | **0.389** | **0.737** | **0.835** | **0.315** | **0.518** |
| SCEVAN | 0.183 | 0.160 | 0.598 | 0.619 | 0.161 | 0.207 |
| CopyKAT | 0.195 | 0.148 | 0.589 | 0.598 | 0.208 | 0.225 |
| CopyVAE | 0.058 | 0.068 | 0.523 | 0.553 | 0.152 | 0.169 |
| inferCNV | 0.306 | 0.298 | 0.604 | 0.571 | 0.262 | 0.235 |
| Numbat | 0.106 | 0.124 | 0.502 | 0.520 | 0.101 | 0.177 |
| Reference-included (Kinchen colon refs, $n = 96$) | | | | | | |
| SCEVAN + refs | 0.225 | 0.177 | 0.593 | 0.638 | 0.209 | 0.257 |
| CopyKAT + refs | 0.232 | 0.195 | 0.591 | 0.660 | 0.190 | 0.261 |
| CopyVAE + refs | 0.108 | 0.110 | 0.559 | 0.595 | 0.157 | 0.178 |
| inferCNV + refs | 0.315 | 0.305 | 0.569 | 0.645 | 0.163 | 0.320 |
| *A375 (DNTR-seq,* 64 *cells;* $\pi_{loss} = 6.8\%$*,* $\pi_{gain} = 64.1\%$*)* | | | | | | |
| Reference-free | | | | | | |
| REFCON | **0.470** | **0.479** | **0.734** | **0.759** | **0.144** | **0.828** |
| SCEVAN | 0.263 | 0.252 | 0.650 | 0.632 | 0.092 | 0.742 |
| CopyKAT | 0.184 | 0.177 | 0.573 | 0.594 | 0.077 | 0.714 |
| inferCNV | 0.063 | 0.069 | 0.502 | 0.515 | 0.078 | 0.651 |
| CopyVAE | −0.092 | −0.073 | 0.432 | 0.463 | 0.066 | 0.612 |
| Numbat | 0.442 | 0.394 | 0.527 | 0.578 | 0.075 | 0.710 |
| Reference-included (Belote melanocyte refs, $n = 200$) | | | | | | |
| SCEVAN + refs | 0.333 | 0.322 | 0.645 | 0.676 | 0.108 | 0.769 |
| CopyKAT + refs | 0.320 | 0.300 | 0.631 | 0.659 | 0.103 | 0.759 |
| CopyVAE + refs | — | — | — | — | — | — |
| inferCNV + refs | 0.356 | 0.336 | 0.602 | 0.615 | 0.108 | 0.706 |
| *scONE-seq astrocytoma (*390 *malignant cells)* | | | | | | |
| Reference-free | | | | | | |
| REFCON | **0.360** | **0.345** | **0.694** | **0.657** | **0.467** | **0.428** |
| CopyKAT | 0.226 | 0.207 | 0.616 | 0.579 | 0.378 | 0.332 |
| SCEVAN | 0.149 | 0.131 | 0.558 | 0.549 | 0.303 | 0.312 |
| inferCNV | 0.159 | 0.150 | 0.541 | 0.541 | 0.289 | 0.310 |
| CopyVAE | −0.005 | 0.006 | 0.504 | 0.502 | 0.266 | 0.278 |
| Whole sample, normal cells unannotated (840 cells run, 390 scored) | | | | | | |
| REFCON refined | **0.510** | **0.483** | **0.777** | **0.720** | **0.602** | **0.505** |
| SCEVAN | 0.338 | 0.272 | 0.635 | 0.615 | 0.406 | 0.385 |
| CopyKAT | 0.327 | 0.274 | 0.639 | 0.614 | 0.414 | 0.376 |
| CopyVAE | 0.201 | 0.244 | 0.656 | 0.599 | 0.383 | 0.346 |
| inferCNV | 0.306 | 0.274 | 0.571 | 0.569 | 0.324 | 0.343 |
| Reference-included (450 non-malignant cells) | | | | | | |
| REFCON refined | **0.517** | **0.486** | **0.779** | **0.721** | **0.605** | **0.508** |
| SCEVAN + refs | 0.361 | 0.293 | 0.651 | 0.627 | 0.427 | 0.401 |
| CopyKAT + refs | 0.336 | 0.277 | 0.641 | 0.616 | 0.417 | 0.382 |
| CopyVAE + refs | 0.201 | 0.244 | 0.656 | 0.599 | 0.383 | 0.346 |
| inferCNV + refs | 0.416 | 0.382 | 0.626 | 0.605 | 0.395 | 0.378 |

Loss: copy number below 1.5. Gain: above 2.5. Subscripts L and G denote loss and gain, and $\pi$ is the share of loss or gain genes. Genes a method does not call are filled with the median prediction (Methods, Section 4.9). Means are over the cells a method returns: on HCT116, 1,462 for SCEVAN, 1,460 for CopyKAT and, without a reference, 1,356 for inferCNV; on A375, 60 of 64 for SCEVAN and CopyKAT. HCT116: 1,467 of the 1,468 paired cells have DNA copy number. A375: 64 cells with finite AneuFinder copy number. Reference panels are supplied as in Schmid et al. [16]: 96 colonic epithelial cells [29] for HCT116, 200 epidermal cells [30] for A375 and the 450 author-annotated non-malignant cells for the astrocytoma. REFCON uses no reference for prediction and has no reference row on HCT116 and A375. On the astrocytoma, REFCON is refined against the 450 cells (Methods, Section 4.8). Astrocytoma whole sample: all 840 cells are run, and REFCON is refined against the 63 confident-normal cells it detects. CopyVAE has no reference argument, so its whole-sample and reference rows are one run. CopyVAE gave no output on A375 with the reference cells. Numbat: Supplementary Note 4. Best per column within each block in bold.

**Supplementary Table 5** Agreement between the copy number profiles of patient-derived organoid samples. REFCON, SCEVAN, CopyKAT and CopyVAE were run on each sample without a reference. Each sample's profile covers 39 chromosome arms, and agreement is the Pearson correlation between the profiles of two samples. Pairs are grouped by how the two samples are related

| Method | Pairs | Median $r$ | Minimum $r$ | Pairs with $r \leq 0$ |
|---|---|---|---|---|
| *All pairs of samples from the same line (10 lines)* | | | | |
| REFCON | 148 | 0.96 | 0.78 | 0 |
| SCEVAN | 148 | 0.68 | −0.80 | 23 |
| CopyKAT | 148 | 0.33 | −0.49 | 24 |
| CopyVAE | 148 | 0.24 | −0.40 | 23 |
| *Pairs of samples from different patients* | | | | |
| REFCON | 950 | 0.57 | 0.18 | 0 |
| SCEVAN | 950 | 0.26 | −0.52 | 150 |
| CopyKAT | 950 | 0.18 | −0.54 | 200 |
| CopyVAE | 950 | 0.09 | −0.51 | 320 |
| *Pairs from the two lines of patient PANFR0489* | | | | |
| REFCON | 30 | 0.45 | 0.32 | 0 |
| SCEVAN | 30 | 0.05 | −0.35 | 13 |
| CopyKAT | 30 | 0.08 | −0.36 | 8 |
| CopyVAE | 30 | 0.00 | −0.23 | 15 |
| *Pairs split from one passage* | | | | |
| REFCON | 14 | 0.98 | 0.92 | 0 |
| SCEVAN | 14 | 0.72 | −0.52 | 2 |
| CopyKAT | 14 | 0.47 | −0.20 | 2 |
| CopyVAE | 14 | 0.24 | −0.07 | 1 |
| *PANFR0562, pairs without a TGF-β sample* | | | | |
| REFCON | 55 | 0.96 | 0.89 | 0 |
| SCEVAN | 55 | 0.65 | −0.58 | 10 |
| CopyKAT | 55 | 0.27 | −0.49 | 12 |
| CopyVAE | 55 | 0.27 | −0.33 | 8 |
| *PANFR0562, pairs with a TGF-β sample* | | | | |
| REFCON | 50 | 0.95 | 0.89 | 0 |
| SCEVAN | 50 | 0.68 | −0.63 | 4 |
| CopyKAT | 50 | 0.34 | −0.24 | 4 |
| CopyVAE | 50 | 0.25 | −0.34 | 7 |
| *PANFR0562, control pairs (6 untreated samples of Figure 5B)* | | | | |
| REFCON | 15 | 0.95 | 0.92 | 0 |
| SCEVAN | 15 | 0.62 | −0.58 | 5 |
| CopyKAT | 15 | 0.25 | −0.20 | 3 |
| CopyVAE | 15 | 0.32 | 0.05 | 0 |

| Comparison | Method | $n$ | REFCON | Method | $P$ |
|---|---|---|---|---|---|
| Median within-line $r$ per line | SCEVAN | 10 lines | 0.96 | 0.73 | 0.002 |
| | CopyKAT | 10 lines | 0.96 | 0.46 | 0.002 |
| Nearest biopsy is the own patient's | SCEVAN | 48 samples | 46 | 32 | $1.2 \times 10^{-4}$ |
| | CopyKAT | 48 samples | 46 | 27 | $2.1 \times 10^{-5}$ |

Data: Raghavan et al. [33]. Sample profile: the median over the sample's cells of each cell's mean prediction over the genes of an arm. Pairs from different patients: one sample from each of two patients. Pairs split from one passage: two samples taken from one parent culture into different conditions. PANFR0562 pairs are grouped by whether one of the two samples was grown under TGF-$\beta$ (6, 21, 38 or 73 days). Samples without TGF-$\beta$ include the controls, three minimal-medium samples and the sample 35 days after TGF-$\beta$ removal. inferCNV returned flat profiles for every organoid sample (its per-gene values are zero for most entries) and has no correlations. Lower block: REFCON against each method. Median within-line $r$ per line: the median over lines of each line's median within-line correlation, two-sided paired Wilcoxon signed-rank test over the 10 lines (REFCON higher in all 10). Nearest biopsy: the number of organoid samples whose profile is closest (Pearson) to the biopsy of their own patient among the 22 biopsy profiles, two-sided exact McNemar test on the discordant samples.

**Supplementary Table 6** Clone shares in the PANFR0489R organoid line from the original study and from REFCON. Each column pair gives the percentage of a sample's cells in a study clone or clone group (left) and in the matched REFCON clone (right)

| | C vs A | | D+E+F vs B | | F vs B1 | | D+E vs B2 | |
|---|---|---|---|---|---|---|---|---|
| Sample | Study | REFCON | Study | REFCON | Study | REFCON | Study | REFCON |
| biopsy | 0.0 | – | 0.4 | 3.1 | 0.0 | – | 0.4 | – |
| p2 | 90.2 | 89.6 | 9.8 | 10.4 | 5.7 | 8.3 | 4.1 | 2.2 |
| p7 | 21.5 | 29.3 | 77.4 | 70.7 | 52.6 | 61.5 | 24.8 | 9.3 |
| p9 | 2.3 | 6.4 | 97.4 | 93.6 | 8.3 | 37.3 | 89.1 | 56.3 |
| p10 | 1.0 | 2.1 | 98.9 | 97.9 | 25.5 | 30.6 | 73.4 | 67.4 |
| p11 | 0.7 | 10.2 | 99.2 | 89.8 | 5.4 | 16.7 | 93.8 | 73.1 |
| Mean \|difference\| | 4.6 | | 4.3 | | 11.4 | | 15.4 | |
| Pearson $r$ | 0.995 | | 0.996 | | 0.871 | | 0.962 | |

Study: inferCNV clones with the non-malignant biopsy cells as reference (Raghavan et al. [33], Figure S5J). Study clones C to F are as labeled there. REFCON: no reference. Supplementary Note 1 defines REFCON's clones and describes how the study's values were read and matched to them. Clone A carries the 3p loss. Clone B carries the whole chromosome 9 loss, and B1 and B2 are its forms with a stronger and a weaker 7p gain. Biopsy cells carry no A, B1 or B2 label. REFCON's biopsy share of clone B is the fraction of the biopsy's malignant cells in the lower group of the chromosome 9 split. –: not available. Mean |difference| and Pearson $r$ are over passages 2 to 11.

**Supplementary Table 7** Aneuploid versus diploid classification by REFCON, SCEVAN and CopyKAT on the 94 atlas tumors and on 17 held-out patients of two further studies. Each sample was processed whole, without a reference. Part A: medians over tumors and paired tests. Part B: tumors with a score below 0.5. Part C: scores pooled over the cells of each cancer type and malignant-share group. Part D: scores pooled over the cells of each held-out cohort

| | Median | | | vs SCEVAN | | vs CopyKAT | |
|---|---|---|---|---|---|---|---|
| Metric | REFCON | SCEVAN | CopyKAT | H/L[a] | $p$ | H/L[a] | $p$ |
| *Part A. 94 atlas tumors, per tumor* | | | | | | | |
| Aneuploid F1 | 0.926 | 0.954 | 0.876 | 32/53 | 0.16 | 50/40 | 0.091 |
| Aneuploid recall | 0.981 | 0.960 | 0.785 | 41/37 | 0.53 | 60/25 | $< 0.001$ |
| Aneuploid precision | 0.975 | 0.997 | 0.996 | 16/49 | $< 0.001$ | 26/44 | 0.23 |
| Diploid recall | 0.990 | 0.972 | 0.945 | 55/28 | 0.017 | 70/20 | $< 0.001$ |
| Diploid F1 | 0.947 | 0.969 | 0.866 | 50/38 | 0.19 | 66/26 | $< 0.001$ |
| Cells labeled | 1.000 | 0.990 | 0.976 | 81/0 | $< 0.001$ | 88/0 | $< 0.001$ |

| Below 0.5 | REFCON | SCEVAN | CopyKAT |
|---|---|---|---|
| *Part B. Atlas tumors below 0.5 (this method only)* | | | |
| Aneuploid F1 | 20 (5) | 21 (3) | 29 (10) |
| Aneuploid recall | 17 (5) | 19 (0) | 35 (15) |
| Diploid recall | 4 (2) | 10 (2) | 25 (17) |
| Immune aneuploid $> 50\%$[b] | 1 (1) | 5 (3) | 15 (13) |

| | | Aneuploid F1 | | | Diploid recall | | |
|---|---|---|---|---|---|---|---|
| Atlas tumors | Tumors | REFCON | SCEVAN | CopyKAT | REFCON | SCEVAN | CopyKAT |
| *Part C. Pooled over cells, by cancer type and malignant share* | | | | | | | |
| Glioblastoma | 10 | 0.71 | 0.79 | 0.35 | 0.99 | 0.93 | 0.49 |
| Pancreatic | 23 | 0.90 | 0.83 | 0.66 | 0.95 | 0.88 | 0.83 |
| Colorectal | 15 | 0.95 | 0.96 | 0.84 | 0.91 | 0.99 | 0.95 |
| Melanoma | 6 | 0.63 | 0.38 | 0.53 | 1.00 | 0.11 | 0.19 |
| Liver | 8 | 0.85 | 0.77 | 0.72 | 0.98 | 0.98 | 0.93 |
| Ovarian | 8 | 0.88 | 0.93 | 0.94 | 0.98 | 0.91 | 0.92 |
| Head and neck | 16 | 0.82 | 0.70 | 0.70 | 0.86 | 0.69 | 0.60 |
| Renal cell | 8 | 0.87 | 0.96 | 0.81 | 1.00 | 0.87 | 0.67 |
| Share $\leq 10\%$[c] | 9 | 0.10 | 0.03 | 0.02 | 0.89 | 0.28 | 0.32 |
| Share 10–20% | 10 | 0.84 | 0.66 | 0.69 | 0.98 | 0.90 | 0.91 |
| Share 20–80% | 49 | 0.88 | 0.95 | 0.73 | 0.95 | 0.88 | 0.72 |
| Share 80–90% | 10 | 0.89 | 0.98 | 0.96 | 0.94 | 0.97 | 0.92 |
| Share $> 90\%$ | 16 | 0.84 | 0.76 | 0.56 | 0.94 | 0.93 | 0.64 |

| Method | Call rate | Aneuploid | | | Diploid | | |
|---|---|---|---|---|---|---|---|
| | | Prec. | Rec. | F1 | Prec. | Rec. | F1 |
| *Part D. Pooled over cells:* 17 *held-out patients of two further studies* | | | | | | | |
| 12 *melanoma patients [17]* (2,285 *cells)* | | | | | | | |
| REFCON | 1.00 | **0.72** | **0.98** | **0.83** | **0.99** | **0.88** | **0.93** |
| SCEVAN | 1.00 | 0.46 | 0.96 | 0.63 | 0.98 | 0.65 | 0.78 |
| CopyKAT | 0.24 | 0.69 | 0.24 | 0.36 | 0.75 | 0.16 | 0.26 |
| 5 *colorectal patients [35], all tissues together* (194,815 *cells)* | | | | | | | |
| REFCON | 1.00 | **0.90** | **0.96** | **0.93** | **0.99** | **0.98** | **0.99** |
| SCEVAN | 0.62 | 0.40 | 0.86 | 0.55 | 0.98 | 0.40 | 0.57 |
| CopyKAT | 0.57 | 0.44 | 0.90 | 0.59 | **0.99** | 0.36 | 0.53 |

Unlabeled cells count as misses. Atlas tumors: malignant against all annotated non-malignant cells. Part A: median over tumors and the two-sided paired Wilcoxon signed-rank $p$ value of REFCON against each other method. [a]Tumors in which REFCON scored higher / lower. Tumors with equal scores are left out of the test. F1 is 0 when a class present in the tumor is never called. Aneuploid precision is undefined when a method labels no cell aneuploid. Its median is over the tumors with a value, and its tests use the tumors where both methods have one (79 tumors). Part B: number of tumors with a score below 0.5, and in parentheses the tumors below 0.5 for this method only. Diploid recall is the share of non-malignant cells labeled diploid. [b]Tumors with more than half of their immune cells labeled aneuploid, over the 91 tumors with at least 10 immune cells. Part C: pooled over the cells of each group. [c]Malignant share: malignant cells over the malignant and immune cells of the tumor. Malignant and non-malignant cells per share group: 202 and 20,297 ($\leq$ 10%), 1,708 and 11,398 (10–20%), 25,370 and 36,370 (20–80%), 13,245 and 9,659 (80–90%) and 19,174 and 2,183 (> 90%). Part D: pooled over the cells of each cohort. *Call rate* is the fraction of cells the method labels. Best per column in bold. Supplementary Data 2 and 3 give the values per sample.

**Supplementary Table 8** Percentage of immune cells labeled aneuploid inside 94 tumors of eight cancer types from the Curated Cancer Cell Atlas, by cell type and by cancer type. Each tumor sample was processed whole by every method, without a reference, and scored on its immune cells

| Cells | *n* | REFCON | SCEVAN | CopyKAT |
|---|---|---|---|---|
| *By cell type* | | | | |
| T cells | 25,081 | 1.2 | 15.0 | 5.7 |
| Macrophages | 18,293 | 1.2 | 4.5 | 22.6 |
| B cells | 5,613 | 5.2 | 5.5 | 2.2 |
| Plasma cells | 1,917 | 2.0 | 32.6 | 30.7 |
| Dendritic cells | 1,456 | 19.4 | 9.9 | 26.9 |
| NK cells | 1,221 | 0.0 | 22.5 | 5.1 |
| Monocytes | 502 | 0.2 | 3.6 | 9.2 |
| Mast cells | 100 | 3.0 | 18.0 | 41.0 |
| **All immune cells** | 54,183 | 2.1 | 11.0 | 12.6 |
| Unlabeled, all immune cells | 54,183 | 0.0 | 21.6 | 25.4 |
| *By cancer type* | | | | |
| Glioblastoma | 3,379 | 0.6 | 5.7 | 44.5 |
| Pancreatic | 13,608 | 4.5 | 9.2 | 14.2 |
| Colorectal | 7,689 | 4.1 | 0.0 | 1.2 |
| Melanoma | 10,656 | 0.1 | 19.0 | 7.0 |
| Liver | 339 | 3.2 | 1.2 | 1.5 |
| Ovarian | 2,405 | 1.1 | 1.1 | 0.8 |
| Head and neck | 10,346 | 1.2 | 18.4 | 11.8 |
| Renal cell | 5,761 | 0.2 | 10.1 | 22.6 |

Tumors: the 94 atlas samples (Supplementary Data 2) with raw UMI counts and at least 10 malignant cells, from eight studies, one per cancer type: glioblastoma [7] (10x samples), pancreatic [33], colorectal [44], melanoma [45], liver [46], ovarian [47], head and neck [48] and renal cell carcinoma [49] (seven clear cell and one chromophobe tumor). Normal-tissue samples are excluded. Cell types: the atlas annotations [34], taken from the original studies or made by the atlas and checked against known marker genes. Only immune types are scored. Stromal and epithelial cells, oligodendrocytes, cells without a type and erythrocytes are not scored. Immune cells are treated as diploid. Labeled aneuploid: labeled aneuploid by REFCON or CopyKAT, or tumor by SCEVAN, as a percentage of all immune cells of the row, unlabeled cells included. Unlabeled: the percentage left unlabeled. $n$: immune cells.

**Supplementary Table 9** Specificity of REFCON, SCEVAN and CopyKAT on the fully normal controls: the PBMC 3K healthy-donor dataset [36] and healthy cells of 18 organs from Tabula Sapiens [25, 26]. Each sample was processed whole, without a reference. Part A: PBMC 3K. Part B: per control group. Part C: Tabula Sapiens per organ. Part D: paired tests over the 19 control samples

| Sample | Method | Cells | Call rate | FP | Specificity | Mean $\sigma$ |
|---|---|---|---|---|---|---|
| *PBMC 3K healthy-donor dataset [36]* | | | | | | |
| PBMC 3K | REFCON | 2,700 | 1.00 | 0 | **1.00** | 0.083 |
| PBMC 3K | SCEVAN | 2,700 | 0.55 | 1,089 | 0.27 | — |
| PBMC 3K | CopyKAT | 2,700 | 0.45 | 393 | 0.68 | — |

All cells are non-malignant, so every aneuploid label is a false positive (FP). Call rate: the fraction of cells a method labels. Specificity: TN/(TN + FP) over the cells a method labels. Mean $\sigma$: REFCON's mean per-cell dispersion of the predicted copy number ratios (Methods, Section 4.7). The sample-level threshold is $T_\sigma = 0.125$. $\sigma$ is not given for SCEVAN and CopyKAT, whose outputs are on other scales. Best specificity in bold.

**Part B.** Specificity per cohort, among classified cells

| Method | Classified | FP | Specificity |
|---|---|---|---|
| *PBMC (healthy donor)* | | | |
| REFCON | 2,700 | 0 | **1.000** |
| SCEVAN | 1,495 | 1,089 | 0.272 |
| CopyKAT | 1,228 | 393 | 0.680 |
| *Tabula Sapiens (18 organs)* | | | |
| REFCON | 14,400 | 2,212 | **0.846** |
| SCEVAN | 14,309 | 7,432 | 0.481 |
| CopyKAT | 14,279 | 6,845 | 0.521 |

Classified: the cells a method labels. Specificity as in Part A. Best specificity per group in bold.

**Part C.** Tabula Sapiens: specificity among classified cells per organ, with the call rate of SCEVAN and CopyKAT

| Organ | Donor | REFCON | SCEVAN call | SCEVAN | CopyKAT call | CopyKAT | Mean $\sigma$ |
|---|---|---|---|---|---|---|---|
| Bladder | TSP27 | **0.83** | 1.00 | 0.28 | 1.00 | 0.40 | 0.077 |
| Fat | TSP21 | **1.00** | 0.99 | 0.18 | 0.99 | 0.83 | 0.076 |
| Heart | TSP12 | 0.42 | 1.00 | 0.59 | 1.00 | **0.67** | 0.078 |
| Kidney | TSP2 | **0.94** | 1.00 | 0.23 | 1.00 | 0.22 | 0.077 |
| Large Intestine | TSP27 | 0.75 | 1.00 | **0.87** | 1.00 | 0.14 | 0.080 |
| Liver | TSP19 | **0.83** | 0.93 | 0.37 | 0.93 | 0.36 | 0.077 |
| Lung | TSP2 | **1.00** | 1.00 | 0.27 | 1.00 | 0.30 | 0.079 |
| Lymph Node | TSP25 | **1.00** | 0.99 | 0.29 | 0.98 | 0.93 | 0.075 |
| Mammary | TSP27 | 0.50 | 1.00 | **0.83** | 1.00 | 0.18 | 0.081 |
| Muscle | TSP27 | **1.00** | 1.00 | 0.78 | 1.00 | 0.32 | 0.080 |
| Ovary | TSP28 | **1.00** | 0.99 | 0.15 | 0.99 | 0.43 | 0.072 |
| Pancreas | TSP9 | **1.00** | 1.00 | 0.74 | 1.00 | 0.77 | 0.081 |
| Prostate | TSP8 | **0.88** | 0.99 | 0.56 | 0.99 | 0.56 | 0.077 |
| Skin | TSP21 | **1.00** | 0.99 | 0.66 | 0.99 | 0.42 | 0.078 |
| Small Intestine | TSP26 | **1.00** | 1.00 | 0.13 | 1.00 | 0.87 | 0.078 |
| Stomach | TSP27 | **1.00** | 1.00 | 0.39 | 1.00 | 0.45 | 0.078 |
| Tongue | TSP27 | 0.57 | 0.99 | 0.77 | 0.99 | **0.80** | 0.085 |
| Uterus | TSP4 | 0.50 | 1.00 | 0.56 | 1.00 | **0.72** | 0.084 |

One 10x sample of 800 cells per organ, from 11 donors who contribute no training or validation cells. REFCON labels every cell. The REFCON, SCEVAN and CopyKAT columns are specificities among classified cells, and the call columns the fraction of cells SCEVAN and CopyKAT label. Mean $\sigma$ as in Part A. Best specificity per organ in bold.

**Part D.** Specificity of REFCON against each method over the 19 control samples

| Method | Samples | Median REFCON | Median method | H/L[a] | $p$ |
|---|---|---|---|---|---|
| SCEVAN | 19 | 1.00 | 0.39 | 14/5 | $7.9 \times 10^{-4}$ |
| CopyKAT | 19 | 1.00 | 0.45 | 16/3 | $7.9 \times 10^{-4}$ |

Samples: PBMC 3K and the 18 Tabula Sapiens organs. Specificity among classified cells, as in Part C. [a]Samples in which REFCON scored higher / lower. $p$: two-sided paired Wilcoxon signed-rank test.

**Supplementary Table 10** Detection AUROC by copy number event size on the three cohorts with paired single-cell DNA, every method without a reference. Events are contiguous gain or loss runs in the consensus DNA profile of each cohort, grouped by genomic length (Part A) or by number of genes (Part B). For each size class, the genes of its events are the positives and the copy-neutral genes the negatives

| *Part A. By genomic length (Mb)* | | | | |
|---|---|---|---|---|
| Method | focal (<3 Mb) | mid (3–20 Mb) | arm (20–100 Mb) | chr (>100 Mb) |
| *HCT116 (near-diploid; no chromosome-scale events)* | | | | |
| REFCON | 0.49 | 0.68 | 0.81 | — |
| SCEVAN | 0.40 | 0.59 | 0.58 | — |
| CopyKAT | 0.46 | 0.53 | 0.56 | — |
| CopyVAE | 0.47 | 0.49 | 0.50 | — |
| inferCNV | 0.50 | 0.50 | 0.50 | — |
| *A375 (reference-free; near-triploid)* | | | | |
| REFCON | 0.62 | 0.62 | 0.75 | 0.75 |
| SCEVAN | 0.55 | 0.61 | 0.62 | 0.60 |
| CopyKAT | 0.49 | 0.53 | 0.57 | 0.57 |
| CopyVAE | 0.53 | 0.44 | 0.48 | 0.44 |
| inferCNV | 0.50 | 0.49 | 0.48 | 0.49 |
| *scONE-seq astrocytoma (reference-free)* | | | | |
| REFCON | 0.57 | 0.62 | 0.78 | — |
| SCEVAN | 0.50 | 0.51 | 0.53 | — |
| CopyKAT | 0.51 | 0.52 | 0.61 | — |
| CopyVAE | 0.48 | 0.48 | 0.46 | — |
| inferCNV | 0.50 | 0.50 | 0.49 | — |

| *Part B. By event length in genes, relative to the 8-gene model bin* | | | | |
|---|---|---|---|---|
| Method | <8 genes | 8–16 genes | 16–64 genes | ≥64 genes |
| *HCT116 (reference-free; near-diploid)* | | | | |
| REFCON | 0.40 | 0.59 | 0.71 | 0.73 |
| SCEVAN | 0.54 | 0.55 | 0.51 | 0.57 |
| CopyKAT | 0.52 | 0.54 | 0.51 | 0.54 |
| CopyVAE | 0.49 | 0.40 | 0.43 | 0.50 |
| inferCNV | 0.49 | 0.49 | 0.49 | 0.50 |
| *A375 (reference-free; near-triploid)* | | | | |
| REFCON | — | 0.53 | 0.66 | 0.75 |
| SCEVAN | — | 0.51 | 0.58 | 0.62 |
| CopyKAT | — | 0.54 | 0.51 | 0.57 |
| CopyVAE | — | 0.42 | 0.46 | 0.47 |
| inferCNV | — | 0.50 | 0.50 | 0.48 |
| *scONE-seq astrocytoma (reference-free)* | | | | |
| REFCON | 0.55 | 0.55 | 0.51 | 0.74 |
| SCEVAN | 0.50 | 0.51 | 0.49 | 0.53 |
| CopyKAT | 0.47 | 0.50 | 0.53 | 0.59 |
| CopyVAE | 0.51 | 0.49 | 0.48 | 0.47 |
| inferCNV | 0.50 | 0.50 | 0.50 | 0.50 |

Per-cell AUROC as in the main benchmark (Methods, Section 4.9), except that the classes come from the consensus profile below. Genes a method does not call are filled with the median prediction. Loss is scored with the negative of the prediction and gain with the prediction, and 0.5 is chance. Consensus profile: the per-gene mean DNA copy number over cells divided by its median. Events: runs of at least 3 genes within a chromosome above 1.15 (gain) or below 0.85 (loss). Copy-neutral genes: consensus within $[0.90, 1.10]$. Part B classes are relative to REFCON's 8-gene bin. —: no event in the class, or one event only (A375, fewer than 8 genes), not scored. Events per class, Part A (focal/mid/arm/chromosome): HCT116 8/11/6/0, A375 7/8/15/3, astrocytoma 261/29/10/0. Part B ($<8$/8–16/16–64/$\geq 64$ genes): HCT116 6/4/7/8, A375 1/2/10/20, astrocytoma 136/75/63/26.

**Supplementary Table 11** Per-seed and ensemble metrics. Three models trained with the released recipe at context length 2,048 (seeds 1, 2 and 3) are scored alone, in every pair and all together. The released REFCON averages seeds 2 and 3

| Configuration | Pearson | Spearman | $AUROC_L$ | $AUROC_G$ |
|---|---|---|---|---|
| *CCLE validation (7 lines)* | | | | |
| Seed 1 | 0.701 | 0.722 | **0.871** | 0.921 |
| Seed 2 | 0.716 | **0.738** | 0.865 | 0.930 |
| Seed 3 | 0.712 | 0.727 | 0.864 | 0.928 |
| Seeds 1 + 2 | 0.713 | 0.734 | **0.870** | 0.928 |
| Seeds 1 + 3 | 0.708 | 0.727 | 0.868 | 0.926 |
| REFCON 2-model ensemble (seeds 2 + 3) | **0.718** | 0.736 | 0.866 | **0.931** |
| Seeds 1 + 2 + 3 | 0.714 | 0.734 | 0.869 | 0.929 |
| *HCT116 (DNTR-seq, $n$=1,467)* | | | | |
| Seed 1 | 0.432 | 0.389 | 0.740 | 0.834 |
| Seed 2 | 0.420 | 0.380 | 0.728 | 0.833 |
| Seed 3 | 0.422 | 0.385 | 0.738 | 0.827 |
| Seeds 1 + 2 | **0.434** | 0.390 | 0.737 | **0.838** |
| Seeds 1 + 3 | 0.433 | **0.392** | **0.742** | 0.834 |
| REFCON 2-model ensemble (seeds 2 + 3) | 0.429 | 0.389 | 0.737 | 0.835 |
| Seeds 1 + 2 + 3 | **0.434** | **0.392** | 0.740 | 0.837 |
| *A375 (DNTR-seq, $n$=64)* | | | | |
| Seed 1 | **0.493** | **0.508** | 0.738 | **0.774** |
| Seed 2 | 0.452 | 0.469 | 0.717 | 0.751 |
| Seed 3 | 0.469 | 0.473 | 0.743 | 0.757 |
| Seeds 1 + 2 | 0.482 | 0.496 | 0.732 | 0.768 |
| Seeds 1 + 3 | 0.489 | 0.499 | **0.745** | 0.770 |
| REFCON 2-model ensemble (seeds 2 + 3) | 0.470 | 0.479 | 0.734 | 0.759 |
| Seeds 1 + 2 + 3 | 0.483 | 0.494 | 0.738 | 0.767 |

| Configuration | Pearson | Spearman | $AUROC_L$ | $AUROC_G$ |
|---|---|---|---|---|
| *CCLE esoph. (7 lines)* | | | | |
| Seed 1 | 0.545 | 0.633 | **0.724** | 0.812 |
| Seed 2 | 0.539 | 0.626 | 0.695 | 0.812 |
| Seed 3 | 0.554 | 0.634 | 0.720 | 0.816 |
| Seeds 1 + 2 | 0.550 | 0.640 | 0.714 | 0.818 |
| Seeds 1 + 3 | **0.555** | 0.641 | 0.723 | 0.818 |
| REFCON 2-model ensemble (seeds 2 + 3) | 0.553 | 0.639 | 0.711 | 0.819 |
| Seeds 1 + 2 + 3 | **0.555** | **0.643** | 0.717 | **0.820** |
| *CCLE mixed (3 lines)* | | | | |
| Seed 1 | 0.448 | 0.450 | 0.715 | 0.711 |
| Seed 2 | 0.446 | 0.448 | 0.714 | 0.757 |
| Seed 3 | **0.467** | **0.481** | 0.707 | 0.762 |
| Seeds 1 + 2 | 0.456 | 0.458 | 0.717 | 0.741 |
| Seeds 1 + 3 | 0.463 | 0.473 | 0.714 | 0.742 |
| REFCON 2-model ensemble (seeds 2 + 3) | **0.467** | 0.476 | 0.721 | **0.764** |
| Seeds 1 + 2 + 3 | 0.464 | 0.473 | **0.723** | 0.751 |
| *MIX-seq (49 lines)* | | | | |
| Seed 1 | 0.549 | 0.594 | 0.764 | 0.829 |
| Seed 2 | 0.555 | 0.610 | 0.772 | 0.839 |
| Seed 3 | 0.561 | 0.607 | 0.772 | 0.834 |
| Seeds 1 + 2 | 0.562 | 0.613 | 0.772 | 0.841 |
| Seeds 1 + 3 | 0.561 | 0.607 | 0.770 | 0.835 |
| REFCON 2-model ensemble (seeds 2 + 3) | **0.566** | **0.618** | **0.775** | **0.843** |
| Seeds 1 + 2 + 3 | **0.566** | 0.616 | 0.774 | 0.841 |
| *scONE-seq ($n$=390)* | | | | |
| Seed 1 | 0.355 | 0.346 | 0.696 | 0.657 |
| Seed 2 | **0.362** | 0.344 | 0.693 | 0.658 |
| Seed 3 | 0.351 | 0.339 | 0.692 | 0.654 |
| Seeds 1 + 2 | **0.362** | **0.349** | **0.697** | **0.659** |
| Seeds 1 + 3 | 0.357 | 0.345 | 0.696 | 0.657 |
| REFCON 2-model ensemble (seeds 2 + 3) | 0.360 | 0.345 | 0.694 | 0.657 |
| Seeds 1 + 2 + 3 | 0.361 | 0.347 | 0.696 | 0.658 |

The seeds differ only in their random seed. An ensemble is the per-cell mean of its members, renormalized to mean one over finite bins. Seeds 2 and 3 were chosen for the release by validation Pearson correlation. Metrics are per-cell means against paired single-cell DNA (HCT116, A375, astrocytoma) and per cell line pseudobulk means against bulk DNA (CCLE validation with 150 cells per line, esophageal and mixed-tissue lines, MIX-seq). Subscripts L and G denote loss and gain. Predictions are unrefined. Best per column within each cohort in bold.

**Supplementary Table 12** Ablations of the released model. Each variant was retrained with the released recipe at context length 2,048 with one choice changed, using seeds 2 and 3 (those of the released members), and the two models were averaged per cell. Every variant was run without a reference. Part A: rotary positions. Part B: input normalization. Part C: context length. Part D: training additions left out one at a time

**Part A.** Rotary positions in megabases (released), over the bin index, or removed

| | A375 ($n = 64$) | | | HCT116 ($n$ = 1,467) | | | Val. |
|---|---|---|---|---|---|---|---|
| Variant | $r_P$ | $\text{AUROC}_\text{L}$ | $\text{AUROC}_\text{G}$ | $r_P$ | $\text{AUROC}_\text{L}$ | $\text{AUROC}_\text{G}$ | $r$ |
| Released (megabase RoPE) | **0.470** | **0.734** | **0.759** | **0.429** | **0.737** | **0.835** | **0.718** |
| RoPE over the bin index | 0.461 | 0.732 | 0.751 | 0.313 | 0.734 | 0.740 | 0.697 |
| ablate RoPE | 0.443 | 0.728 | 0.756 | 0.394 | 0.714 | 0.809 | 0.658 |

Rotary position embeddings (RoPE) act on the query and key projections of every attention layer. Released: each bin's genomic offset in megabases. Bin index: the bin's rank in the window. In both ablated rows, the chromosome boundary of a bridge window is still found from the genomic positions, so the two chromosomes stay apart. $r_P$: per-cell mean Pearson correlation with paired single-cell DNA (Supplementary Note 3). Subscripts L and G denote loss and gain. Val.: mean per cell line pseudobulk Pearson correlation $r$ on the seven held-out CCLE validation lines, 150 cells per line (Methods, Section 4.3). Differences within the seed spread do not separate a variant from the release (Methods, Section 4.10). Best per column in bold.

**Part B.** Input normalization: masked InstanceNorm (released), none, or BatchNorm

| | A375 ($n = 64$) | | | HCT116 ($n$ = 1,467) | | | Val. |
|---|---|---|---|---|---|---|---|
| Variant | $r_P$ | $\text{AUROC}_\text{L}$ | $\text{AUROC}_\text{G}$ | $r_P$ | $\text{AUROC}_\text{L}$ | $\text{AUROC}_\text{G}$ | $r$ |
| Released (InstanceNorm) | **0.470** | **0.734** | **0.759** | **0.429** | **0.737** | **0.835** | **0.718** |
| remove input normalization | 0.131 | 0.597 | 0.556 | 0.213 | 0.680 | 0.666 | 0.475 |
| InstanceNorm → BatchNorm | 0.238 | 0.634 | 0.634 | 0.183 | 0.657 | 0.621 | 0.479 |

Masked InstanceNorm scales each cell's log-expression input by its mean and standard deviation over the real genes of the window, before binning (Methods, Section 4.2). The variants remove input normalization or replace it with BatchNorm. Metrics as in Part A. Best per column in bold.

**Part C.** Context length of 512, 1,024 or 2,048 genes

| Cohort | $C = 512$ | $C$ = 1,024 | $C$ = 2,048 |
|---|---|---|---|
| CCLE validation (7 lines) | 0.625 | 0.674 | **0.718** |
| scONE-seq astrocytoma | 0.245 | 0.300 | **0.360** |
| HCT116 (DNTR-seq) | 0.284 | 0.383 | **0.429** |
| CCLE esophageal | 0.491 | 0.535 | **0.553** |
| CCLE mixed-tissue | 0.421 | 0.445 | **0.467** |
| Mean Δ vs $C$ = 2,048 | −0.09 | −0.04 | — |

Per-cell mean Pearson correlation with paired single-cell DNA (astrocytoma, HCT116) and per cell line pseudobulk Pearson correlation with bulk DNA (CCLE validation with 150 cells per line, esophageal and mixed-tissue lines). Each column averages two models trained at that context length. Δ: difference from $C$ = 2,048, averaged over the four test cohorts. Per-seed metrics at $C$ = 2,048 are in Supplementary Table 11. Best per row in bold.

**Part D.** Training additions left out one at a time

| Variant | Pearson | | | | | | | False events | |
|---|---|---|---|---|---|---|---|---|---|
| | Val. | HCT116 | A375 | scONE | Esoph. | Mixed | MIX-seq | 25% | 50% |
| Released ensemble | 0.718 | 0.429 | 0.470 | **0.360** | 0.553 | 0.467 | **0.566** | 0.64 | 5.38 |
| no healthy cells | 0.715 | **0.430** | 0.465 | 0.354 | 0.554 | 0.467 | 0.563 | 1.10 | 6.32 |
| no count thinning | 0.725 | 0.425 | **0.478** | 0.354 | 0.557 | 0.459 | **0.566** | **0.59** | **4.99** |
| no removal/zeroing | **0.726** | 0.408 | 0.466 | 0.348 | 0.550 | 0.469 | 0.563 | 1.03 | 6.39 |
| no separation | 0.696 | 0.416 | 0.458 | 0.332 | **0.563** | **0.470** | 0.534 | 1.59 | 8.34 |

Left out: the 4,200 healthy Tabula Sapiens training cells labeled diploid (no healthy cells), binomial count thinning (no count thinning), gene removal and zeroing (no removal/zeroing), or the separation of the two chromosomes in a bridge window, with the megabase positions kept (no separation). Pearson: mean per cell line pseudobulk correlation with bulk DNA on the seven held-out CCLE validation lines (Val., 150 cells per line), the CCLE esophageal (Esoph.) and mixed-tissue (Mixed) lines and the 49 MIX-seq lines, and per-cell mean correlation with paired single-cell DNA on HCT116, A375 and the astrocytoma (scONE). False events: arm-level copy number events per cell on the 1,200 cells of the two healthy validation donors (TSP7 and TSP10). Each model is called at its own matched-sensitivity threshold, the absolute arm value at which it calls 25% (or 50%) of the DNA-defined arm events of the ten held-out CCLE test lines in the correct direction. A DNA-defined arm event differs by at least one copy from the line's ploidy. Every left-out run has a higher HCT116 loss AUROC than the release (0.751 to 0.765, against 0.737). Best per column in bold, and for false events the fewest.

**Supplementary Table 13** Stitching ablation. The released two-model ensemble was run without a reference, and only the step that merges the window predictions into one genome-wide profile per cell was changed. Part A: per-cell metrics on the three cohorts with paired single-cell DNA. Part B: per cell line metrics on the seven held-out CCLE validation lines

| Merge variant | $r_P$ | $\mathrm{AUROC_L}$ | $\mathrm{AUROC_G}$ |
|---|---|---|---|
| *HCT116 (near-diploid)* | | | |
| Released pipeline | **0.429** | **0.737** | **0.835** |
| without bidirectional averaging | 0.417 | 0.716 | 0.833 |
| without bridge windows | 0.333 | 0.700 | 0.774 |
| Sequential chaining (not joint LSQ) | 0.397 | 0.715 | 0.829 |
| *A375 (near-triploid)* | | | |
| Released pipeline | 0.470 | 0.734 | **0.759** |
| without bidirectional averaging | **0.471** | **0.745** | 0.754 |
| without bridge windows | 0.260 | 0.630 | 0.639 |
| Sequential chaining (not joint LSQ) | 0.385 | 0.700 | 0.717 |
| *scONE-seq (aneuploid)* | | | |
| Released pipeline | **0.360** | **0.694** | 0.657 |
| without bidirectional averaging | 0.358 | 0.687 | **0.659** |
| without bridge windows | 0.272 | 0.646 | 0.624 |
| Sequential chaining (not joint LSQ) | 0.280 | 0.659 | 0.617 |

Released pipeline: each window averaged with its gene-order-reversed pass (bidirectional averaging), chromosome windows plus bridge windows, and a joint least-squares fit in log space over all windows (Methods, Section 4.4). Without bidirectional averaging: forward passes only. Without bridge windows: chromosome windows only. Sequential chaining: each chromosome stitched on its own, then joined through the bridge windows in order. $r_P$: per-cell mean Pearson correlation with paired single-cell DNA. Subscripts L and G denote loss and gain. Genes without a prediction are filled with the median prediction. Best per column within each cohort in bold.

**Part B.** Mean per cell line pseudobulk Pearson and Spearman correlation with bulk DepMap copy number on the seven held-out CCLE validation lines

| Merge variant | Pearson | Spearman |
|---|---|---|
| Released pipeline | 0.718 | 0.734 |
| without bidirectional averaging | 0.714 | 0.734 |
| without bridge windows | 0.450 | 0.444 |
| Sequential chaining (not joint LSQ) | **0.729** | **0.735** |

Validation lines: DAOY, BT549, HOS, SW1990, SNU449, KMRC3 and OVSAHO (2,255 cells; Methods, Section 4.1), the cohort used for model selection. Rows as in Part A. Best per column in bold.

**Supplementary Table 14** Sensitivity of the refinement to the SVD rank $k$ on the scONE-seq astrocytoma. All 840 cells were refined at $k = 0$ to 10, against the 63 confident-normal cells REFCON detects or against the 450 author-annotated non-malignant cells, and the 390 malignant cells were scored against paired single-cell DNA

| $k$ | Pearson | Spearman | $AUROC_L$ | $AUROC_G$ | $AUPRC_L$ | $AUPRC_G$ |
|---|---|---|---|---|---|---|
| *Refined against the 63 self-detected confident-normal cells (Methods, Section 4.5)* | | | | | | |
| 0 | 0.507 | 0.483 | 0.776 | 0.720 | 0.598 | 0.503 |
| 1 | **0.510** | **0.485** | **0.778** | **0.721** | 0.601 | 0.504 |
| 2 | **0.510** | 0.483 | 0.777 | 0.720 | **0.602** | 0.504 |
| 3† | **0.510** | 0.483 | 0.777 | 0.720 | **0.602** | **0.505** |
| 4 | 0.508 | 0.481 | 0.776 | 0.719 | 0.599 | 0.503 |
| 5 | 0.507 | 0.481 | 0.776 | 0.719 | 0.599 | 0.502 |
| 6 | 0.480 | 0.450 | 0.759 | 0.704 | 0.560 | 0.490 |
| 7 | 0.459 | 0.436 | 0.752 | 0.697 | 0.540 | 0.474 |
| 8 | 0.456 | 0.431 | 0.750 | 0.694 | 0.537 | 0.473 |
| 9 | 0.455 | 0.430 | 0.749 | 0.694 | 0.536 | 0.473 |
| 10 | 0.449 | 0.425 | 0.746 | 0.691 | 0.533 | 0.467 |
| *Refined against the 450 author-annotated non-malignant cells* | | | | | | |
| 0 | **0.517** | **0.492** | **0.782** | **0.724** | **0.607** | 0.504 |
| 1 | 0.516 | 0.491 | 0.781 | **0.724** | 0.604 | 0.506 |
| 2 | 0.515 | 0.485 | 0.779 | 0.720 | 0.603 | 0.507 |
| 3† | **0.517** | 0.486 | 0.779 | 0.721 | 0.605 | **0.508** |
| 4 | 0.512 | 0.479 | 0.776 | 0.718 | 0.599 | 0.506 |
| 5 | 0.511 | 0.484 | 0.779 | 0.719 | 0.604 | 0.503 |
| 6 | 0.510 | 0.482 | 0.779 | 0.718 | 0.602 | 0.502 |
| 7 | 0.501 | 0.472 | 0.772 | 0.714 | 0.588 | 0.498 |
| 8 | 0.483 | 0.455 | 0.761 | 0.707 | 0.563 | 0.487 |
| 9 | 0.476 | 0.450 | 0.759 | 0.704 | 0.554 | 0.481 |
| 10 | 0.474 | 0.447 | 0.757 | 0.703 | 0.551 | 0.480 |

Truth: paired scWGS (Yu et al. [12]). Each ensemble member is refined at rank $k$ (Methods, Section 4.8), and the refined members are averaged per cell. $k = 0$ applies only the diploid-bias correction. Without refinement, REFCON's per-cell Pearson correlation is 0.360 (Supplementary Table 4). Subscripts L and G denote loss and gain. †Released default, fixed before this analysis. Best per column within each panel in bold, ties at three decimals included.

**Supplementary Table 15** Loss and gain AUROC under absolute and ploidy-relative copy number classes on the three cohorts with paired single-cell DNA. Predictions are the reference-free predictions of Supplementary Table 4. Only the class labels of the DNA truth change

| | $\text{AUROC}_\text{L}$ | | $\text{AUROC}_\text{G}$ | |
|---|---|---|---|---|
| Method | absolute | ploidy-rel. | absolute | ploidy-rel. |
| *HCT116 (DNTR-seq; $\pi_{gain}$ 0.14→0.13)* | | | | |
| REFCON | **0.737** | **0.733** | **0.835** | **0.835** |
| SCEVAN | 0.598 | 0.597 | 0.619 | 0.619 |
| CopyKAT | 0.589 | 0.590 | 0.598 | 0.599 |
| CopyVAE | 0.523 | 0.528 | 0.553 | 0.554 |
| inferCNV | 0.604 | 0.604 | 0.571 | 0.571 |
| *A375 (DNTR-seq; $\pi_{gain}$ 0.64→0.15)* | | | | |
| REFCON | **0.734** | **0.764** | **0.759** | **0.805** |
| SCEVAN | 0.650 | 0.624 | 0.632 | 0.671 |
| CopyKAT | 0.573 | 0.592 | 0.594 | 0.608 |
| CopyVAE | 0.432 | 0.460 | 0.463 | 0.469 |
| inferCNV | 0.502 | 0.516 | 0.515 | 0.504 |
| *scONE-seq astrocytoma ($\pi_{gain}$ 0.27→0.32)* | | | | |
| REFCON | **0.694** | **0.675** | **0.657** | **0.655** |
| SCEVAN | 0.558 | 0.554 | 0.549 | 0.548 |
| CopyKAT | 0.616 | 0.603 | 0.579 | 0.580 |
| CopyVAE | 0.504 | 0.504 | 0.502 | 0.503 |
| inferCNV | 0.541 | 0.539 | 0.541 | 0.539 |

Absolute classes: loss CN $< 1.5$ and gain CN $> 2.5$, as in the main benchmark. Ploidy-relative classes: loss CN/ploidy $< 0.85$ and gain $> 1.15$, where ploidy is the cell's mean DNA copy number over scored genes. $\pi_{\text{gain}}$: share of gain genes, absolute → ploidy-relative. Subscripts L and G denote loss and gain. Best per column within each cohort in bold.

**Supplementary Table 16** Transformer contribution on the three cohorts with paired single-cell DNA. The released REFCON is compared with the same inference pipeline in which the transformer is replaced by the mean of the normalized log-expression within each 8-gene bin. Both were run without a reference

| Method | $r_P$ | $r_S$ | $\text{AUROC}_\text{L}$ | $\text{AUROC}_\text{G}$ |
|---|---|---|---|---|
| *HCT116 (DNTR-seq, $n$=1,467)* | | | | |
| REFCON (transformer) | **0.429** | **0.389** | **0.737** | **0.835** |
| no transformer (averaging) | 0.161 | 0.183 | 0.639 | 0.638 |
| *A375 (DNTR-seq, $n$=64)* | | | | |
| REFCON (transformer) | **0.470** | **0.479** | **0.734** | **0.759** |
| no transformer (averaging) | 0.177 | 0.189 | 0.624 | 0.606 |
| *scONE-seq ($n$=390)* | | | | |
| REFCON (transformer) | **0.360** | **0.345** | **0.694** | **0.657** |
| no transformer (averaging) | 0.250 | 0.255 | 0.642 | 0.617 |

Both rows use the same input normalization (masked InstanceNorm of $\log(1 + \text{UMI})$), 8-gene bins, overlapping 2,048-gene chromosome windows and bridge windows, bidirectional averaging and joint least-squares stitching (Methods, Section 4.4). REFCON: the released two-model ensemble. Metrics: per-cell mean Pearson ($r_P$) and Spearman ($r_S$) correlation and AUROC against paired single-cell DNA. Subscripts L and G denote loss and gain. Best per column within each cohort in bold.

**Supplementary Table 17** Copy number inference with fewer genes and fewer counts per cell. Each model was run without a reference on HCT116, A375, the 49 MIX-seq lines and the scONE-seq astrocytoma after 30%, 50% or 80% of the genes were removed, or after each cell's counts were thinned to 8,000, 4,000, 2,000 or 1,000. The released REFCON is compared with the models trained without gene removal and zeroing or without count thinning (Supplementary Table 12, Part D). The table gives the unperturbed Pearson correlation and the percentage of it kept at each level

| | | Genes removed | | | Counts per cell | | | |
|---|---|---|---|---|---|---|---|---|
| Model | Unperturbed $r$ | 30% | 50% | 80% | 8,000 | 4,000 | 2,000 | 1,000 |
| *HCT116 (1,467 cells)* | | | | | | | | |
| Released | 0.429 | 90 | 85 | 61 | 97 | 92 | 84 | 74 |
| No removal/zeroing | 0.408 | 96 | 89 | 63 | — | — | — | 71 |
| No count thinning | 0.425 | — | — | 61 | 96 | 91 | 82 | 72 |
| *A375 (64 cells)* | | | | | | | | |
| Released | 0.470 | 92 | 88 | 67 | 96 | 89 | 77 | 63 |
| No removal/zeroing | 0.466 | 94 | 90 | 69 | — | — | — | 66 |
| No count thinning | 0.478 | — | — | 64 | 96 | 89 | 78 | 65 |
| *MIX-seq (49 lines)* | | | | | | | | |
| Released | 0.566 | 94 | 90 | 64 | 95 | 91 | 86 | 80 |
| No removal/zeroing | 0.563 | 95 | 90 | 60 | — | — | — | 77 |
| No count thinning | 0.566 | — | — | 63 | 94 | 89 | 84 | 78 |
| *Astrocytoma (390 malignant cells)* | | | | | | | | |
| Released | 0.360 | 102 | 99 | 93 | 97 | 93 | 88 | 81 |
| No removal/zeroing | 0.348 | 100 | 99 | 97 | — | — | — | 81 |
| No count thinning | 0.354 | — | — | 92 | 98 | 95 | 90 | 81 |

Unperturbed $r$: Pearson correlation without perturbation, the per-cell mean against paired single-cell DNA (HCT116, A375, astrocytoma) or the mean per cell line pseudobulk correlation against bulk DepMap copy number (MIX-seq). Other columns: percentage of the unperturbed $r$ of the same row. Genes removed: three random gene sets per level, averaged. Counts per cell: binomial thinning of each cell to the given total. All rows use the same gene sets and thinned counts. Released: the released two-model ensemble. No removal/zeroing and No count thinning: the two-seed models of Supplementary Table 12, Part D, each tested on its left-out perturbation and on one level of the other. —: not run.

### Supplementary Notes

**Supplementary Note 1. Organoid clones: definition and checks.** REFCON's per-cell predictions were averaged over chromosome arms and over 280 genomic windows of about 10 Mb. Within each organoid line, each window was tested for two groups of cells with a two-component Gaussian mixture. A window was called clonal when the Bayesian information criterion preferred two components, Ashman's $D$ was at least 2 and the smaller group held at least 5% of cells. In PANFR0489R, clone B is the lower group of a two-component mixture on chromosome 9, split into two subclones by the 7p window call. In PANFR0489, the late clone carries the distal 1q and 14q calls and the early clone the chromosome 11 call. Clone shares are the fractions of cells per sample. The clone tree is inferred from shared events. The checks against the original study and against SCEVAN follow.

Without a reference, SCEVAN's and CopyKAT's organoid profiles are near-flat: the median largest absolute arm value per sample is 0.12 and 0.05, against 0.23 and 0.10 in their own biopsies (REFCON: 0.52 and 0.54). Their low agreement between samples of one line therefore mostly reflects noise. The original study tracked the PANFR0489R clones with inferCNV and the biopsy's normal cells [33] (its Figures S5I and S5J). We read its clone frequencies per passage from the vector graphics of Figure S5J. Each clone is one line, the legend colors give the clone letters, and the axis is logarithmic (ticks at 100, 10, 1 and 0.1%; points on the axis were read as zero). The values sum to 100.0–100.1% at every passage. The study's heatmap in Figure S5I covers the biopsy and passage 2 only (1,246 cells) and uses its own clone letters. We therefore matched the study's clones to REFCON's by their course over passages (Supplementary Table 6). The study's own copy number supports this match. Relative to the 7p gain that every cell of the line carries, chromosome 9 is not lost at passage 2 (Figure S5I) and is lost at passage 11 (Figure S5M), when the study's clones D, E and F hold 99% of cells. The study's passage 2 cells also show the 11p gain, the 6p loss and the lower 3p that separate REFCON's culture clones from the biopsy. As a second check, SCEVAN [19] was run on the pooled organoid cells of each line with the biopsy's normal cells as its reference (`norm_cell`, with the call of Supplementary Note 4): 4,167 PANFR0489R cells with 2,115 normal cells, and 4,151 PANFR0489 cells with 84 normal cells. Its clone shares per passage follow REFCON's (Supplementary Figure 3), its per-cell values agreed with REFCON's on every clonal segment (Spearman 0.84 to 0.94), and its cells, split into two groups per segment as for REFCON, fall onto REFCON's clones. In PANFR0489R the chromosome 9 groups of the two methods agree for 95.1% of cells. In PANFR0489, 89% of the cells SCEVAN places in the late group (distal 1q loss and distal 14q gain) and 94% of those with chromosome 11 loss belong to REFCON's late and early clones. Without the normal cells, SCEVAN's profiles of the PANFR0489R passages did not agree with each other (median correlation −0.02, against 0.94 with the normal cells).

**Supplementary Note 2. Cases where labeling failed.** REFCON labels whole clusters. A cluster is labeled aneuploid when the median per-cell dispersion of its cells, relative to that of the confident-normal cells, is at least $\tau_1 = 1.05$ (Methods, Section 4.7). A sample with fewer than two confident-normal cells is labeled by the sample-level rule ($T_\sigma$). Labeling can therefore fail in four ways. The tumor cells are too few to form a cluster, normal cells cluster with tumor cells, a normal cluster lies above $\tau_1$, or a malignant cluster lies below it. We replayed the labeling of each case and recorded the composition and ratio of every cluster. The replays reproduce the published labels cell by cell. REFCON labeled no cell aneuploid in 15 of the 94 atlas tumors.

*Tumor cells too few to form a cluster.* In two test patients, no cell was labeled aneuploid. Wang patient s0115 has 171 tumor cells among 37,727 (0.45%). Their per-cell ratio is higher than that of the normal cells (median 1.10 against 0.88, AUROC 0.87), and they share predicted copy-number changes. They are spread over 5 of the 7 clusters and make up at most 2.4% of any cluster, so every cluster stays below $\tau_1$ (0.79 to 0.97). SCEVAN and CopyKAT reached an aneuploid F1 of at most 0.03 on these two patients.

*Normal cells clustered with tumor cells.* In the pancreatic biopsy PANFR0526, 7 of the 12 immune cells fall in two clusters of malignant cells. Over the atlas, 249 of the 1,137 immune cells REFCON labeled aneuploid (22%, in 33 tumors) lie in clusters that are at least half malignant.

*Normal clusters above $\tau_1$.* In Wang patient s0920, one cluster of 2,556 cells, 79% of them fibroblasts and 1.3% tumor cells, has a ratio of 1.138. It holds 2,522 of the 2,862 normal cells REFCON labeled aneuploid in the 17 test patients. Only 5 of the sample's 1,972 confident normals are fibroblasts. REFCON's per-cell dispersion rises with library size, and the fibroblasts are more deeply sequenced than the immune cells (median 3,243 against 875 UMI). Their dispersion is 1.31 times that of the immune cells, and 1.08 to 1.17 times at matched depth. When the liver metastasis was run alone (Supplementary Figure 4), fibroblasts formed a cluster just above $\tau_1$ in s1231 (ratio 1.071) and in s0115 (1.055). In these runs, REFCON labeled 2,357 and 1,135 normal cells aneuploid, 2,301 and 1,021 of them fibroblasts. In the whole-sample runs, 3,489 of these 3,492 cells were labeled diploid (Supplementary Figure 5). Over the atlas, 825 of the 1,137 immune cells REFCON labeled aneuploid (73%, in 15 tumors) lie in clusters with fewer than 5% malignant cells, 743 of them in clusters with a ratio below 1.10. The four such clusters made mostly of immune cells have ratios of 1.069 to 1.228. The largest is a dendritic-cell cluster in PANFR0557 (479 cells, ratio 1.078), and two are B-cell clusters, in SMC25 and SMC08. Within a tumor and cell type, the dispersion of immune cells rises with library size (median Spearman correlation 0.57).

*Malignant clusters below $\tau_1$.* In five atlas tumors, only REFCON had a malignant recall below 0.5. In four of them, the malignant cells form clusters below $\tau_1$: melanoma p21 (1.027), ovarian P7 (602 of its 802 malignant cells at 1.015, while a second malignant cluster at 1.375 was labeled aneuploid), glioblastoma 114 (0.928) and clear cell renal SI_22368 (0.750 and 0.913). Melanoma p26 has no confident normals, so the sample-level rule applied and labeled every cell diploid. SCEVAN and CopyKAT found most malignant cells of these five tumors (recall 0.68 to 0.99).

Labeling fails mainly for clusters near $\tau_1$ and for small tumor populations. Labels of such clusters are best checked against marker genes or cell-type annotations.

**Supplementary Note 3. Per-cell copy number of the DNTR-seq cell lines.** For the two DNTR-seq cohorts (HCT116 and A375; Zachariadis et al. [11], GSE144296), per-cell DNA reads are aligned to GRCh38 with BWA-MEM [52], processed with samtools [53] (mate-score fixing, coordinate sorting, and duplicate marking), and passed through AneuFinder v1.34 [54] (1 Mb bins, its e-divisive segmentation option, paired-end, hg38). Per-cell bin copy number is exported and projected to gene level using the per-gene genomic coordinates. DNA cells are matched to RNA cells by their shared plate-and-well identifier, which differs only by a DNA- versus RNA-modality suffix. Of the 1,656 HCT116 RNA cells, 1,468 have paired DNTR-seq DNA; the remaining 188 are Smart-seq2-only and are excluded from per-cell evaluation, although the expression-based methods were run on all 1,656 cells. Of the 1,468 paired cells, 1,467 have an evaluable per-cell profile (one lacks segmentation coverage) and constitute the per-cell HCT116 benchmark set. The A375 cohort uses the same pipeline and parameters; 64 of its 96 cells pass our per-cell filter on the AneuFinder output (finite per-cell segmentation, $\geq$ 50% bin coverage) and carry the per-cell ground truth used in Section 2.2, while the remaining 32 are excluded from per-cell metric computation, although every method was run on all 96 cells.

**Supplementary Note 4. External-method configurations, pseudo-reference injection and gene coverage.** We compare REFCON to five scRNA-seq CN callers. Each expression-based method has one call, used unchanged in every analysis (no per-cohort tuning). Only the cells given to a run change between analyses (one cell line, a pool of lines, a whole sample), together with the reference cells where stated, and CopyVAE's clone number:

- SCEVAN [19] 1.0.3: `pipelineCNA` with `SUBCLONES = FALSE` and `ClonalCN = FALSE`, otherwise its defaults. In reference-included runs, the reference cells are passed as `norm_cell`. We read its per-gene copy number matrix and its per-cell tumor or normal label.
- CopyKAT [18] 1.1.0 with its defaults (`cell.line = no`, `ngene.chr = 5`, `win.size = 25`, `KS.cut = 0.1`, Euclidean distance, genome hg20). Reference cells are passed as `norm.cell.names` in reference-included runs. We read its gene-level copy number file and its per-cell label. We fixed one bug in CopyKAT and relaxed one filter. Each change follows a fixed rule and was applied to every run that failed in the same way. First, to find its diploid baseline, CopyKAT clusters the cells into a few groups. When one group holds a single cell, which one outlying cell can cause, selecting that group's columns returns a vector instead of a one-column matrix. The run then stops with the error "dim(X) must have a positive length". No CopyKAT setting avoids this, its cell-line mode included. We kept these subsets as matrices (`drop = FALSE` in R) at the seven places in the code where a one-cell group can occur (two in `baseline.norm.cl`, one in `CNA.MCMC` and four in `copykat`). The fix changes only code that would otherwise stop, so a run that completes gives the same output with or without it. The failed runs were repeated with the same call and the fixed code. This applied to the 15 MIX-seq lines that failed when run alone (Section 2.2) and to the runs of the clustering analyses (Section 2.3) that failed the same way. Second, CopyKAT removes every cell with fewer than `ngene.chr` expressed genes on some chromosome. On the melanoma cohort this removes every Smart-seq2 cell, and the run stops with "all cells are filtered". These runs were repeated with `ngene.chr = 1` and the fixed code. One melanoma patient gave no output even then, and its cells count as unlabeled. Third, CopyKAT's own gene annotation places a few genes on

chromosome X or Y that lie on autosomes in GRCh38 (for example *SNORA70* and *DDX11L16*). Its per-cell chromosome check then removed every cell of some inputs, so the genes its annotation places on X or Y were removed from its input in every run.

- CopyVAE [20] with a maximum copy number of 6 (its default is 15), 25-gene bins, 200 epochs (both its defaults) and no clamping of a normal clone (an option of our wrapper), with a fixed random seed. Its clone number is the number of populations in the input: one for one cell line or one tumor's malignant cells, the number of lines in a pool, and one more when normal or reference cells are included. It has no reference argument. We read its copy number per bin, placed on genes with its own bin-to-gene map.
- inferCNV [14, 17], run as the Python implementation inferCNVpy 0.6.1 on total-count-normalized, log-transformed expression with its defaults (window of 100 genes, step 10, sex chromosomes excluded). In reference-included runs, the reference cells are passed through its reference key. With `calculate_gene_values = True`, we read its per-gene values.
- Numbat [39] 1.5.2, run on HCT116 and A375 only. Numbat adds population-phased B-allele frequencies to expression, so it needs each cohort's raw reads, aligned and genotyped. Reprocessing the raw reads of every cohort was impractical, and the astrocytoma's raw RNA reads are not public. Its per-cell results are in Supplementary Table 4. Each cell's RNA reads (GSE144296) were aligned with STAR 2.7.11b to GRCh38 (GENCODE v46). Allele counts were obtained with cellsnp-lite 1.2.3 and phased with Eagle 2.4.1 against the 1000 Genomes GRCh38 reference panel. Numbat was run with the settings its user guide gives for non-UMI data and for high-purity samples without normal cells: the built-in HCA expression reference, `gamma = 5` (non-UMI data), `call_clonal_loh = TRUE` and the default `t = 1e-5`. It received RNA only. With these settings Numbat stopped early on both lines, and each run was completed by a fixed rule. On HCT116 its entropy filter removed the only copy-number segment after the first iteration, so HCT116 was run with `max_entropy = 0.8`, the value the Numbat documentation gives for sparse allele coverage (spatial data). On A375 the final clone step failed on a three-cell clone. Every iteration of Numbat first drops clones of at most 50 cells (`min_cells`), and we applied the same filter to that final step. It runs after the per-cell posteriors are written and does not change them. On HCT116, Numbat received the 1,468 cells with paired DNA. Its per-gene output is the posterior expected copy-number ratio of the segment holding the gene (1 outside called segments).

Every expression-based method receives the same input as REFCON: raw integer counts in genomic order, and for the melanoma cohort [17] (Smart-seq2 [56]) the released TPM-derived integer matrix. Plotting steps are switched off, as they only draw. Each output is read on the method's own scale, and genes a method does not call stay missing until scoring (Supplementary Note 5). Reference-free and reference-included runs use the same call. A reference-included run adds the reference cells to the input and names them as the reference. Reference cells are given only in the reference-included copy-number benchmarks (HCT116 and A375 with the pseudo-references below, and the astrocytoma with its 450 non-malignant cells) and in one organoid check (Supplementary Note 1). In every other analysis, including all classification and clustering runs and the organoid comparisons, each method works on the sample alone and finds its own baseline cells. For classification, SCEVAN's tumor and normal labels and CopyKAT's aneuploid and diploid labels are read as aneuploid and diploid. CopyKAT's low-confidence labels count as the class they name. Cells that SCEVAN filters, cells that CopyKAT leaves undefined and cells a method does not return are unlabeled.

*Pseudo-reference injection.* Pseudo-reference injection follows Schmid et al. [16], who gave each cell line an external reference of the same tissue and technology. For each cell line cohort, up to 200 lineage-matched normal cells are subsampled from a public reference dataset and concatenated to the tumor expression matrix. Each expression-based external method is then run on the combined matrix, with the injected cells named as its reference (above; CopyVAE receives them as cells). The reference cohorts used in Section 2.2 are colonic epithelial cells from Kinchen et al. [29] for HCT116 (96 libraries) and normal epidermal cells from Belote et al. [30] for A375 (200 cells, 129 of them annotated melanocytes). All 96 Kinchen libraries are used, and the 200 epidermal cells are drawn at random from all Belote cells with a fixed seed.

*Gene coverage.* In the copy number benchmark, genes a method does not call are filled with the cell's median prediction (Supplementary Note 5). To confirm that REFCON's advantage is not an artifact of this convention, we recomputed each pairwise comparison on the genes the competitor itself calls, with no filling. REFCON's per-cell correlation with the paired single-cell DNA remained higher than every competitor across all three cohorts (HCT116, A375, scONE) and all four expression-based callers (12 of 12 comparisons; Pearson advantage +0.03 to +0.57). The same held for the per cell line comparison against bulk copy number on the esophageal and mixed-tissue lines (8 of 8; advantage +0.21 to +0.59). On HCT116, this test discards close to half of the genome for CopyKAT and SCEVAN, yet still favors REFCON.

**Supplementary Note 5. Benchmark protocol.** Supplementary Note 4 gives the configuration of every compared method, including Numbat, and the pseudo-reference injection.

Each method's predictions are placed on the dataset's own genes, and missing entries are filled with the per-cell median of finite predictions before metric computation. This genome-wide median filling is required because CopyKAT covers 47% of evaluable genes on HCT116, SCEVAN 50%, while REFCON and inferCNV cover above 99% and CopyVAE 95%; without filling, low-coverage methods would be evaluated on a smaller gene denominator than REFCON, biasing per-cell aggregates. Supplementary Note 4 repeats every comparison without filling.

The five expression-based methods compared in this work emit predictions in different numerical spaces: REFCON outputs CN ratios renormalized to a per-cell mean of one, CopyKAT and SCEVAN emit smoothed log-scale values of transformed expression relative to their baseline cells, CopyVAE emits integer copy number per bin, and inferCNV emits smoothed log expression relative to its reference. Applying a single fixed cutoff to every method's output would either privilege a method whose scale matches the threshold or penalize a method whose scale does not, and would conflate prediction quality with the choice of an arbitrary cutoff. We therefore apply the loss and gain thresholds to the ground truth only and use metrics that need no cutoff on the predictions (Methods, Section 4.9). Loss AUROC and AUPRC use the negated prediction as the score. Because cell line ploidy is often near-triploid, an absolute gain threshold labels a large fraction of the genome as gain. Supplementary Table 15 therefore repeats the AUROC with classes relative to each cell's ploidy.

Unsupervised cell line clustering (Section 2.3) uses a benchmark pipeline distinct from the classification clustering of Section 4.6: each method's predicted copy-number matrix is projected onto its top 20 principal components, standardized, and partitioned by three algorithms, $k$-means (10 initializations), Ward hierarchical clustering, and a tied-covariance Gaussian mixture (one initialization), applied identically to every method at the known number of populations with random seed 42, and scored by the adjusted Rand index against the known cell line identities. Supplying the true population count to every method isolates copy-number-profile quality from cluster-count model selection. The reference-based methods are run both per cell line and pooled; REFCON is run on each line alone. Runs of single lines are joined on the genes present in every line's input (7,767 to 11,554 genes per cohort) before the projection, and pooled runs use their input's genes. For display (Figure 4B and Supplementary Figure 2), the standardized components are embedded with UMAP [32] (30 neighbors, minimum distance 0.3, random seed 42). For the composition analyses (Figure 4C,D), SCEVAN, CopyKAT, CopyVAE and inferCNV are run on the seven esophageal lines at cohort sizes of one to seven lines, with six random line compositions per intermediate size, and on NCIH1650 paired with each of the other 20 lines of the MIX-seq lung panel in turn (all cells of both lines; CopyVAE with one clone per line).

### Supplementary Data

Each Supplementary Data file is provided separately.

**Supplementary Data 1. Cell line and healthy-cell roster.** Two tables. The cell line entries of the training, held-out CCLE validation, test and MIX-seq clustering splits, with CCLE name, DepMap identifier, tissue of origin and cell count. The 142 training rows (60,179 cells) correspond to 140 DepMap models: MCF7 (ACH-000019) appears in its standalone dataset and in the CCLE panel, and COLO320 (ACH-000202) contributes two sublines, with 9,000 training cells subsampled from the deposited cells. COLO680N, TE6 and NCI-H838 are test lines that are also among the MIX-seq lines. CellBench's fifth line, NCI-H2228, is a training line and is not used for testing. The healthy Tabula Sapiens cells by split, organ and donor, with the CZ CELLxGENE Discover dataset, version and collection identifiers.

**Supplementary Data 2. Atlas tumor samples.** The 94 tumor samples of the Curated Cancer Cell Atlas with study, cancer type, original accession and cell counts, the share of immune cells each method labels aneuploid (or tumor) or leaves unlabeled, and each method's per-tumor aneuploid F1, recall and precision, diploid recall and share of cells labeled (Supplementary Table 7).

**Supplementary Data 3. Per-patient classification.** Each method's classification of every held-out tumor patient and control sample, and of the colorectal patients run per tissue (column `run`): cells, malignant cells, true and false positives and negatives, malignant and non-malignant cells left unlabeled, call rate, and aneuploid and diploid precision, recall and F1.
